# Human Milk Oligosaccharide Utilization in Intestinal Bifidobacteria is Governed by a Global Transcriptional Regulator NagR

**DOI:** 10.1101/2022.04.06.487429

**Authors:** Aleksandr A. Arzamasov, Aruto Nakajima, Mikiyasu Sakanaka, Miriam N. Ojima, Takane Katayama, Dmitry A. Rodionov, Andrei L. Osterman

**Author notes:** Adress correspondence to Aleksandr A. Arzamasov, and Andrei L. Osterman,.

## Abstract

*Bifidobacterium longum* subsp. *infantis* (*B. infantis*) is a prevalent beneficial bacterium that colonizes the human neonatal gut and is uniquely adapted to efficiently use human milk oligosaccharides (HMOs) as a carbon and energy source. Multiple studies have focused on characterizing the elements of HMO utilization machinery in *B. infantis*; however, the regulatory mechanisms governing the expression of these catabolic pathways remain poorly understood. A bioinformatic regulon reconstruction approach used in this study implicated NagR, a transcription factor from the ROK family, as a negative global regulator of genomic loci encoding lacto-*N*-biose/galacto-*N*-biose (LNB/GNB), lacto-*N*-tetraose (LNT), and lacto-*N*-neotetraose (LNnT) utilization pathways in *B. infantis*. This conjecture was corroborated by transcriptome profiling upon *nagR* genetic inactivation and experimental assessment of binding of recombinant NagR to predicted DNA operators. The latter approach also implicated *N*-acetylglucosamine (GlcNAc), a universal intermediate of LNT and LNnT catabolism, and its phosphorylated derivatives as plausible NagR transcriptional effectors. Reconstruction of NagR regulons in various *Bifidobacterium* lineages revealed multiple regulon expansion events, suggesting evolution from a local regulator of GlcNAc catabolism in ancestral bifidobacteria to a global regulator controlling foraging of mixtures of GlcNAc-containing host-derived glycans in mammalian gut-colonizing *B. infantis* and *Bifidobacterium bifidum*.

**Importance:** The predominance of bifidobacteria in the gut of breastfed infants is attributed to the ability of these bacteria to utilize human milk oligosaccharides (HMOs). Thus, individual HMOs such as lacto-*N*-tetraose (LNT) and lacto-*N*-neotetraose (LNnT) are considered promising prebiotics that would stimulate the growth of bifidobacteria and confer multiple health benefits to preterm and malnourished children suffering from impaired (stunted) gut microbiota development. However, the rational selection of HMO-based prebiotics is hampered by the incomplete knowledge of regulatory mechanisms governing HMO utilization in target bifidobacteria. This study describes NagR-mediated transcriptional regulation of LNT and LNnT utilization in *Bifidobacterium longum* subsp. *infantis*. The elucidated regulatory network appears optimally adapted to simultaneous utilization of multiple HMOs, providing a rationale to add HMO mixtures (rather than individual components) into infant formulas. The study also provides insights into the evolutionary trajectories of complex regulatory networks controlling carbohydrate metabolism in bifidobacteria.

## Introduction

Bifidobacteria are Gram-positive, anaerobic, saccharolytic microorganisms that colonize the digestive tracts of humans and various animals (1). Certain *Bifidobacterium* species, namely *Bifidobacterium longum* subsp. *infantis* (*B. infantis*), *Bifidobacterium bifidum*, and *Bifidobacterium breve*, often predominate the human neonatal gut microbiota (GM) during breastfeeding (2–5), and their predominance is directly linked with the healthy development of an infant (6, 7). Among health-promoting effects attributed to bifidobacteria inhabiting the neonatal gut are protection from pathogen colonization (8–10) and modulation of the immune system towards the tolerance to commensal microorganisms (5, 11).

Decreased *Bifidobacterium* abundance is characteristic of stunted GMs observed in preterm infants (14) and children suffering from severe acute malnutrition (73). Therapeutic approaches aimed at restoring the bifidobacterial population in these affected groups include administering exogenous *Bifidobacterium* species (e.g., *B. infantis*) as probiotics or/and food formulas containing prebiotics that would selectively stimulate the growth of endogenous bifidobacteria in the gut and thus confer beneficial properties to the infant (14–16, 73). Since the prevalence of bifidobacteria in the neonatal gut is often attributed to their ability to selectively utilize dietary human milk oligosaccharides (HMOs) (2, 17–21), these milk glycans are considered “natural” prebiotics and added to infant formulas.

HMOs are the third most abundant (5 to 20 g/L) component of human milk after lactose and lipids and are not assimilated by the infant (22–24). HMO building blocks include glucose, galactose, *N*-acetylglucosamine (GlcNAc), L-fucose, and *N*-acetylneuraminic (sialic) acid; these units form more than a hundred linear or branched oligosaccharide structures (22, 25, 26). Most HMOs are composed of a lactose core (Galβ1-4Glc) conjugated to one or multiple lacto-*N*-biose (LNB, Galβ1-3GlcNAc) or *N*-acetyllactosamine units (Galβ1-4GlcNAc) (22, 23, 26). The resulting HMO structures are denoted as type I and type II chains, respectively, with lacto-*N*-tetraose (LNT) and lacto-*N*-neotetraose (LNnT) as archetypes.

Previous studies have revealed substantial variation in HMO utilization strategies and capabilities within the *Bifidobacterium* genus (19, 27). For example, *B. bifidum* uses a set of membrane-attached glycoside hydrolases (GHs) that extracellularly degrade HMOs to di- and monosaccharides (28–32) and then imports and catabolizes liberated LNB and lactose (19). In contrast, *B. infantis* and *B. breve* import HMOs using ATP-binding cassette (ABC) transporters and then degrade uptaken oligosaccharides to monosaccharides intracellularly using a repertoire of exo-acting GHs (17, 20, 27).

*B. infantis*, which is widely used in probiotic and synbiotic formulations (8, 15, 16, 73), possesses several unique genomic clusters (e.g., HMO cluster I or H1) that encode the most elaborate HMO uptake (33, 34) and intracellular degradation machinery (35–38) among bifidobacteria (**Fig. 1A-B**). While the molecular mechanisms of HMO utilization in *B. infantis* have received considerable attention, the regulatory mechanisms governing this catabolic process remain poorly understood. Previous studies demonstrated that LNT and LNnT induce a profound and surprisingly similar transcriptomic response in *B. infantis* ATCC 15697, with multiple genomic clusters (*nag*, *lnp*, H1; **Fig. 1B**) being upregulated, pointing to a possible global regulatory mechanism(s) (39, 40). Uncovering this yet unknown mechanism in addition to fundamental importance has a potential translational value for the rational selection of individual HMOs for prebiotic and synbiotic formulations.

**FIG 1.**
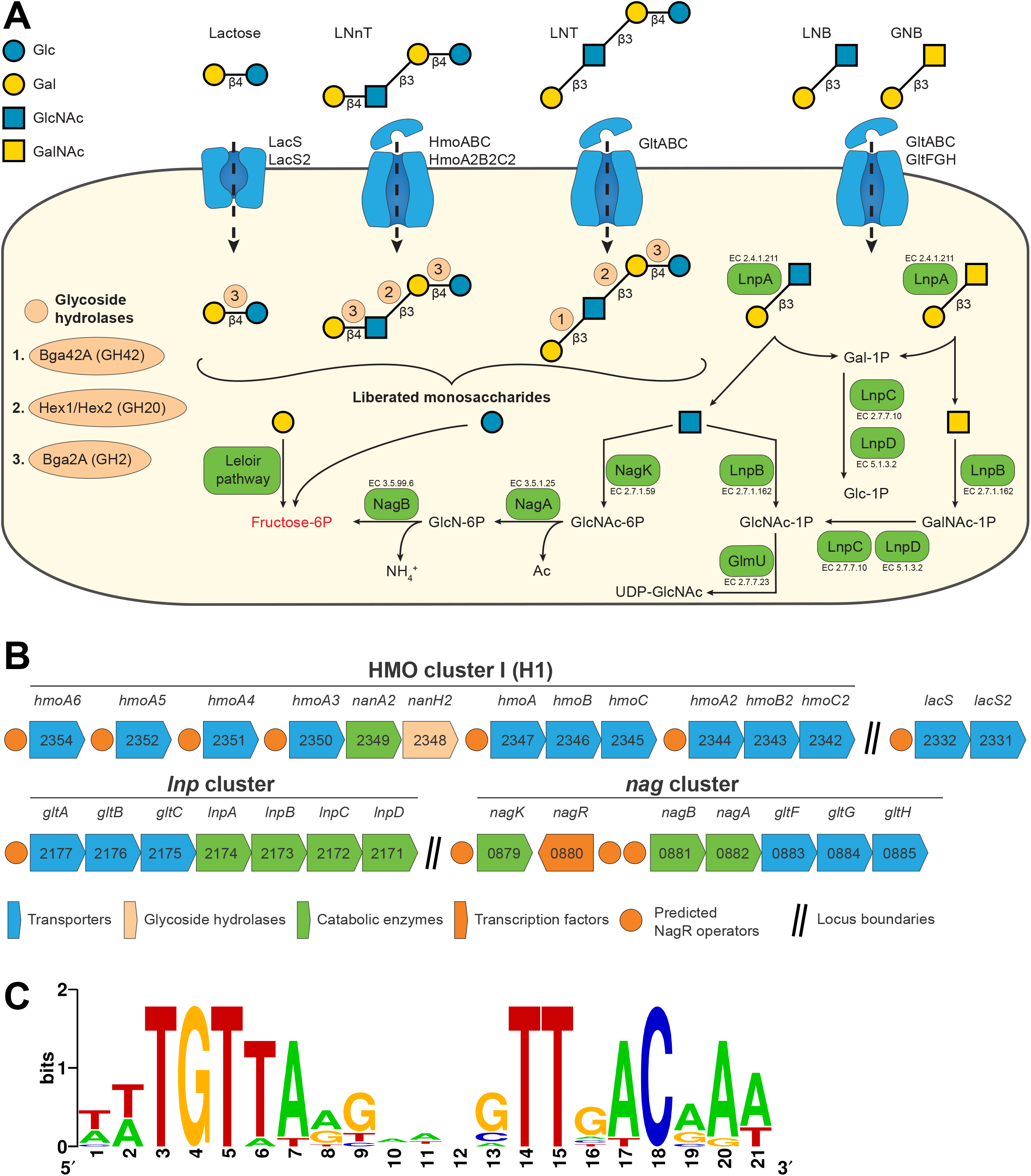
Reconstructed NagR regulon in *B. infantis* ATCC 15697. (A) Schematic representation of LNT, LNnT, and LNB/GNB utilization pathways in *B. infantis* ATCC 15697. *Step 1*: HMOs and their constituents are transported into the cell by various transport systems. *Step 2*: once inside the cell, HMOs are degraded from the non-reducing end by a coordinated action of exo-acting GHs. Breakdown of glycosidic bonds by specific GHs is indicated in light orange circles. *Step 3*: released monosaccharides are converted to fructose-6P and enter the bifid shunt. (B) Genomic loci constituting the reconstructed NagR regulon in *B. infantis* ATCC 15697. Numbers represent locus tags in the Blon_XXXX format (GenBank accession no. CP001095.1). (C) NagR-binding motif in *B. infantis* ATCC 15697 based on 11 predicted operator sequences.

Comparative genomic analysis across multiple related genomes is a powerful approach for reconstructing transcriptional regulatory networks (regulons) controlling carbohydrate utilization (41–43). Our earlier bioinformatic analysis conducted on a limited set of *Bifidobacterium* genomes implicated NagR, a transcription factor (TF) from the ROK family, as a regulator of *nag* and *lnp* clusters (**Fig. 1B**) encoding GlcNAc and lacto-*N*-biose/galacto-*N*-biose (LNB/GNB) catabolic pathways, respectively, in *B. bifidum, B. breve, B. infantis*, and *Bifidobacterium longum* subsp. *longum* (*B. longum*) (42). James et al. further experimentally confirmed this prediction in *B. breve* UCC2003 (44). However, it was unclear whether and how this knowledge translated to the global regulation of the extensive HMO utilization machinery in *B. infantis*.

Here we reconstructed NagR regulons in a substantially larger collection of *Bifidobacteriaceae* genomes focusing on HMO-utilizing species. This analysis revealed multiple putative NagR-binding sites (operators) in the *B. infantis* ATCC 15697 genome, suggesting the role of NagR as a global negative regulator of LNB/GNB, LNT, and LNnT utilization in this bacterium. This conjecture was confirmed by transcriptome profiling of the *nagR* knockout mutant and by direct assessment of the binding of recombinant NagR to its predicted operators. The inferred NagR regulon structure suggests that at the transcriptional level, *B. infantis* is adapted to simultaneous utilization of multiple HMOs, providing guidelines for using rationally formulated mixtures rather than individual oligosaccharides as prebiotics. The reconstructed NagR regulons also provided insights into the evolution of complex regulatory networks controlling carbohydrate metabolism in bifidobacteria shaped by the ecological niche of these commensal microorganisms.

## Results

### Genomic reconstruction reveals the complexity of the NagR regulon in *B. infantis*

We used a Position Weight Matrix (PWM)-based approach to reconstruct the NagR regulon in *B. infantis* ATCC 15697 and identified 11 potential NagR-binding sites (operators) in promoter regions of genes/operons encoding components of HMO utilization pathways including six previously unknown NagR operators within in the HMO cluster I (H1) (**Fig. 1A-B and Table S3**). Among the new putative NagR regulon members were genes encoding (i) LNnT (type II HMOs) ABC transporters (HmoABC, HmoA2B2C2) (33), (ii) substrate-binding components of ABC transporters possibly involved in HMO uptake (HmoA3, HmoA4, HmoA5, HmoA6) (33), (iii) *N*-acetylneuraminate lyase NanA2 and exo-α-sialidase NanH2 (GH33) (35), (iv) lactose permeases (LacS, LacS2) (**Table 1** and **Fig. 1A**). Overall, the reconstructed regulon contained 29 genes. The inferred NagR-binding motif had a palindrome structure (**Fig. 1C**), a common feature of transcriptional regulators from the ROK family (45).

**TABLE 1.**
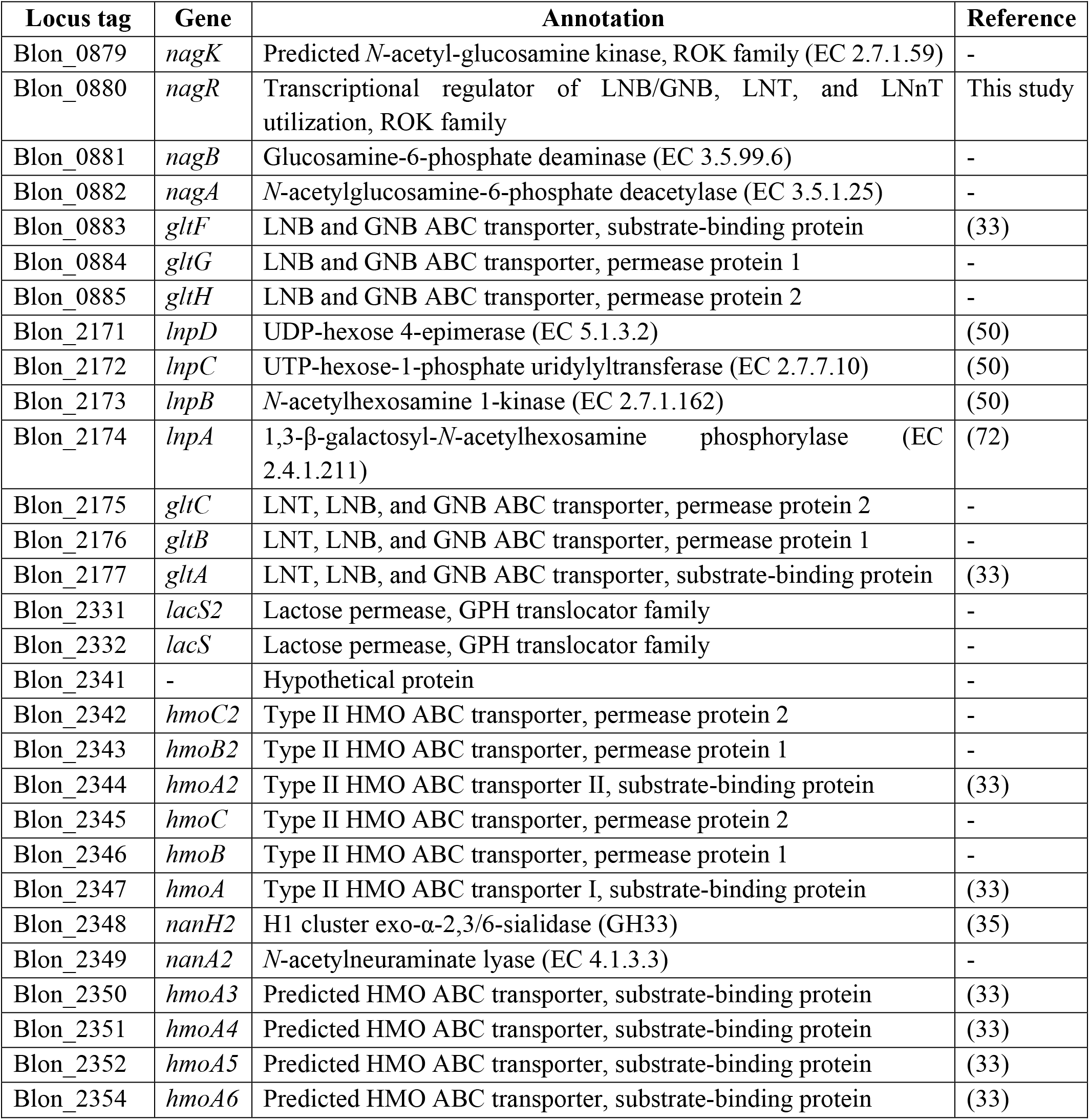
Composition of the reconstructed NagR regulon in *B. infantis* ATCC 15697.

ROK-family TFs can function as transcriptional activators or repressors (45). To infer the possible mode of action of NagR, we analyzed the position of predicted NagR operators relative to −35 and −10 promoter elements recognized by bacterial RNA polymerase holoenzyme. Ten out of 11 predicted NagR operators overlapped either with −10 or −35 sequences (**Fig. S1**), suggesting that NagR was a potential transcriptional repressor. Therefore, based on the genomic reconstruction, we hypothesized that NagR was a global negative regulator of genomic loci involved in LNB/GNB, LNT, and LNnT utilization in *B. infantis* ATCC 15697.

### Engineered *nagR* insertional mutant displays comparable yet distinct physiological properties compared to the parental wild-type strain

To experimentally study the proposed regulatory role of NagR, we generated a *nagR* knockout mutant of *B. infantis* ATCC 15697 (*nagR*-KO) by insertional mutagenesis and verified the insertion position using genomic PCR (**Fig. S2A-B**). The *nagR-KO* mutant displayed a growth rate comparable to the wild-type (WT) strain in the medium supplemented with lactose (MRS-CS-Lac) but grew significantly slower (linear regression; P_adj_ < 0.05) in the medium supplemented with a mixture of HMOs purified from pooled breastmilk (MRS-CS-HMO) (**Fig. 2**). Unexpectedly, *nagR*-KO exhibited decreased growth in MRS-CS containing sucrose but not fructooligosaccharides (FOS) (**Fig. 2**).

**FIG 2.**
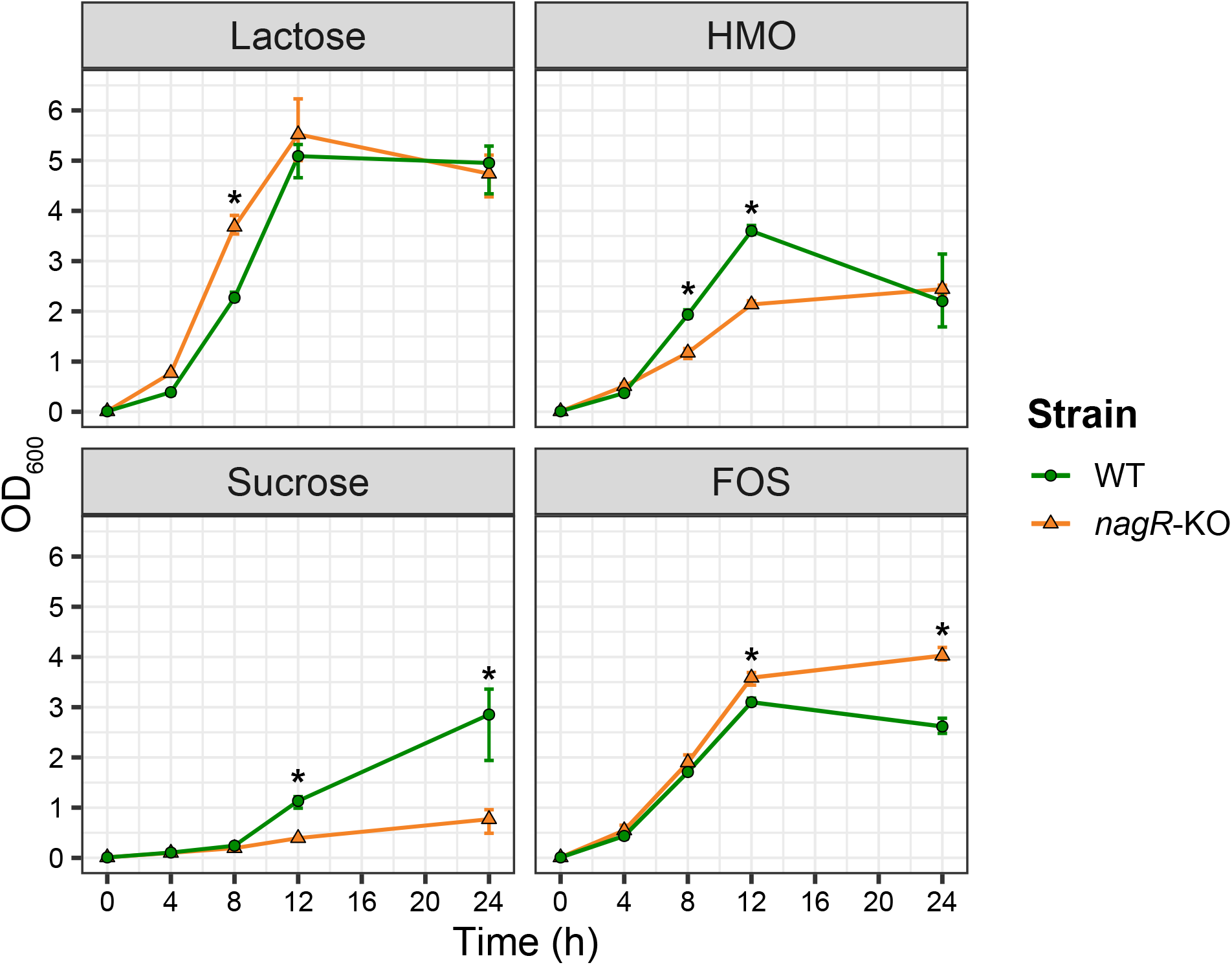
Growth curves of *B. infantis* ATCC 15697 WT and *nagR*-KO strains in MRS-CS supplemented with various carbon sources (1% w/v). Growth was monitored by measuring OD_600_ at specific time points. Data points represent the mean of three biological replicates. Error bars depict 95% confidence intervals for the mean obtained by bootstrapping. Timepoints where the OD_600_ values for WT and *nagR*-KO strains were significantly different (*, *P*_adj_<0.05) were identified using a linear regression. Bonferroni correction was used to adjust for multiple testing.

HMO consumption profiling of WT and *nagR*-KO strains grown in MRS-CS-HMO revealed that both strains completely salvaged lactose, LNT, LNnT, 2′- and 3-fucosyllactose (FL), and difucosyllactose (DFL) present in the HMO mixture (**Fig. S2C**). Notably, the *nagR*-KO mutant consumed LNT faster compared to WT and displayed significantly delayed consumption (linear regression; P_adj_ < 0.05) of large fucosylated HMOs, namely lacto-*N*-fucopentaoses (LNFP) I/II/III, and lacto-*N*-difucohexaoses (LNDFH) I/II (**Fig. S2C**). The mutant also expelled less fucose into the medium (**Fig. S2C**). Organic acid production profiling of supernatants of WT and *nagR*-KO grown in MRS-CS-HMO showed that both strains released similar quantities of acetic acid; however, the *nagR*-KO mutant produced significantly more formic and less lactic acid (linear regression; P_adj_ < 0.05) (**Fig. S2D**). These data demonstrated comparable yet distinct physiological properties of WT and *nagR*-KO strains, prompting a follow-up transcriptome profiling to infer genes differentially expressed in the mutant.

### Comparative transcriptomics corroborates the role of NagR as a global negative regulator of LNB/GNB, LNT, and LNnT utilization in *B. infantis*

We used RNA-seq to compare transcriptomes of *nagR*-KO and WT strains of *B. infantis* ATCC 15697 grown in MRS-CS supplemented with lactose or LNnT, yielding four experimental conditions (**Fig. 3A**). The choice of carbon sources was based on a previous study, where LNnT had induced many HMO utilization genes, and lactose had been used as a comparator (39). Principle component analysis (PCA) of TMM-normalized counts revealed that each experimental condition formed a distinct cluster (**Fig. 3B**). Specifically, PCA separated samples by carbon source (PC1, 38.8% of the total variance) and strain (PC2, 21.1% of the total variance).

**FIG 3.**
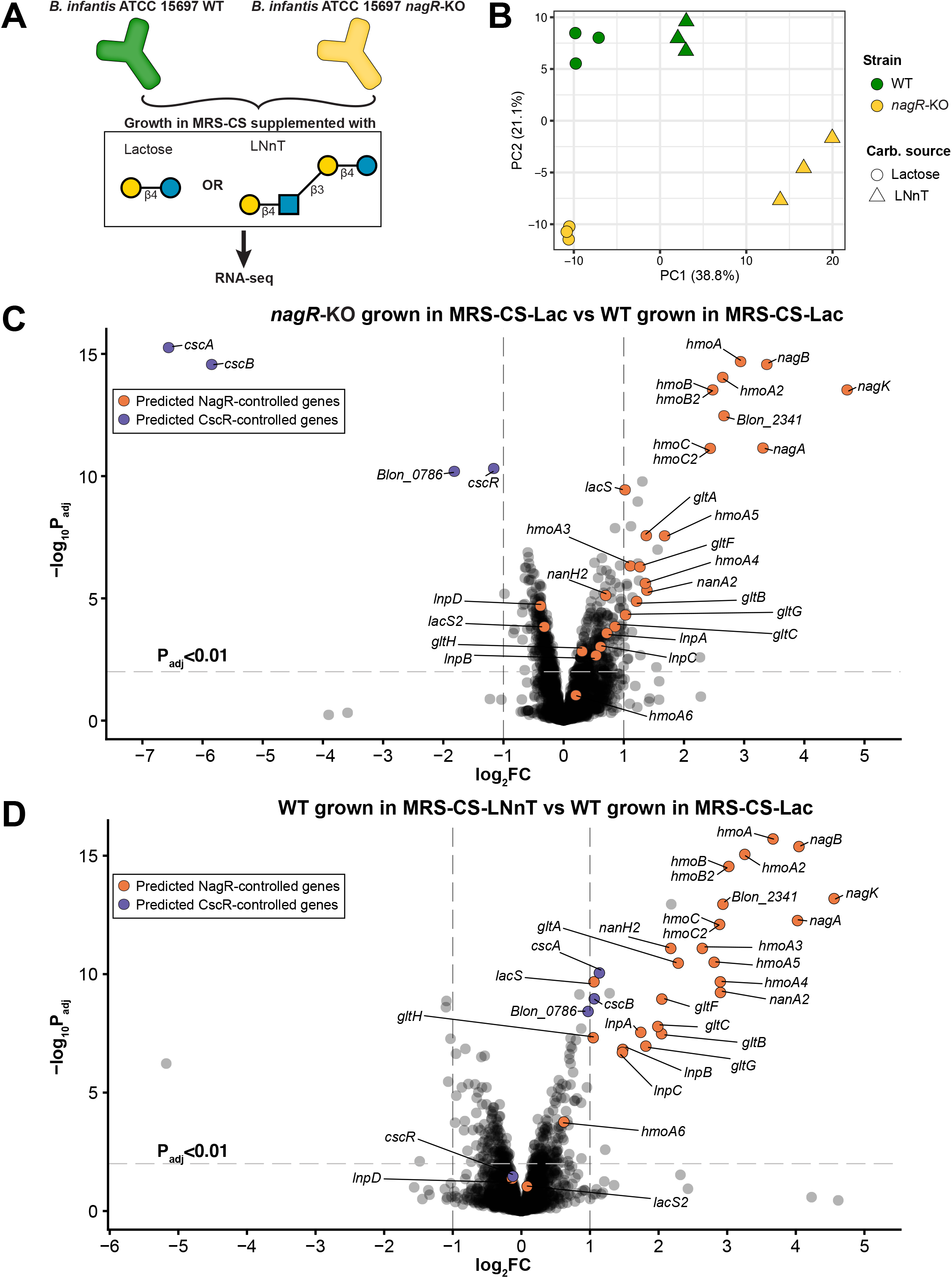
RNA-seq of WT and *nagR*-KO strains grown in MRS-CS supplemented with lactose and LNnT. (A) Schematic representation of the experimental design. (B) PCA of TMM-normalized count data. Each data point represents one sample. (C) and (D) Volcano plots depicting the log2FC of gene expression versus the –log_10_P_adj_. (C) compares *nagR*-KO and WT strains grown in MRS-CS-Lac, whereas (D) compares the WT strain grown in MRS-CS-LNnT and MRS-CS-Lac. Criteria for calling differentially expressed genes were the follows: P_adj_ < 0.01 and absolute FC > 2. Genes constituting the predicted NagR and CscR regulons are labeled and colored in orange and purple, respectively.

Linear modeling implemented in the *limma* framework (46) revealed significant upregulation (Fold change (FC) > 2 and P_adj_ < 0.01) of multiple *nag*, *lnp*, and H1 cluster genes in the *nagR*-KO strain grown in MRS-CS-Lac compared to WT grown in MRS-CS-Lac (**Table S2A** and **Fig. 3C**). Overall, 19 out of 29 genes constituting the predicted NagR regulon were upregulated. These results demonstrated that NagR, in line with the bioinformatic prediction, functioned as a transcriptional repressor of the *nag*, *lnp*, and H1 loci. In addition, we observed: (i) significant upregulation of *malEFG* encoding an ABC transport system for maltose (33) and (ii) significant downregulation (FC < −2 and P_adj_ < 0.01) of *cscA, cscB*, and *cscR* in the *nagR*-KO mutant (**Table S2A** and **Fig. 3C**). The latter genes are involved in sucrose uptake and catabolism (47) and are predicted to be controlled by a local regulator from the LacI family, CscR (42). Notably, downregulation of the *csc* genes was consistent with decreased growth of the *nagR*-KO mutant in the medium supplemented with sucrose (**Fig. 2**). No potential NagR operators were identified in the promoter regions of *malEFG* and *csc* cluster genes, suggesting that the effect of the *nagR* knockout on the expression of these genes was indirect.

RNA-seq of the WT strain revealed that 25 out of 29 genes constituting the predicted NagR regulon were upregulated during growth in MRS-CS-LNnT compared to MRS-CS-Lac (**Table S2B** and **Fig. 3D**). Interestingly, the expression of *nag*, *lnp*, and H1 cluster genes was higher in the WT strain cultured in the presence of LNnT than in the *nagR*-KO mutant cultured in MRS-CS-Lac (**Fig. S3**). These observations point to the presence of additional transcriptional activation mechanism(s) that may contribute to the upregulation of these genomic clusters during the growth of *B. infantis* in the medium containing LNnT. We also observed a significant upregulation of *malK* (**Table S2B** and **Fig. S3**) encoding a shared ATPase component that energizes most carbohydrate-specific ABC transporters in *B. infantis* (42). The upregulation of *malK* was likely tied with the enhanced energetic needs associated with growth in the presence of LNnT, specifically upregulation of genes encoding multiple ABC transport systems (GltFGH, GltABC, HmoABC, HmoA2B2C2, HmoA3-A5).

Overall, the obtained transcriptomic data were consistent with our bioinformatic prediction and demonstrated that NagR regulated LNB/GNB, LNT, and LNnT utilization in *B. infantis* by repressing genomic loci encoding (i) transporters of respective glycans, (ii) GlcNAc and LNB/GNB catabolic pathways.

### Interaction of NagR with predicted operators is dependent on GlcNAc and its phosphorylated derivatives

To test the interaction between NagR and its predicted operator sequences, we cloned the *nagR* gene from *B. infantis* ATCC 15697 and expressed the recombinant protein as a fusion with an N-terminal His-tag in *E. coli* BL21/DE3. Electrophoretic Mobility Shift Assay (EMSA) demonstrated that NagR specifically bound DNA fragments (probes) containing predicted NagR operators located upstream of *nagK, nagB, gltA, hmoA, hmoA2*, and *hmoA3* genes (**Fig. 4A-B, Fig. 5**, and **Table S1**). Titration with increasing concentrations of NagR revealed that probes *nagK*, *nagB_I, hmoA, hmoA2* had a high affinity for NagR (EC_50_: 13-23 nM), whereas probes *gltA, hmoA3* had a moderate one (EC_50_: 140-180 nM) (**Fig. 5**). Reactions with probes *nagB_II* and *hmoA6* did not manifest in robust shifts even at high (> 500 nM) protein concentrations (**Fig. S4A**). These results showed that NagR exhibited different affinities to its various operator sequences. Notably, we also observed a significant negative correlation (Pearson R = −0.84, p < 0.05) between calculated EC50 values for probes and expression FCs for respective genes in the RNA-seq experiment (*nagR*-KO vs. WT grown in MRS-CS-Lac) (**Fig. 3C and 4C**), indicating the more tightly NagR bound to an operator, the stronger was the observed upregulation of the respective gene in *nagR*-KO.

**FIG 4.**
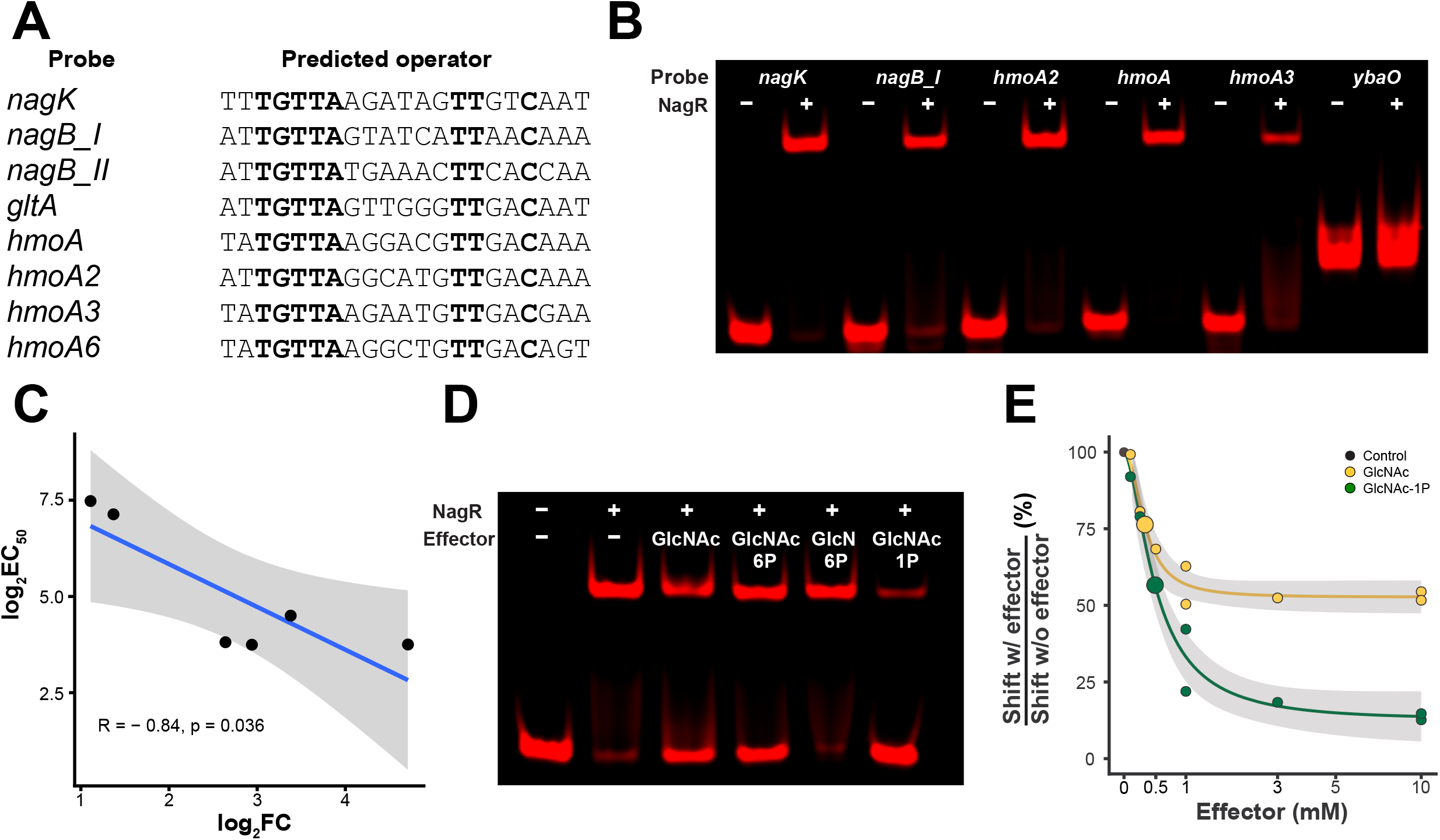
Interactions of recombinant NagR with predicted operators and screening of NagR effectors. (A) Predicted NagR operator sequences identified in the promoter regions of listed genes. Conserved nucleotides are in bold. The full sequences of probes used in EMSA experiments are given in Table S1. (B) An EMSA gel showing interactions of NagR with DNA probes containing predicted operators. NagR concentration was 25 nM, and probe concentrations were 1 nM. The *ybdO* probe was used as a control for non-specific binding. (C) Correlation between probe EC50 values determined via EMSA and expression fold-changes of cognate genes in the RNA-seq experiment (*nagR*-KO vs. WT grown in MRS-CS-Lac). (D) Effect of various GlcNAc metabolism intermediates (10 mM) on the interaction between NagR (25 nM) and the *hmoA* probe (1 nM). (E) EC50 values of selected NagR effector molecules. The y-axis depicts the ratio 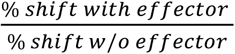. Calculated EC_50_ are shown as big circles. 25 nM of NagR and 1 nM of the *hmoA* probe were used.

**FIG 5.**
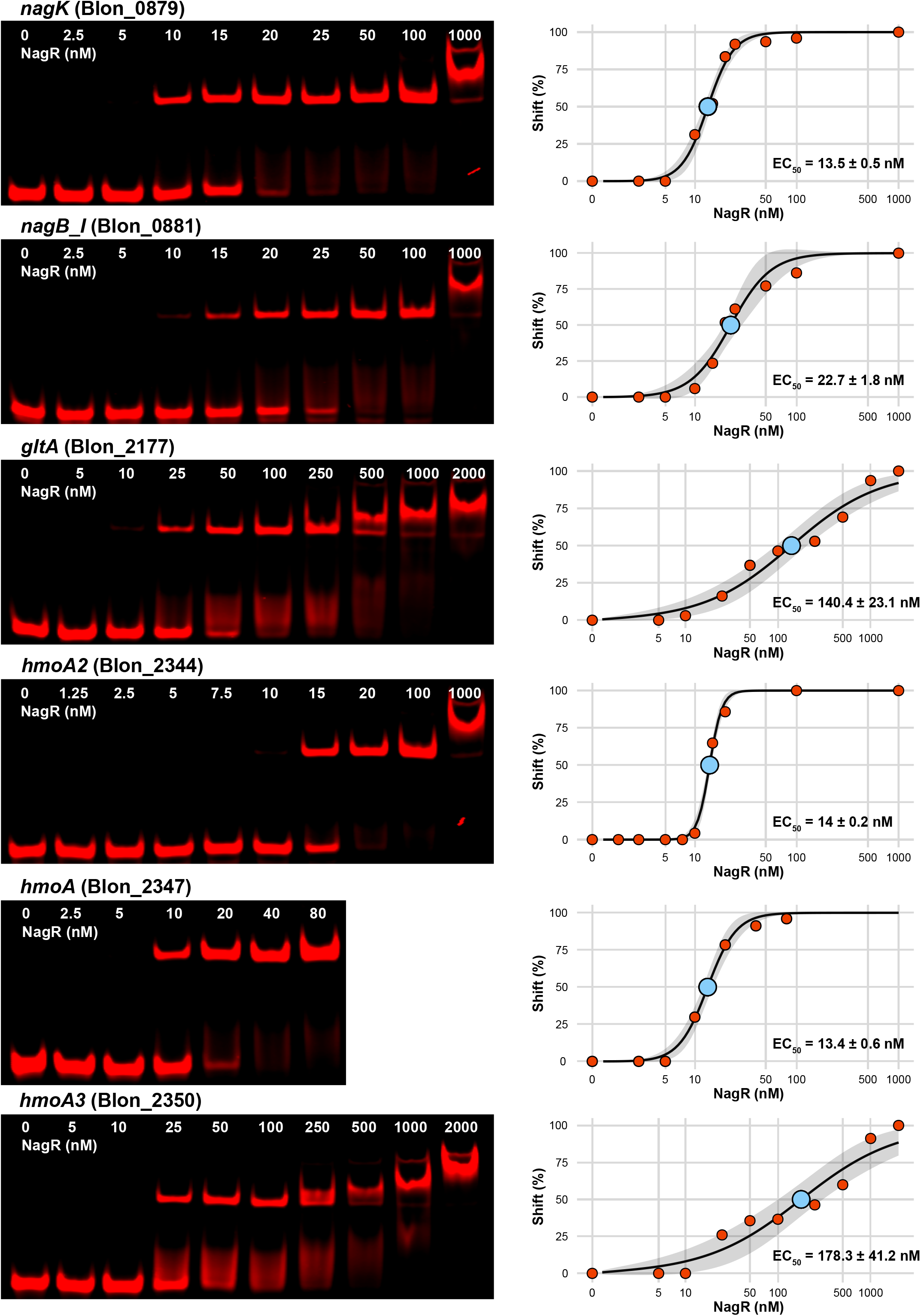
EMSA gels depicting titration of DNA probes (1 nM) containing predicted NagR operators with recombinant NagR. Gels were quantified, and the results were approximated by the 4PL equation. NagR concentration at which half of the probe is shifted (EC_50_) is shown. Grey shading depicts 95% confidence intervals. The x-axis is in the Log10 scale.

To identify potential NagR effector molecules, we used a probe with a high affinity to NagR (*hmoA*) and added various GlcNAc metabolism intermediates (GlcNAc, GlcNAc-6P, GlcN-6P, GlcNAc-1P) to the binding reaction. GlcNAc, GlcNAc-6P, and GlcNAc-1P, but not GlcN-6P disrupted the NagR-DNA complex at saturating (10 mM) concentration (**Fig. 4D**). Titration of the effectors revealed that GlcNAc-6P lost its complex disrupting effect at 1 mM, whereas GlcNAc and GlcNAc-1P lost their effect at 0.1 mM (**Fig. S4B**); the calculated effector EC50 values were 0.33 ± 0.05 mM for GlcNAc and 0.49 ± 0.06 mM for GlcNAc-1P (**Fig. 4E**).

Overall, the EMSA results demonstrated that (i) NagR bound its predicted operator sequences with various affinities and (ii) multiple GlcNAc metabolism intermediates disrupted NagR-DNA interactions *in vitro* and thus served as plausible NagR transcriptional effectors in *B. infantis*.

### NagR regulon was a subject of evolutionary expansion in *Bifidobacteriaceae*

We used the PWM-based approach to identify NagR binding motifs and reconstruct regulons in 25 genomes representing various phylogenetic lineages within the *Bifidobacteriaceae* family to trace the potential evolutionary history of this gene regulatory network. Overall, the size and composition of the reconstructed NagR regulons markedly varied among the studied genomes (**Fig. 6A and Table S3**). For instance, early diverged *Bifidobacterium* species isolated from insects and dairy products possessed concise NagR regulons comprised of a single genomic locus (*nag*) encoding GlcNAc and predicted *N*,*N*′-diacetylchitobiose catabolic pathways (see **Supplementary Results**). This observation suggests that in ancestral bifidobacteria, the role of NagR was likely confined to local regulation of the GlcNAc catabolic pathway.

**FIG 6.**
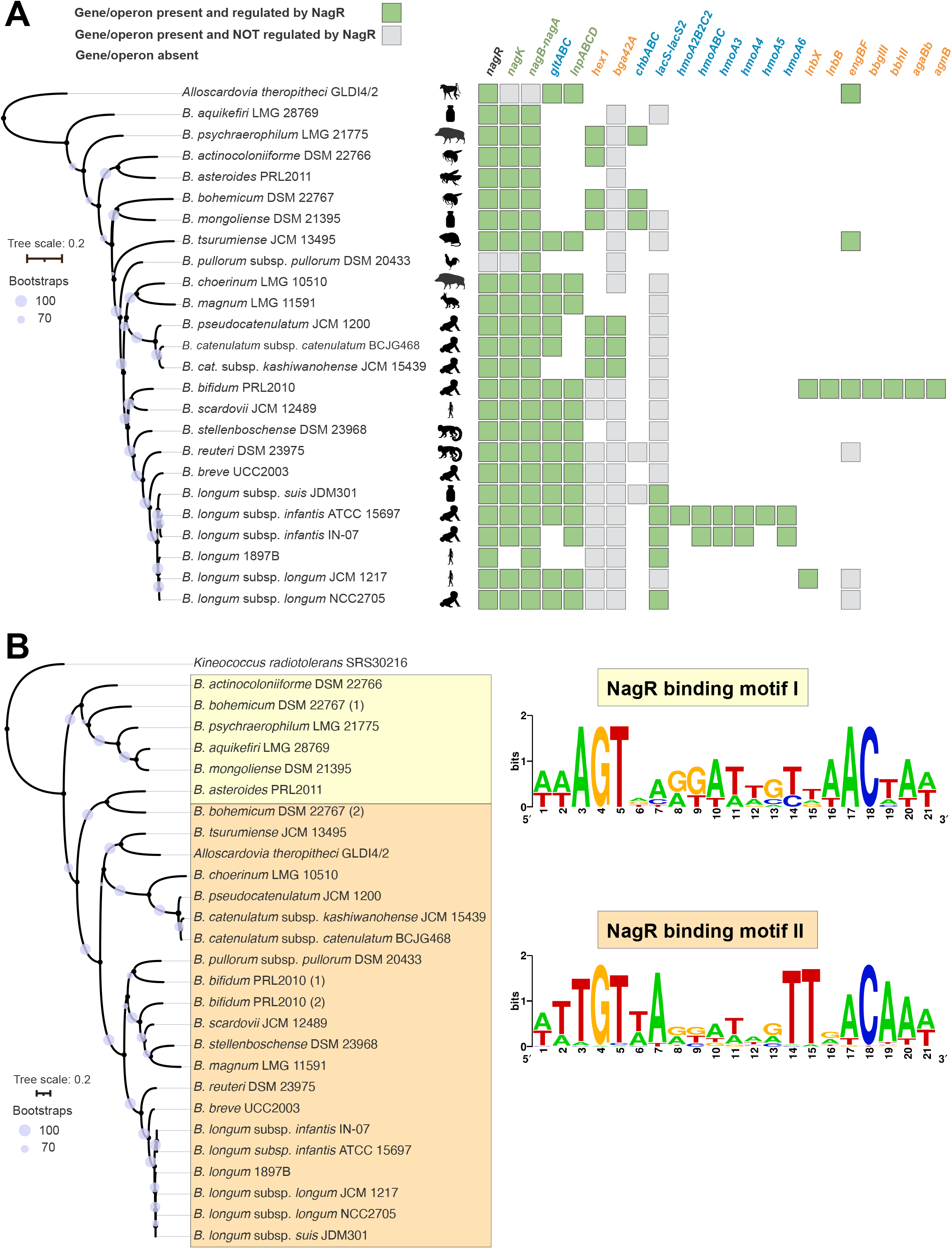
Evolution of the NagR regulon and binding motif within the *Bifidobacteriaceae* family. (A) NagR regulon composition in 25 *Bifidobacteriaceae* strains mapped on a species tree built based on the alignment of 247 core genes. Bootstrap values are shown as purple circles. Black symbols indicate strain isolation sources. Regulon members are colored according to their function: catabolic enzymes are green, GHs in orange, and transporters in blue. (B) NagR-binding motifs mapped on a tree of NagR proteins from 25 *Bifidobacteriaceae* strains. Bootstrap values are shown as purple circles. NagR paralogs in *B. bohemicum* and *B. bifidum* are denoted by numbers (1 and 2).

We observed gradual NagR regulon expansion in multiple *Bifidobacterium* genomes isolated from mammalian hosts (**Fig. 6A, Table S3**, and **Supplementary Results**). The largest and most complex NagR regulons were identified in strains isolated from the human neonatal gut, namely *B. infantis* ATCC 15697 and *B. bifidum* PRL2010. Notably, while in *B. infantis*, the NagR regulon expanded to include multiple HMO transporters encoded within the H1 cluster (**Fig. 1B**), in *B. bifidum*, the reconstructed regulon contained genes encoding multiple GHs involved in extracellular degradation of HMOs and mucin *O*-glycans (see **Supplementary Results**). Additionally, the NagR regulon expansion was accompanied by minor variations in the NagR-binding motif (**Fig. 6B** and **Supplementary Results**). Taken together, comparative reconstruction of the NagR regulons suggests an evolutionary expansion from a local regulation of GlcNAc catabolism in ancestral bifidobacteria toward a global regulation of various host glycan (e.g., HMOs) utilization in bifidobacterial species isolated from the mammalian neonatal gut.

## Discussion

### Regulation of HMO utilization in *B. infantis*

The predominance of bifidobacteria in the infant gut is linked to their ability to consume and use HMOs as a carbon source (17–21, 34). *B. infantis* possesses a unique genomic locus (H1) that encodes multiple catabolic enzymes and components of ABC transporters that endow this species with the ability to utilize a multitude of HMOs (17, 19, 27). Previous studies demonstrated that a pooled HMO mixture, as well as individual HMOs (LNT and LNnT), induce expression of H1 and *nag* cluster genes in *B. infantis* ATCC 15697, suggesting that the H1 cluster acts as an HMO-inducible unit and is co-regulated with the GlcNAc catabolic pathway (39, 40). However, the regulatory mechanisms underlying this phenomenon were not elucidated.

In this study, we have established NagR-mediated repression of H1, *lnp*, and *nag* cluster genes in *B. infantis* ATCC 15697 by combining PWM-based regulon reconstruction with transcriptome profiling of the *nagR*-KO mutant. The composition of the NagR regulon suggests that this global TF regulates utilization of LNB/GNB, LNT, LNnT, and potentially other decorated (e.g., sialylated) type I and II HMOs in *B. infantis*. We have also demonstrated concentration-dependent binding of recombinant NagR to its predicted operators *in vitro*. The EC_50_ values inferred from EMSA experiments negatively correlated with fold-change values for upregulated genes in the *nagR*-KO mutant. Thus, the degree of gene repression by NagR is strongly dependent on the affinity of this TF to its cognate operator sequences in the promoter regions of corresponding genes.

We identified GlcNAc and its phosphorylated derivatives, GlcNAc-6P and GlcNAc-1P, as potential NagR transcriptional effectors in *B. infantis*. This result is somewhat unexpected since James et al. previously described GlcNAc-6P, but not GlcNAc, as the NagR transcriptional effector in *B. breve* UCC2003 (44). This discrepancy may reflect a metabolic adaptation to a more global nature of NagR regulon in *B. infantis*, although, alternatively, it may be due to the differences in the experimental approach, which in our case was based on the use of purified recombinant NagR for EMSA experiments versus crude cell lysate as in (44). EMSA data indicated that the acetyl group of these intermediary metabolites played a crucial role in NagR-effector interactions, whereas phosphorylation of GlcNAc appeared to be dispensable. While GlcNAc and GlcNAc-6P have been described as transcriptional effectors of the ROK family TFs (44, 48, 49), the potential effector role of GlcNAc-1P is novel and unexpected. Although *N*-acetylhexosamine 1-kinase (LnpB) can phosphorylate GlcNAc to GlcNAc-1P (50), no enzymes that convert the latter to GlcNAc-6P and thus shunt it to the GlcNAc catabolic pathway have been described in prokaryotes (51). In contrast, GlcNAc-1P can be converted to UDP-GlcNAc by GlcNAc-1P uridyltransferase (GlmU) and enter the peptidoglycan biosynthesis pathway (52). Therefore, additional studies are required to assess the biological significance of GlcNAc-1P functioning as a potential NagR transcriptional effector.

Based on the obtained data, we propose a model (**Fig. 7**) where the release of GlcNAc during degradation of LNT and LNnT by intracellular GHs results in derepression of *nag*, *lnp*, and H1 clusters, including genes encoding LNT and LNnT transporters. This model explains the similarity of transcriptomic responses of *B. infantis* ATCC 15697 during growth in media supplemented with LNT or LNnT (39, 40) and suggests that utilization of any GlcNAc-containing glycan by this bacterium will result in upregulation of NagR-controlled genes. Among these glycans might be particular fucosylated HMOs (e.g., LNFP I) and milk *N*-glycans imported by different ABC transport systems (34, 73). Consistent with this notion, a previous study demonstrated that *B. infantis* upregulates *nag* and H1 cluster genes (53) when utilizing *N*-glycosylated human lactoferrin. Our model, however, does not all explain all transcriptional responses observed during HMO utilization by *B. infantis*. The transcriptomic profile of the WT strain grown in MRS-CS-LNnT suggests that additional mechanisms may co-activate the expression of NagR regulon genes (particularly within the *lnp* cluster) in physiologically inducing conditions.

**FIG 7.**
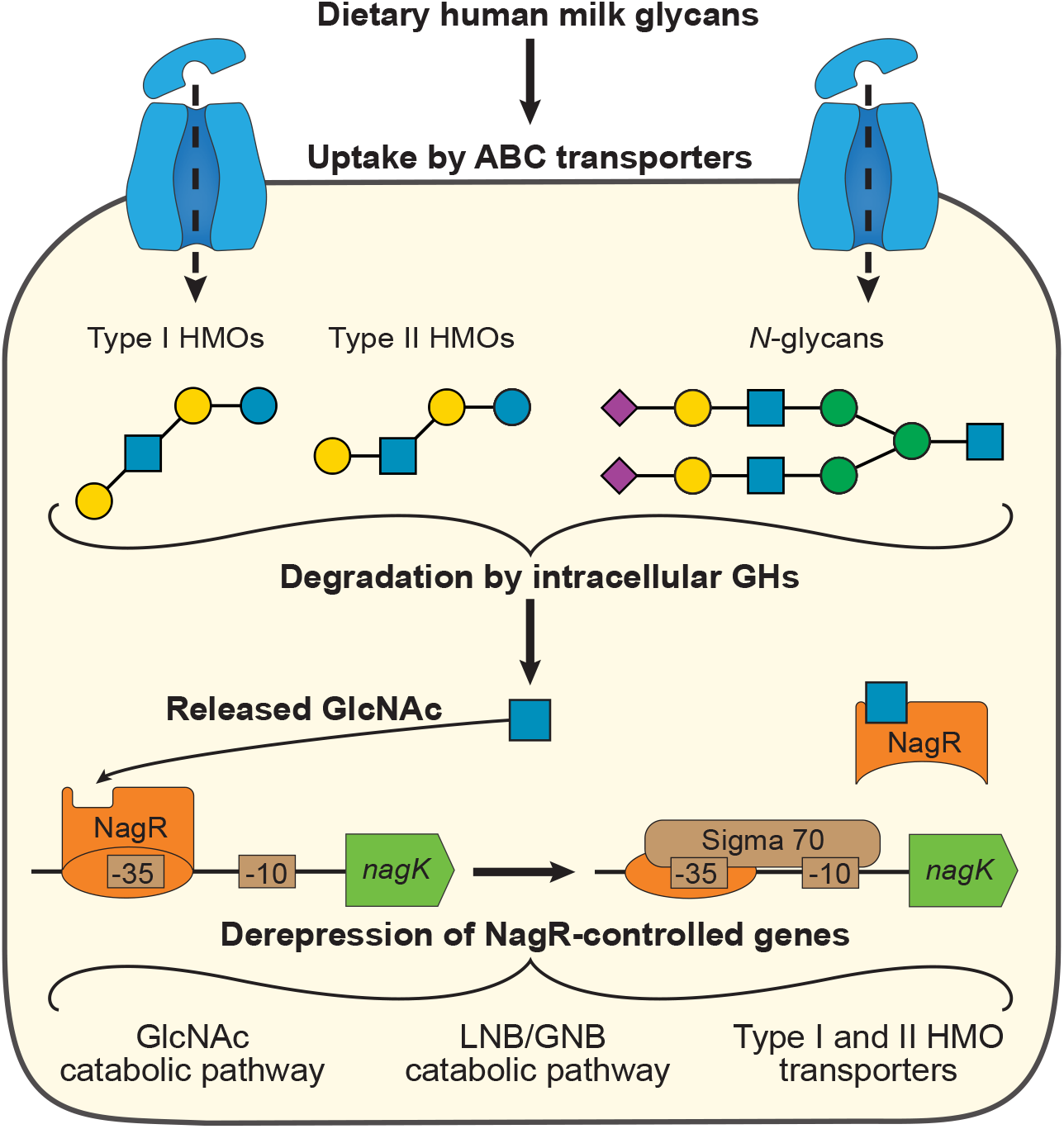
Model of NagR-mediated regulation of HMO utilization in *B. infantis. Step 1*: GlcNAc-containing milk glycans (e.g., LNT and LNnT) are uptaken into the cell by various ABC transporters. *Step 2*: once inside the cell, the glycans are degraded by intracellular GHs, and GlcNAc is released. *Step 3*: released GlcNAc interacts with NagR and disrupts the NagR-operator complex, leading to derepression of NagR-controlled genes.

The structure of the NagR-mediated transcriptional network in *B. infantis* likely reflects the evolutionary adaptation of this bacterium to simultaneous foraging of multiple distinct HMOs and other milk glycans. This notion suggests that using a mixture of LNT and LNnT (and potentially other HMOs) rather than individual oligosaccharides as a prebiotic may be a more efficient solution for selective stimulation of *B. infantis* growth in the neonatal gut since it considers the nuanced regulatory mechanisms and physiology of the target organism.

### Evolution of the NagR regulon in bifidobacteria

Evolution of *B. infantis* was shaped by its ecological niche, specifically the need to forage dietary milk glycans (e.g., HMOs) abundantly present in the gut of breastfed infants (17). This adaptation led to the emergence of several unique genomic clusters, such as H1, that, as we demonstrated, are transcriptionally controlled by NagR. However, other *Bifidobacterium* species inhabiting the neonatal gut, such as *B. breve* and *B. longum*, do not harbor the H1 cluster (42) and have less complex NagR regulons (42, 44). Thus, to elucidate the evolutionary history of the NagR regulon expansion, we reconstructed it in 25 representative genomes spanning 18 *Bifidobacterium* and one *Alloscardovia* species isolated from various hosts and environments.

The regulon structure in early diverged species (e.g., *B. asteroides* and *B. aquikefiri*) suggests that NagR functioned as a local regulator of a single genomic locus involved GlcNAc and possibly *N*,*N*′-diacetylchitobiose catabolism (**Table 2**). Not surprisingly, the activity of this TF would be modulated by GlcNAc or/and GlcNAc-6P. During colonization of mammalian hosts, bifidobacteria were exposed to various GlcNAc-containing glycans, which resulted in the emergence of genomic loci involved in the metabolism of these carbohydrates. For example, most bifidobacteria isolated from mammals (including humans) harbor the *lnp* cluster encoding the LNB/GNB utilization pathway. This observation marks the next evolutionary step of NagR-mediated regulation, namely transitioning from a local to a multi-locus (*nag* and *lnp*) controlling TF. An independent alternative regulon expansion scenario presumably occurred in *B. catenulatum* and *B. pseudocatenulatum* lacking *lnp* genes. In these strains, NagR expanded to control the potential LNB/LNT utilization machinery (*gltABC-nagK-hex1-nagB-nagA* and *bga42A* genes) (**Table 2**). Such regulon structure is consistent with a recently published transcriptomic dataset where LNFP I induced the *gltABC-nagK-hex1-nagB-nagA* cluster in *B. pseudocatenulatum* (54). Although LNFP I is imported into the cell by a different ABC transport system (21, 54), the GlcNAc released during the degradation of this oligosaccharide was likely responsible for the derepression of NagR-controlled genes in this species.

**TABLE 2.** Characteristics of NagR orthologs and paralogs from various *Bifidobacterium* species.

| TF | Species | % identity* | Effector | Regulated pathway | Ref. |
| --- | --- | --- | --- | --- | --- |
| NagR | *B. infantis* ATCC 15697 | 100 | GlcNAc, GlcNAc-1P, GlcNAc-6P | LNB/GNB, LNT, and LNnT utilization | This study |
| NagR | *B. breve* UCC2003 | 84 | GlcNAc-6P | LNB/GNB, LNT utilization | (44) |
| NagR | *B. bifidum* PRL2010 | 60/53** | Unknown | LNB/GNB, HMO, and mucin *O*-glycan utilization<sup>#</sup> | This study |
| NagR | *B. pseudocatenulatum* JCM 1200 | 40 | Unknown | LNB and LNT utilization<sup>#</sup> | This study |
| NagR | *B. asteroides* PRL2011 | 40 | Unknown | GlcNAc catabolism<sup>#</sup> | This study |
| NahR | *B. breve* UCC2003 | 45 | GlcNAc | LNnT utilization | (44) |
| AtsR | *B. breve* UCC2003 | 40 | GlcNAc-6S | GlcNAc-6S utilization | (59) |
| NglR | *B. infantis* Bg 2D9 | 40 | Unknown | Complex *N*-glycan utilization<sup>#</sup> | (73) |

\* Sequence identity with NagR from *B. infantis* ATCC 15697
\*\* Percentages for two NagR copies in *B. bifidum* PRL2010

Multiple further NagR regulon expansion events occurred in prevalent infant-associated species with the highest HMO utilization potential: *B. infantis* and *B. bifidum*. In *B. infantis*, the regulon included multiple H1 cluster genes predominantly encoding HMO transporters, whereas in *B. bifidum*, it expanded to control extracellular GHs involved in HMO and mucin *O*-glycan degradation. These observations provide a fascinating example of how catabolic machinery corresponding to two distinct (convergent) strategies of HMO utilization (intracellular vs. extracellular) became controlled by the same regulatory system. The structure of the NagR regulon partially explains the upregulation of *nag*, *lnp*, and genes encoding extracellular GHs in *B. bifidum* PRL2010 during growth in a mucin-based medium (18). The predicted NagR operators also coincide with a palindromic motif in promoter regions of these upregulated genes (18).

Regulon expansion has been previously described in various bacterial lineages (55, 56), including bifidobacteria (57, 58); however, the underlying evolutionary rationale of such events was not always clear. Here we propose that the general theme of the NagR regulon evolution in the *Bifidobacterium* genus was an expansion to include genes involved in catabolism of GlcNAc-containing host glycans, given the ability of NagR to sense GlcNAc and/or its phosphorylated derivatives. Interestingly, the evolution of gene regulatory networks governing utilization of GlcNAc-containing glycans in bifidobacteria was not limited to the NagR regulon expansion. Another scenario involved duplication(s) of the *nagR* gene after speciation followed by the functional divergence of emerged paralogs. For example, in *B. breve* UCC2003, while NagR controls LNB/GNB and LNT utilization pathways, its paralogs, NahR and AtsR, repress the gene encoding an LNnT transporter and a genomic locus involved in utilization of GlcNAc-6S (a mucin *O*-glycan-constituting saccharide released by 6-sulfo-β-*N*-acetylglucosaminidase BbhII), respectively (44, 59, 60) (**Table 2**). Interestingly, different GlcNAc derivatives serve as effector molecules of these TFs: GlcNAc for NahR and GlcNAc-6S for AtsR (44, 59). Another NagR paralog in *B. infantis* 2D9, NglR, potentially controls a genomic locus involved in utilizing complex *N*-glycans (73) (**Table 2**). Overall, these observations illustrate how bifidobacteria adapted to regulate the foraging of host glycans during colonization of mammalian hosts and shed more light on how complex regulatory networks emerge and evolve.

## Methods

### Reagents and bacterial strains

Reagents were purchased from Alfa-Aesar (Tewksbury, MA, USA), Ambion (Austin, TX, USA), Combi-Blocks (San Diego, CA, USA), Sigma-Aldrich (St. Louis, MO, USA), and Invitrogen (Carlsbad, CA, USA) unless indicated otherwise. Synthetic LNnT (>95% purity) was generously donated by DSM (Heerlen, Netherlands). The HMO mixture was prepared from pooled human milk (21). The experimental protocol was reviewed and approved by the Ethics Committee of Kyoto University (R0046); the study was performed per the Declaration of Helsinki, and informed consent was obtained from all mothers (all subjects). Oligonucleotides were synthesized by Integrated Genomic Technologies (Coralville, IA, USA). Phusion High-Fidelity DNA Polymerase, restriction enzymes, and Quick Ligase were purchased from New England BioLabs (Ipswich, MA, USA).

The type strain of *B. infantis* (ATCC 15697 = JCM 1222) was obtained from Japan Collection of Microorganisms (RIKEN BioResource Research Center, Tsukuba, Japan). *Escherichia coli* DH5α and One Shot TOP10 (Invitrogen) were used for genetic manipulations. *E. coli* BL21/DE3 (New England BioLabs) was used for recombinant NagR overexpression.

### Bioinformatic analysis

A previously established comparative genomic approach was used to identify putative NagR operator sequences and reconstruct regulons in *B. infantis* ATCC 15697 and other selected strains (42, 57). For the initial NagR regulon reconstruction, we built a PWM based on data available in the RegPrecise database (42, 61) using SignalX (62). To improve identification of NagR operators, we built additional PWMs representing two different NagR-binding motifs specific for distant bifidobacterial lineages. Constructed PWMs were used to search for new potential NagR operators using GenomeExplorer (62) with the following parameters: (i) −500 to +50 bp position relative to the first codon of a gene, (iii) site score threshold of 4.3. Identified sites were screened using the consistency check and phylogenetic footprinting approaches to filter out false positives (63). NagR-binding motifs were visualized via WebLogo (64). Positions of the −10 and −35 promoter elements were determined via similar PMW-based searches based on data available for *B. breve* (65) and *B. longum* (66). Promoter regions were aligned using Pro-Coffee (67). Details on additional genomic analysis of *Bifidobacterium* strains and phylogenetic inference are available in Supplementary Methods.

### Targeted *nagR* gene disruption in *B. infantis* ATCC 15697

A single-crossover recombination event was used to inactivate the *nagR* gene (Blon_0880) in *B. infantis* ATCC 15697. Briefly, a BamHI-digested, 2.0 kb fragment of pBS423 (68) that carries the pUC ori and a spectinomycin resistance gene was self-ligated to generate pTK2051, a plasmid incapable of replicating in bifidobacteria. The internal region of *nagR* was then amplified by PCR using primers NagR_I/NagR_II (**Table S1**) and genomic DNA as a template. The amplified 0.5 kbp fragment was inserted into the BamHI site of pTK2051 using the In-Fusion Snap Assembly kit (Takara Bio USA, Mountain View, CA, USA). The resulting suicide plasmid was introduced into *B. infantis* by electroporation (34). Transformed cells were screened for spectinomycin resistance, and the clones were subsequently subjected to a genomic PCR analysis at the *nagR* locus using primers NagR_III/NagR_IV (**Table S1** and **Fig. S2A**). The amplicon was directly sequenced to ensure that the suicide plasmid was integrated into the intended site.

### Culture conditions

*B. infantis* was routinely grown in Gifu anaerobic medium (Nissui Pharmaceutical, Tokyo, Japan) or Lactobacilli MRS Broth w/o Dextrose (MRS-CS; Alpha Biosciences, Baltimore, MD, USA) with 0.34% (w/v) sodium ascorbate, 0.029% (w/v) L-cysteine-HCl monohydrate (BeanTown Chemical, Hudson, NH, USA). The MRS-CS medium was supplemented with lactose, LNnT, or a mixture of neutral HMOs at a final concentration of 1% (w/v). Cultures were incubated at 37 °C in AnaeroPack system (Mitsubishi Gas Chemical Company, Tokyo, Japan) or an anaerobic chamber maintained with a gas mix of 10% H_2_, 10% CO_2_, and 80% N_2_ (Coy Laboratory Products, Grass Lake, MI, USA). Growth was monitored either by measuring culture turbidity in McFarland units using Densitometer DEN-1B (Grant Instruments, Shepreth, UK) or optical density at 600 nm (OD_600_) using DU800 Spectrophotometer (Beckman Coulter, Brea, CA, USA). *E. coli* strains were cultured in Luria-Bertani broth at 37°C with vigorous agitation. Where appropriate, growth media were supplemented with spectinomycin (15 μg/ml for *B. infantis* ATCC 15697 *nagR*-KO, 75 μg/ml for *E. coli* DH5α) or kanamycin (50-60 μg/ml for all other *E. coli* strains). Details on measuring HMO consumption and organic acid production are described in Supplementary Methods.

### Transcriptome analysis (RNA-seq)

For RNA-seq experiments, *B. infantis* ATCC 15697 WT and *nagR*-KO strains were grown as described above in MRS-CS (without spectinomycin) supplemented with either 1% lactose or 1% LNnT. Samples (2 ml; biological triplicates) were collected at the early-mid exponential phase (OD_600_=0.35) and immediately pelleted in a prechilled centrifuge at 4,800 × g for 5 min. Cell pellets were snap-frozen in liquid nitrogen and stored at −80 °C until further use. RNA was extracted as described previously (69) with minor modifications; the detailed protocol can be found in Supplementary Methods. Ribosomal RNA was depleted with NEBNext rRNA Depletion Kit for Bacteria (New England Biolabs). Barcoded libraries were made with NEBNext Ultra II Directional RNA Library Prep Kit for Illumina (New England Biolabs). Libraries were pooled and sequenced (single-end 75 bp reads) on Illumina NextSeq 500 using High Output V2 Kit (Illumina, San Diego, CA, USA). Sequencing data were analyzed as described previously (70) with certain modifications; the detailed pipeline is described in Supplementary Methods.

### Cloning, expression, and purification of recombinant NagR

Codon-optimized nucleotide sequence of *nagR* (Blon_0880) was synthesized by GeneArt Gene Synthesis (Thermo Fisher Scientific, Waltham, MA, USA), PCR-amplified using primers NagR_HisN_F and NagR_HisN_R (**Table S1**), digested by BamHI and SalI, and ligated into a pre-digested in-house pET-49b(+) vector conferring resistance to kanamycin. The ligation mixture was introduced into *E. coli* One Shot TOP10 by chemical transformation, and transformants were then selected based on kanamycin resistance. The recombinant NagR was expressed as fusion with N-terminal His-tag under control of a T7 promoter in *E. coli* BL21/DE3. Cells were grown in LB medium (50 mL) at 37 °C to an OD_600_ of ~0.6 and then transferred to 16 °C. Protein expression was induced by adding 0.2 mM isopropyl-β-D-thiogalactopyranoside. Cells were grown at 16° C overnight and collected by centrifuging at 4,800 × g for 15 min. Harvested cells were resuspended in a lysis buffer containing 10 mM HEPES buffer (pH 7.0), 100 mM NaCl, 0.15% Brij-35, and 5 mM β-mercaptoethanol. Cells were lysed by a freeze-thaw cycle, followed by sonication using Misonix Sonicator 3000 (Misonix Inc, Farmingdale, NY, USA). The cell debris was removed; the soluble fraction was loaded onto a Ni-NTA agarose mini-column (0.2 mL) (Qiagen, Hilden, Germany). The column was washed with 10 column volumes of At buffer (50 mM Tris-HCl (pH 8.0), 500 mM NaCl, 20 mM imidazole, 0.3% Brij-35, 5 mM β-mercaptoethanol) and 10 column volumes of At buffer with 1 M NaCl. Captured proteins were eluted with 0.6 mL of At buffer with 300 mM imidazole. The eluted protein fraction was concentrated and buffer-exchanged into 10 mM Tris-HCl (pH 8.0) with 50 mM NaCl using 30 kDa Amicon Ultra 0.5 mL Centrifugal Filters (MilliporeSigma, Burlington, MA, USA). Protein concentration was determined by Qubit Protein Assay Kit (Invitrogen).

### Electrophoretic mobility shift assays

Oligonucleotides containing predicted 21-bp NagR operators and surrounding genomic regions (14 bp from each end) were synthesized by Integrated DNA Technologies. DNA fragment sequences, sizes, and labels used for testing are given in **Table S1**. Double-stranded labeled DNA probes were obtained by annealing IRD700-labelled oligonucleotides with unlabeled complementary oligonucleotides (ratio 1:5) in 4 mM Tris-HCl (pH 8.0), 20 mM NaCl, 0.4 mM EDTA in Mastercycler PRO Thermal Cycler (Eppendorf) overnight. Binding reactions were carried out with a final volume of 20 μl in binding buffer containing 10 mM Tris–HCl (pH 7.5), 50 mM KCl, 5 mM MgCl_2_, 2.25 mM DTT, 0.125% Tween 20, and 2.5% glycerol. DNA probes (1 nM) were incubated with increasing concentrations of the purified NagR (0-2000 nM) for 60 min at room temperature. Reaction mixtures were loaded on a Novex 6% DNA Retardation gel (Thermo Fisher Scientific) and run in 0.5x Tris-Borate-EDTA buffer (Thermo Fisher Scientific) at 100 V and room temperature for 45 min in XCell SureLock Mini-Cell Electrophoresis System (Thermo Fisher Scientific). Gels were visualized using Odyssey CLx (Li-COR Biosciences, Lincoln, NE, USA). Bands were quantified in Image Studio v5.2 (Li-COR Biosciences). The resulting data were imported into R and approximated by a 4-parameter logistic (4PL) equation in the *drc* package (71) to calculate EC_50_ values. In the 4PL model, the lower limit was fixed at 0 and the upper limit at 1. To identify possible NagR effectors, binding reactions were carried out with 25 nM of NagR and the addition of 0.1-10 mM of GlcNAc or its phosphorylated derivatives (GlcNAc-6P, GlcN-6P, GlcNAc-1P). EC_50_ values for effectors were calculated using the 4PL equation with the upper limit fixed at 1.

## Supporting information

Supplementary Results and Methods

## Data availability

The RNA-seq dataset is deposited on Gene Expression Omnibus (https://www.ncbi.nlm.nih.gov/geo/query/acc.cgi?acc=GSE196064). Raw EMSA gel quantification, growth, HMO consumption, and organic acid production data are available on GitHub (https://github.com/Arzamasov/NagR_manuscript). Code detailing the data analysis steps is available via GitHub and in Supplementary Code File 1.

## Abbreviations

HMOs: human milk oligosaccharides
LNB: lacto-*N*-biose
GNB: galacto-*N*-biose
LNT: lacto-*N*-tetraose
LNnT: lacto-*N*-neotetraose
GlcNAc: *N*-acetylglucosamine
GM: gut microbiota
GH: glycoside hydrolase
ABC: ATP-binding cassette
TF: transcription factor
PWM: position weight matrix
WT: wild-type
PCA: principal component analysis
FC: fold change
EMSA: electrophoretic mobility shift assay
4PL: 4-parameter logistic

## Acknowledgments

We thank Semen Leyn (SBP) for help with computational analyses; Kazi Ahsan (Washington University in St. Louis) for guidance in RNA isolation experiments; Kang Liu and Brian James (SBP Genomics Core) for library preparation and sequencing; Daniel Beiting (University of Pennsylvania) for the DIY.transcriptomics course; Irina Rodionova (University of California San Diego) and Oleg Kurnasov (SBP) for guidance in protein expression and purification experiments; Motomitsu Kitaoka (Niigata University), Junko Hirose (Kyoto Women’s University), Tadasu Urashima (Obihiro University of Agriculture and Veterinary Medicine) for technical support; RIKEN BRC through the National BioResource Project of the MEXT/AMED, Japan for providing pBS423, Glycom A/S and DSM for generously providing LNnT.

This work was supported by JSPS KAKENHI (grant 21H02116 to T.K) and NIH (grant DK30292 to A.L.O).

Conceptualization: A.A.A; data curation: A.A.A; formal analysis: A.A.A, investigation: A.A.A, A.N., M.S., M.N.O; methodology: A.A.A, A.N., M.S., M.N.O; supervision: T.K., D.A.R, A.L.O; visualization: A.A.A; writing – original draft: A.A.A with input from A.N; writing – review & editing: T.K., D.A.R, A.L.O.

A.O. and D.R. are co-founders of Phenobiome Inc., a company pursuing development of computational tools for predictive phenotype profiling of microbial communities. Employment of M.S and M.N.O at Kyoto University is supported by Morinaga Milk Industry Co., Ltd. All other authors declare no conflict of interest.

---

**FIG S1.** Multiple sequence alignments of promoter regions of NagR-regulated genes in *Bifidobacterium* spp. Predicted NagR operators are highlighted in yellow. Predicted promoter elements (−35 and −10 sequences) and in bold black. Experimentally described transcription start sites (TSS) in *B. breve* UCC2003 (65) are in bold blue. Predicted Shine-Dalgarno sequences are in bold red. Coding sequences are highlighted in grey; initiation codons are underlined. Conserved nucleotides are marked with asterisks.

**FIG S2**. Construction and metabolic profiling of the *B. infantis* ATCC 15697 *nagR*-KO mutant. (A) Schematic representation of insertional inactivation of *nagR*. The region used for a single crossover recombination event is in orange. The primers for genomic PCR (green) were designed to anneal outside the region used for recombination. (B) Results of genomic PCR. The amplicon sizes were expected to be 679 bp and 3164 bp for WT and *nagR*-KO strains, respectively. Four clones were analyzed. (C) HMO consumption and (D) organic acid production profiles of *B. infantis* ATCC 15697 WT and *nagR*-KO strains grown in MRS-CS-HMO. Data points represent the mean of three biological replicates. Error bars depict 95% confidence intervals for the mean. Timepoints where the metabolite concentrations for WT and *nagR*-KO strains were significantly different (*, *P*_adj_<0.05) were identified using a linear regression. Bonferroni correction was used to adjust for multiple testing.

**FIG S3**. Heatmap depicting the expression of genes from Table S2A-B across four experimental conditions (WT and *nagR*-KO strains grown in MRS-CS supplemented either with Lac or LNnT). Genes constituting the predicted NagR regulon are in bold. Genomic clusters from Fig. 1B are marked by lines.

**FIG S4**. (A) EMSA gels depicting titration of DNA probes (1 nM) containing predicted NagR operators with recombinant NagR. (B) EMSA gels depicting titration of the NagR-DNA probe complex with various effector molecules. 25 nM of NagR and 1 nM of the *hmoA* probe were used in all reactions.

**Table legends**

**TABLE S1**. Oligonucleotides used in this study. IRD oligonucleotides are 5’-labeled with the IRD700 dye, whereas RC oligonucleotides are unlabeled. Predicted NagR operators are in bold.

**TABLE S2**. List of differentially expressed genes (P_adj_ < 0.01; absolute log_2_FC > 1) between selected experimental conditions in the RNA-seq experiment. (A) *nagR*-KO grown in MRS-CS-Lac vs. WT grown in MRS-CS-Lac. (B) WT grown in MRS-CS-LNnT vs. WT grown in MRS-CS-Lac.

**TABLE S3**. Reconstructed NagR regulons and predicted NagR operators in 25 *Bifidobacteriaceae* genomes.

## References

1. Alessandri G, van Sinderen D, Ventura M. 2021. The genus bifidobacterium: From genomics to functionality of an important component of the mammalian gut microbiota running title: Bifidobacterial adaptation to and interaction with the host. Comput Struct Biotechnol J 19:1472–1487.

2. Matsuki T, Yahagi K, Mori H, Matsumoto H, Hara T, Tajima S, Ogawa E, Kodama H, Yamamoto K, Yamada T, Matsumoto S, Kurokawa K. 2016. A key genetic factor for fucosyllactose utilization affects infant gut microbiota development. Nat Commun 7 11939.

3. Vatanen T, Franzosa EA, Schwager R, Tripathi S, Arthur TD, Vehik K, Lernmark Å, Hagopian WA, Rewers MJ, She J-X, Toppari J, Ziegler A-G, Akolkar B, Krischer JP, Stewart CJ, Ajami NJ, Petrosino JF, Gevers D, Lähdesmäki H, Vlamakis H, Huttenhower C, Xavier RJ. 2018. The human gut microbiome in early-onset type 1 diabetes from the TEDDY study. Nature 562:589–594.

4. Tsukuda N, Yahagi K, Hara T, Watanabe Y, Matsumoto H, Mori H, Higashi K, Tsuji H, Matsumoto S, Kurokawa K, Matsuki T. 2021. Key bacterial taxa and metabolic pathways affecting gut short-chain fatty acid profiles in early life. ISME J https://doi.org/10.1038/s41396-021-00937-7.

5. Laursen MF, Sakanaka M, von Burg N, Mörbe U, Andersen D, Moll JM, Pekmez CT, Rivollier A, Michaelsen KF, Mølgaard C, Lind MV, Dragsted LO, Katayama T, Frandsen HL, Vinggaard AM, Bahl MI, Brix S, Agace W, Licht TR, Roager HM. 2021. Bifidobacterium species associated with breastfeeding produce aromatic lactic acids in the infant gut. Nat Microbiol 6:1367–1382.

6. Raman AS, Gehrig JL, Venkatesh S, Chang H-W, Hibberd MC, Subramanian S, Kang G, Bessong PO, Lima AAM, Kosek MN, Petri WA, Rodionov DA, Arzamasov AA, Leyn SA, Osterman AL, Huq S, Mostafa I, Islam M, Mahfuz M, Haque R, Ahmed T, Barratt MJ, Gordon JI. 2019. A sparse covarying unit that describes healthy and impaired human gut microbiota development. Science 365.

7. Gehrig JL, Venkatesh S, Chang H-W, Hibberd MC, Kung VL, Cheng J, Chen RY, Subramanian S, Cowardin CA, Meier MF, O’Donnell D, Talcott M, Spears LD, Semenkovich CF, Henrissat B, Giannone RJ, Hettich RL, Ilkayeva O, Muehlbauer M, Newgard CB, Sawyer C, Head RD, Rodionov DA, Arzamasov AA, Leyn SA, Osterman AL, Hossain MI, Islam M, Choudhury N, Sarker SA, Huq S, Mahmud I, Mostafa I, Mahfuz M, Barratt MJ, Ahmed T, Gordon JI. 2019. Effects of microbiota-directed foods in gnotobiotic animals and undernourished children. Science 365.

8. Frese SA, Hutton AA, Contreras LN, Shaw CA, Palumbo MC, Casaburi G, Xu G, Davis JCC, Lebrilla CB, Henrick BM, Freeman SL, Barile D, German JB, Mills DA, Smilowitz JT, Underwood MA. 2017. Persistence of Supplemented Bifidobacterium longum subsp. infantis EVC001 in Breastfed Infants. mSphere 2:e00501–17.

9. Casaburi G, Duar RM, Vance DP, Mitchell R, Contreras L, Frese SA, Smilowitz JT, Underwood MA. 2019. Early-life gut microbiome modulation reduces the abundance of antibiotic-resistant bacteria. Antimicrob Resist Infect Control 8:131.

10. Hirano R, Sakanaka M, Yoshimi K, Sugimoto N, Eguchi S, Yamauchi Y, Nara M, Maeda S, Ami Y, Gotoh A, Katayama T, Iida N, Kato T, Ohno H, Fukiya S, Yokota A, Nishimoto M, Kitaoka M, Nakai H, Kurihara S. 2021. Next-generation prebiotic promotes selective growth of bifidobacteria, suppressing Clostridioides difficile. Gut Microbes 13:1973835.

11. Henrick BM, Rodriguez L, Lakshmikanth T, Pou C, Henckel E, Arzoomand A, Olin A, Wang J, Mikes J, Tan Z, Chen Y, Ehrlich AM, Bernhardsson AK, Mugabo CH, Ambrosiani Y, Gustafsson A, Chew S, Brown HK, Prambs J, Bohlin K, Mitchell RD, Underwood MA, Smilowitz JT, German JB, Frese SA, Brodin P. 2021. Bifidobacteria-mediated immune system imprinting early in life. Cell 184:3884–3898.e11.

12. Fukuda S, Toh H, Hase K, Oshima K, Nakanishi Y, Yoshimura K, Tobe T, Clarke JM, Topping DL, Suzuki T, Taylor TD, Itoh K, Kikuchi J, Morita H, Hattori M, Ohno H. 2011. Bifidobacteria can protect from enteropathogenic infection through production of acetate. Nature 469:543–547.

13. Alessandri G, Ossiprandi MC, MacSharry J, van Sinderen D, Ventura M. 2019. Bifidobacterial Dialogue With Its Human Host and Consequent Modulation of the Immune System. Front Immunol 10:2348.

14. Alcon-Giner C, Dalby MJ, Caim S, Ketskemety J, Shaw A, Sim K, Lawson MAE, Kiu R, Leclaire C, Chalklen L, Kujawska M, Mitra S, Fardus-Reid F, Belteki G, McColl K, Swann JR, Kroll JS, Clarke P, Hall LJ. 2020. Microbiota Supplementation with Bifidobacterium and Lactobacillus Modifies the Preterm Infant Gut Microbiota and Metabolome: An Observational Study. Cell Rep Med 1:100077.

15. Nguyen M, Holdbrooks H, Mishra P, Abrantes MA, Eskew S, Garma M, Oca C-G, McGuckin C, Hein CB, Mitchell RD, Kazi S, Chew S, Casaburi G, Brown HK, Frese SA, Henrick BM. 2021. Impact of Probiotic B. infantis EVC001 Feeding in Premature Infants on the Gut Microbiome, Nosocomially Acquired Antibiotic Resistance, and Enteric Inflammation. Front Pediatr 9:618009.

16. Bajorek S, Duar RM, Corrigan M, Matrone C, Winn KA, Norman S, Mitchell RD, Cagney O, Aksenov AA, Melnik AV, Kopylova E, Perez J. 2021. B. infantis EVC001 Is Well-Tolerated and Improves Human Milk Oligosaccharide Utilization in Preterm Infants in the Neonatal Intensive Care Unit. Front Pediatr 9:795970.

17. Sela DA, Chapman J, Adeuya A, Kim JH, Chen F, Whitehead TR, Lapidus A, Rokhsar DS, Lebrilla CB, German JB, Price NP, Richardson PM, Mills DA. 2008. The genome sequence of Bifidobacterium longum subsp. infantis reveals adaptations for milk utilization within the infant microbiome. Proc Natl Acad Sci USA 105:18964–18969.

18. Turroni F, Bottacini F, Foroni E, Mulder I, Kim J-H, Zomer A, Sánchez B, Bidossi A, Ferrarini A, Giubellini V, Delledonne M, Henrissat B, Coutinho P, Oggioni M, Fitzgerald GF, Mills D, Margolles A, Kelly D, van Sinderen D, Ventura M. 2010. Genome analysis of Bifidobacterium bifidum PRL2010 reveals metabolic pathways for host-derived glycan foraging. Proc Natl Acad Sci USA 107:19514–19519.

19. Asakuma S, Hatakeyama E, Urashima T, Yoshida E, Katayama T, Yamamoto K, Kumagai H, Ashida H, Hirose J, Kitaoka M. 2011. Physiology of consumption of human milk oligosaccharides by infant gut-associated bifidobacteria. J Biol Chem 286:34583–34592.

20. James K, Motherway MO, Bottacini F, van Sinderen D. 2016. Bifidobacterium breve UCC2003 metabolises the human milk oligosaccharides lacto-N-tetraose and lacto-N-neo-tetraose through overlapping, yet distinct pathways. Sci Rep 6.

21. Ojima MN, Asao Y, Nakajima A, Katoh T, Kitaoka M, Gotoh A, Hirose J, Urashima T, Fukiya S, Yokota A, Abou Hachem M, Sakanaka M, Katayama T. 2021. Diversification of a fucosyllactose transporter within the genus Bifidobacterium. Appl Environ Microbiol AEM0143721.

22. Kunz C, Rudloff S, Baier W, Klein N, Strobel S. 2000. Oligosaccharides in human milk: structural, functional, and metabolic aspects. Annu Rev Nutr 20:699–722.

23. Bode L. 2012. Human milk oligosaccharides: every baby needs a sugar mama. Glycobiology 22:1147–1162.

24. Ballard O, Morrow AL. 2013. Human Milk Composition: Nutrients and Bioactive Factors. Pediatr Clin North Am 60:49–74.

25. Wu S, Tao N, German JB, Grimm R, Lebrilla CB. 2010. Development of an annotated library of neutral human milk oligosaccharides. J Proteome Res 9:4138–4151.

26. Chen X. 2015. Human Milk Oligosaccharides (HMOS): Structure, Function, and Enzyme-Catalyzed Synthesis. Adv Carbohydr Chem Biochem 72:113–190.

27. Sakanaka M, Gotoh A, Yoshida K, Odamaki T, Koguchi H, Xiao J-Z, Kitaoka M, Katayama T. 2019. Varied Pathways of Infant Gut-Associated Bifidobacterium to Assimilate Human Milk Oligosaccharides: Prevalence of the Gene Set and Its Correlation with Bifidobacteria-Rich Microbiota Formation. Nutrients 12.

28. Katayama T, Sakuma A, Kimura T, Makimura Y, Hiratake J, Sakata K, Yamanoi T, Kumagai H, Yamamoto K. 2004. Molecular cloning and characterization of Bifidobacterium bifidum 1,2-alpha-L-fucosidase (AfcA), a novel inverting glycosidase (glycoside hydrolase family 95). J Bacteriol 186:4885–4893.

29. Wada J, Ando T, Kiyohara M, Ashida H, Kitaoka M, Yamaguchi M, Kumagai H, Katayama T, Yamamoto K. 2008. Bifidobacterium bifidum lacto-N-biosidase, a critical enzyme for the degradation of human milk oligosaccharides with a type 1 structure. Appl Environ Microbiol 74:3996–4004.

30. Ashida H, Miyake A, Kiyohara M, Wada J, Yoshida E, Kumagai H, Katayama T, Yamamoto K. 2009. Two distinct alpha-L-fucosidases from Bifidobacterium bifidum are essential for the utilization of fucosylated milk oligosaccharides and glycoconjugates. Glycobiology 19:1010–1017.

31. Miwa M, Horimoto T, Kiyohara M, Katayama T, Kitaoka M, Ashida H, Yamamoto K. 2010. Cooperation of β-galactosidase and β-N-acetylhexosaminidase from bifidobacteria in assimilation of human milk oligosaccharides with type 2 structure. Glycobiology 20:1402–1409.

32. Kiyohara M, Tanigawa K, Chaiwangsri T, Katayama T, Ashida H, Yamamoto K. 2011. An exo-alpha-sialidase from bifidobacteria involved in the degradation of sialyloligosaccharides in human milk and intestinal glycoconjugates. Glycobiology 21:437–447.

33. Garrido D, Kim JH, German JB, Raybould HE, Mills DA. 2011. Oligosaccharide Binding Proteins from Bifidobacterium longum subsp. infantis Reveal a Preference for Host Glycans. PLoS One 6.

34. Sakanaka M, Hansen ME, Gotoh A, Katoh T, Yoshida K, Odamaki T, Yachi H, Sugiyama Y, Kurihara S, Hirose J, Urashima T, Xiao J-Z, Kitaoka M, Fukiya S, Yokota A, Lo Leggio L, Abou Hachem M, Katayama T. 2019. Evolutionary adaptation in fucosyllactose uptake systems supports bifidobacteria-infant symbiosis. Sci Adv 5:eaaw7696.

35. Sela DA, Li Y, Lerno L, Wu S, Marcobal AM, German JB, Chen X, Lebrilla CB, Mills DA. 2011. An infant-associated bacterial commensal utilizes breast milk sialyloligosaccharides. J Biol Chem 286:11909–11918.

36. Garrido D, Ruiz-Moyano S, Mills DA. 2012. Release and utilization of N-acetyl-d-glucosamine from human milk oligosaccharides by Bifidobacterium longum subsp. infantis. Anaerobe 18:430–435.

37. Sela DA, Garrido D, Lerno L, Wu S, Tan K, Eom H-J, Joachimiak A, Lebrilla CB, Mills DA. 2012. Bifidobacterium longum subsp. infantis ATCC 15697 α-Fucosidases Are Active on Fucosylated Human Milk Oligosaccharides. Appl Environ Microbiol 78:795–803.

38. Yoshida E, Sakurama H, Kiyohara M, Nakajima M, Kitaoka M, Ashida H, Hirose J, Katayama T, Yamamoto K, Kumagai H. 2012. Bifidobacterium longum subsp. infantis uses two different β-galactosidases for selectively degrading type-1 and type-2 human milk oligosaccharides. Glycobiology 22:361–368.

39. Garrido D, Ruiz-Moyano S, Lemay DG, Sela DA, German JB, Mills DA. 2015. Comparative transcriptomics reveals key differences in the response to milk oligosaccharides of infant gut-associated bifidobacteria. Sci Rep 5.

40. Özcan E, Sela DA. 2018. Inefficient Metabolism of the Human Milk Oligosaccharides Lacto-N-tetraose and Lacto-N-neotetraose Shifts Bifidobacterium longum subsp. infantis Physiology. Front Nutr 5.

41. Ravcheev DA, Godzik A, Osterman AL, Rodionov DA. 2013. Polysaccharides utilization in human gut bacterium Bacteroides thetaiotaomicron: comparative genomics reconstruction of metabolic and regulatory networks. BMC Genomics 14:873.

42. Khoroshkin MS, Leyn SA, Van Sinderen D, Rodionov DA. 2016. Transcriptional Regulation of Carbohydrate Utilization Pathways in the Bifidobacterium Genus. Front Microbiol 7.

43. Rodionov DA, Rodionova IA, Rodionov VA, Arzamasov AA, Zhang K, Rubinstein GM, Tanwee TNN, Bing RG, Crosby JR, Nookaew I, Basen M, Brown SD, Wilson CM, Klingeman DM, Poole FL, Zhang Y, Kelly RM, Adams MWW. 2021. Transcriptional Regulation of Plant Biomass Degradation and Carbohydrate Utilization Genes in the Extreme Thermophile Caldicellulosiruptor bescii. mSystems e0134520.

44. James K, O’Connell Motherway M, Penno C, O’Brien RL, van Sinderen D. 2018. Bifidobacterium breve UCC2003 Employs Multiple Transcriptional Regulators To Control Metabolism of Particular Human Milk Oligosaccharides. Appl Environ Microbiol 84.

45. Kazanov MD, Li X, Gelfand MS, Osterman AL, Rodionov DA. 2013. Functional diversification of ROK-family transcriptional regulators of sugar catabolism in the Thermotogae phylum. Nucleic Acids Res 41:790–803.

46. Ritchie ME, Phipson B, Wu D, Hu Y, Law CW, Shi W, Smyth GK. 2015. limma powers differential expression analyses for RNA-sequencing and microarray studies. Nucleic Acids Res 43:e47.

47. Warchol M, Perrin S, Grill J-P, Schneider F. 2002. Characterization of a purified beta-fructofuranosidase from Bifidobacterium infantis ATCC 15697. Lett Appl Microbiol 35:462–467.

48. Plumbridge JA. 1991. Repression and induction of the nag regulon of Escherichia coli K-12: the roles of nagC and nagA in maintenance of the uninduced state. Mol Microbiol 5:2053–2062.

49. Plumbridge J, Kolb A. 1991. CAP and Nag repressor binding to the regulatory regions of the nagE-B and manX genes of Escherichia coli. J Mol Biol 217:661–679.

50. Nishimoto M, Kitaoka M. 2007. Identification of N-Acetylhexosamine 1-Kinase in the Complete Lacto-N-Biose I/Galacto-N-Biose Metabolic Pathway in Bifidobacterium longum. Appl Environ Microbiol 73:6444–6449.

51. Stiers KM, Muenks AG, Beamer LJ. 2017. Biology, mechanism, and structure of enzymes in the α-D-phosphohexomutase superfamily. Adv Protein Chem Struct Biol 109:265–304.

52. Barreteau H, Kovac A, Boniface A, Sova M, Gobec S, Blanot D. 2008. Cytoplasmic steps of peptidoglycan biosynthesis. FEMS Microbiol Rev 32:168–207.

53. Garrido D, Nwosu C, Ruiz-Moyano S, Aldredge D, German JB, Lebrilla CB, Mills DA. 2012. Endo-β-N-acetylglucosaminidases from infant gut-associated bifidobacteria release complex N-glycans from human milk glycoproteins. Mol Cell Proteomics 11:775–785.

54. Shani G, Hoeflinger JL, Heiss BE, Masarweh CF, Larke JA, Jensen NM, Wickramasinghe S, Davis JC, Goonatilleke E, El-Hawiet A, Nguyen L, Klassen JS, Slupsky CM, Lebrilla CB, Mills DA. 2021. Fucosylated human milk oligosaccharide foraging within the species Bifidobacterium pseudocatenulatum is driven by glycosyl hydrolase content and specificity. Appl Environ Microbiol AEM0170721.

55. Leyn SA, Li X, Zheng Q, Novichkov PS, Reed S, Romine MF, Fredrickson JK, Yang C, Osterman AL, Rodionov DA. 2011. Control of Proteobacterial Central Carbon Metabolism by the HexR Transcriptional Regulator. J Biol Chem 286:35782–35794.

56. Ravcheev DA, Khoroshkin MS, Laikova ON, Tsoy OV, Sernova NV, Petrova SA, Rakhmaninova AB, Novichkov PS, Gelfand MS, Rodionov DA. 2014. Comparative genomics and evolution of regulons of the LacI-family transcription factors. Front Microbiol 5.

57. Arzamasov AA, van Sinderen D, Rodionov DA. 2018. Comparative Genomics Reveals the Regulatory Complexity of Bifidobacterial Arabinose and Arabino-Oligosaccharide Utilization. Front Microbiol 9:776.

58. Lanigan N, Kelly E, Arzamasov AA, Stanton C, Rodionov DA, van Sinderen D. 2019. Transcriptional control of central carbon metabolic flux in Bifidobacteria by two functionally similar, yet distinct LacI-type regulators. Sci Rep 9:17851.

59. Egan M, Jiang H, O’Connell Motherway M, Oscarson S, van Sinderen D. 2016. Glycosulfatase-Encoding Gene Cluster in Bifidobacterium breve UCC2003. Appl Environ Microbiol 82:6611–6623.

60. Katoh T, Maeshibu T, Kikkawa K-I, Gotoh A, Tomabechi Y, Nakamura M, Liao W-H, Yamaguchi M, Ashida H, Yamamoto K, Katayama T. 2017. Identification and characterization of a sulfoglycosidase from Bifidobacterium bifidum implicated in mucin glycan utilization. Biosci Biotechnol Biochem 81:2018–2027.

61. Novichkov PS, Laikova ON, Novichkova ES, Gelfand MS, Arkin AP, Dubchak I, Rodionov DA. 2010. RegPrecise: a database of curated genomic inferences of transcriptional regulatory interactions in prokaryotes. Nucleic Acids Res 38:D111–D118.

62. Mironov AA, Vinokurova NP, Gelfand MS. 2000. Software for analysis of bacterial genomes. Mol Biol 34:222–231.

63. Rodionov DA. 2007. Comparative Genomic Reconstruction of Transcriptional Regulatory Networks in Bacteria. Chem Rev 107:3467–3497.

64. Crooks GE, Hon G, Chandonia J-M, Brenner SE. 2004. WebLogo: A Sequence Logo Generator. Genome Res 14:1188–1190.

65. Bottacini F, Zomer A, Milani C, Ferrario C, Lugli GA, Egan M, Ventura M, van Sinderen D. 2017. Global transcriptional landscape and promoter mapping of the gut commensal Bifidobacterium breve UCC2003. BMC Genomics 18.

66. Kozakai T, Izumi A, Horigome A, Odamaki T, Xiao J, Nomura I, Suzuki T. 2020. Structure of a Core Promoter in Bifidobacterium longum NCC2705. J Bacteriol 202.

67. Erb I, González-Vallinas JR, Bussotti G, Blanco E, Eyras E, Notredame C. 2012. Use of ChIP-Seq data for the design of a multiple promoter-alignment method. Nucleic Acids Res 40:e52.

68. Hirayama Y, Sakanaka M, Fukuma H, Murayama H, Kano Y, Fukiya S, Yokota A. 2012. Development of a Double-Crossover Markerless Gene Deletion System in Bifidobacterium longum: Functional Analysis of the α-Galactosidase Gene for Raffinose Assimilation. Appl Environ Microbiol 78:4984–4994.

69. Rey FE, Faith JJ, Bain J, Muehlbauer MJ, Stevens RD, Newgard CB, Gordon JI. 2010. Dissecting the in vivo metabolic potential of two human gut acetogens. J Biol Chem 285:22082–22090.

70. Amorim CF, Novais FO, Nguyen BT, Misic AM, Carvalho LP, Carvalho EM, Beiting DP, Scott P. 2019. Variable gene expression and parasite load predict treatment outcome in cutaneous leishmaniasis. Sci Transl Med 11.

71. Ritz C, Baty F, Streibig JC, Gerhard D. 2015. Dose-Response Analysis Using R. PLoS One 10:e0146021.

72. Kitaoka M, Tian J, Nishimoto M. 2005. Novel Putative Galactose Operon Involving Lacto-N-Biose Phosphorylase in Bifidobacterium longum. Appl Environ Microbiol 71:3158–3162.

73. Barratt M, Nuzhat S, Ahsan K, Frese SA, Arzamasov AA, Sarker SA, Islam MM, Palit P, Islam MR, Hibberd MC, Nakshatri S, Cowardin CA, Guruge JL, Byrne AE, Venkatesh S, Sundaresan V, Henrick B, Duar RM, Mitchell RD, Casaburi G, Prambs J, Flannery R, Mahfuz M, Rodionov DA, Osterman AL, Kyle D, Ahmed T, Gordon J. 2022. Bifidobacterium longum subsp. infantis strains for treating severe acute malnutrition in Bangladeshi infants. Sci Transl Med in press.

