## Supplementary Results and Methods for "Human Milk Oligosaccharide Utilization in Intestinal Bifidobacteria is Governed by a Global Transcriptional Regulator NagR"

### Supplementary methods

#### Genomic analysis of *Bifidobacteriaceae* strains

Context-driven identification of orthologous genes and reconstruction of HMO utilization pathways were performed in the web-based mcSEED environment, a private clone of the publicly available SEED platform (1). Genomes of 25 strains representing different lineages within the *Bifidobacteriaceae* family were annotated using RAST (2) and imported into mcSEED. Information on functional roles (transporters, glycoside hydrolases, downstream catabolic enzymes, transcriptional regulators) involved in bifidobacterial HMO utilization were collected by extensive literature search using PaperBLAST (3) and by exporting data from Carbohydrate Active Enzyme (CAZy) (4), Transporter Classification (TCDB) (5), and RegPrecise (6) databases.

#### Phylogenetic inference

RAST-annotated versions of genomes were converted to the gff3 format and used as input in Roary (v3.12.0) (7). Pangenome was calculated using a 70 % minimum percentage identity threshold for blastp (8), and 247 core genes were identified. Concatenated nucleotide sequences of the core genes were aligned via MAFFT (v7.313) (9). NagR protein sequences were aligned using MUSCLE (v3.8.31) (10). The resulting multiple sequence alignments were trimmed by the *msaTrim* function in the *microseq* package (11). Phylogenetic trees were built via the maximum likelihood approach implemented in IQ-TREE (v1.6.12); automatic model selection and ultrafast bootstrap (1000 replicates) options were used (12).

#### Analysis of HMO consumption and organic acid production

*B. infantis* ATCC 15697 WT and *nagR*-KO strains grown in the Gifu anaerobic medium were harvested, washed twice with sugar-free MRS-CS, and then used to inoculate MRS-CS medium supplemented with lactose or an HMO mixture (1% w/v) at $OD_{600}$=0.01. Samples (biological triplicates) were collected at indicated time points (0, 4, 8, 12, and 24 h) for monitoring growth ($OD_{600}$) and metabolic profiling. The supernatants were used to analyze HMO consumption and organic acid production. Sugars in the spent media were fluorescence-labeled with anthranilic acid and quantified by normal-phase high-performance liquid chromatography (HPLC) as described previously (13, 14). Fucose in the culture supernatants was quantified using high-performance anion-exchange chromatography with pulsed amperometric detection (HPAEC-PAD) as described previously (15). Organic acids (lactic acid, acetic acid, formic acid, butyric acid, propionic acid, butyric acid, *iso*-valeric acid, valeric acid, and succinic acid) were quantified by anion-exchange chromatography as described previously (16). A linear regression and post hoc comparisons implemented in the *emmeans* package (17) were used to identify time points where the mean concentrations of HMOs/organic acids for WT and *nagR*-KO strains were significantly different. Computed p-values were adjusted for multiple comparisons using Bonferroni correction.

**RNA isolation**

Frozen cell pellets were kept on dry ice and resuspended in a solution containing 250 µL of acid-washed glass beads (212-300 µm), 710 µL of extraction mixture (200 mM NaCl, 20 mM EDTA, 20% SDS), and 500 µL of a mixture of phenol:chloroform:isoamyl alcohol (125:24:1, pH 4.5). Cells were disrupted using Bead Ruptor 12 (Omni International, Kennesaw, GA, USA) by alternating 2 min of homogenizing at 6 m/s and 2 min of cooling on ice. Cellular debris was removed by centrifugation (16,000 × g for 10 min), and the aqueous phase was collected. RNA was precipitated with isopropanol and sodium acetate (pH 5.5) at −20 °C overnight, washed in ice-cold 70% ethanol, and resuspended in 100 µL of nuclease-free water. Total RNA was subjected to two consecutive DNase treatments (30 min each) with Baseline-ZERO DNase (Lucigen, Middleton, WI, USA) and Turbo DNase (Ambion), respectively. Each treatment was followed by cleanup using MEGAclear Transcription Clean-Up Kit (Ambion). RNA was quantified via Qubit Broad Range RNA Assay Kit (Invitrogen), and RNA integrity was confirmed by gel electrophoresis in 1 % agarose and 4200 TapeStation System (Agilent, Santa Clara, CA, USA). All samples had RIN > 7.5. PCR-based checks for potential contamination by residual genomic DNA were performed using primers NagR_III/NagR_IV (**Table S1**).

**RNA-seq data processing**

Raw reads were demultiplexed and quality checked via FastQC (v0.11.9) (18). Illumina sequencing adapters and short reads (< 20 bp) were trimmed by Cutadapt (v3.4) (19). Reads were aligned to rRNA gene sequences extracted from the *B. infantis* ATCC 15697 genome (GenBank accession no. CP001095.1; locus tags Blon_xxxx) using Bowtie2 (v2.4.4) (20) to filter out rRNA reads. Unmapped (filtered) reads were pseudoaligned to the *B. infantis* ATCC 15697 transcriptome via Kallisto (v0.46.2) (21). All subsequent analyses were carried out using R (v4.1.2) in RStudio (v1.4.1717) and Bioconductor (v.3.14) (22). Kallisto transcript quantification data were imported using the *TxImport* package (23) and normalized by the TMM method in the *edgeR* package (24). Genes with < 1 count per million (CPM) in ≤ 3 samples were filtered out. PCA was performed on TMM-normalized count data to control for potential batch effects. Normalized, filtered data were variance-stabilized using the *voom* function in *limma* (25), and differentially expressed genes were identified via linear modeling in *limma* ($P_{adj} < 0.01$; absolute $\log_2$FC > 1) after correcting for multiple testing using the Benjamini-Hochberg procedure (26). PCA and volcano plots were visualized using *ggplot2* (27). Heatmaps were created and visualized using the *pheatmap* package (28); row values were z-transformed and colored according to corresponding z–scores.

### Supplementary results

#### Evolution of the NagR regulon in the *Bifidobacteriaceae* family

To trace the potential evolution of the NagR regulon, we used the PWM-based approach and reconstructed NagR regulons in 25 *Bifidobacteriaceae* genomes, encompassing 18 *Bifidobacterium* and one *Alloscardovia* species harboring NagR orthologs. The genomes were chosen to adequately represent the phylogenetic diversity within the *Bifidobacterium* genus and emphasize strain-specific variation within *B. infantis* and *B. longum* subspecies.

NagR regulons in *Bifidobacterium asteroides* PRL2011 and *Bifidobacterium aquikefiri* LMG 28769 were composed of a cluster containing *nagK*, *nagBA*, and *nagR* genes (homologous to the *nag* cluster genes in *B. infantis*, see **Fig 1B**). Notably, putative NagR-mediated regulation of these four genes was conserved in 22 out of 25 genomes. Reconstructed NagR regulons in *Bifidobacterium psychraerophilum* LMG 21775, *Bifidobacterium bohemicum* DSM 22767, and *Bifidobacterium mongoliense* DSM 21395 additionally contained genes encoding orthologs of β-*N*-acetylglucosaminidase Hex1 (GH20) (29) and a predicted ABC transport system for *N*,*N*'-diacetylchitobiose (ChbABC) (**Fig. 6A** and **Table S3**). These observations suggest that in ancestral bifidobacteria, NagR might function as a local regulator of genomic loci involved in GlcNAc catabolism.

We observed multiple NagR regulon expansion events in *Bifidobacterium* strains isolated from mammalian hosts (**Fig. 6A**). So, the reconstructed regulons in 13 *Bifidobacterium* and *Alloscardovia theropitheci* GLDI4/2 genomes also contained *gltABC* and *lnpABCD* genes (homologous to the *lnp* cluster genes in *B. infantis*, see **Fig. 1B**). Interestingly, three *Bifidobacterium catenulatum*/*pseudocatenulatum* group strains isolated from the human neonatal gut lacked *lnpABCD* orthologs encoding the LNB/GNB catabolic pathway but retained *gltABC* orthologs encoding an LNB/type I HMO (e.g., LNT) transporter. In these strains, NagR regulons consisted of one genomic cluster with *nagBA*, *nagK*, *nagR*, *gltABC*, and *hex1* genes, as well a separate gene encoding an ortholog of β-galactosidase Bga42A (GH42) implicated in LNT utilization in *B. breve* and *B. infantis* (30–32) (**Fig. 6A** and **Table S3**). Notably, orthologs of *bga42A* were found in 20 other genomes; however, we did not identify any putative NagR operators in promoter regions of these genes. Based on the composition of NagR regulons in *B. catenulatum* and *B. pseudocatenulatum*, we propose that this TF might function as a regulator of LNB and LNT utilization in these species.

The largest NagR regulons were identified in *B. infantis* ATCC 15697 and *B. bifidum* PRL2010, both isolated from the human infant gut (**Fig. 6A)**. In *B. infantis* ATCC 15697, NagR expanded to control multiple H1 cluster genes encoding GHs and transporters involved in HMO utilization (**Fig 1B**). Interestingly, the predicted NagR operators in promoter regions of *hmoA3*, *hmoA4*, and *hmoA5* were identical (**Table S3**), reflecting a plausible scenario of multiple gene duplication events in the H1 cluster. In *B. bifidum* PRL2010, we identified putative NagR operators in promoter regions of multiple genes encoding GHs participating in extracellular HMO and mucin *O*-glycan degradation, namely lacto-*N*-biosidases LnbB (GH20) and LnbX (GH136) (33, 34), endo-α-*N*-acetylgalactosaminidase EngBF (GH101) (35), exo-β-galactosidase BbgIII (GH2) (36), exo-6-sulfo-β-*N*-acetylglucosaminidase BbhII (GH20) (37), exo-α-galactosidase AgnB (GH110) (38), and exo-α-*N*-acetylglucosaminidase AgaBb (GH89) (39) (**Fig. 6A** and **Table S3**). The reconstructed NagR regulon structure in *B. bifidum* suggests that this TF might function as a global regulator of HMO and mucin-*O*-glycan degradation in this species.

We observed strain-specific variation in NagR regulon composition in *B. infantis* and *B. longum*. So, the regulon in *B. infantis* IN-07 lacked *gltABC*, *hmoA2B2C2*, and *hmoA5* due to lineage-specific loss of these genes (**Fig. 6A**). Notably, although this strain lost the coding sequences of *gltABC,* it retained the *gltA* promoter containing a predicted NagR-binding site (**Table S3**). This observation points out the importance of the NagR-mediated regulation of the remnant *lnpABCD* genes in the *lnp* cluster. Within *B. longum*, NagR regulons contained *nag* and *lnp* clusters in both NCC2705 and JCM 1217; however, in the latter strain, the regulon also had a gene encoding lacto-*N*-biosidase (LnbX) that cleaves LNT into LNB and lactose (34) (**Fig. 6A**). Therefore, we propose that NagR might regulate not only LNB/GNB but also LNT utilization in *B. longum* JCM 1217.

All NagR protein sequences could be divided into two groups based on their cognate binding motifs (**Fig. 6B**). Both motifs had conserved "GT" and "AC" in positions 4-5 and 17-18, respectively. NagR-binding motif I had a prevalent "A" in position 3, whereas motif II had prevalent a "T" in positions 3, 14, and 15, as well as an "A" in position 7. The differences between the binding motifs were likely due to the differences in primary structures of respective TFs; for example, NagR from *B. aquikefiri* (motif I representative) and *B. infantis* ATCC 15697 (motif II representative) shared only 38 % sequence identity. Interestingly, unlike other species, *B. bohemicum* and *B. bifidum* harbored two NagR paralogs; however, their motifs could be separated only in the *B. bohemicum* case.

We also noted a discrepancy between positions of branches corresponding to *A. theropitheci* in species (**Fig. 6A)** and gene (**Fig. 6B**) phylogenetic trees. The *A. theropitheci* branch was an outgroup in the species tree, reflecting the phylogenetic differences between *Bifidobacterium* and *Alloscardovia* genera. However, in the gene tree, NagR from *A. theropitheci* clustered together with NagR sequences from bifidobacteria isolated from mammals (e.g., *Bifidobacterium tsurumiense* JCM 13495). This observation indicates a possible horizontal transfer of the NagR regulon genes from *Bifidobacterium* spp. to *A. theropitheci*.

The obtained results suggest that NagR gradually evolved from a local regulator of GlcNAc catabolism in ancestral bifidobacteria to a global regulator of various host glycan utilization in human-residential bifidobacteria via regulon expansion.

**References**


1. Overbeek R, Begley T, Butler RM, Choudhuri JV, Chuang H-Y, Cohoon M, de Crécy-Lagard V, Diaz N, Disz T, Edwards R, Fonstein M, Frank ED, Gerdes S, Glass EM, Goesmann A, Hanson A, Iwata-Reuyl D, Jensen R, Jamshidi N, Krause L, Kubal M, Larsen N, Linke B, McHardy AC, Meyer F, Neuweger H, Olsen G, Olson R, Osterman A, Portnoy V, Pusch GD, Rodionov DA, Rückert C, Steiner J, Stevens R, Thiele I, Vassieva O, Ye Y, Zagnitko O, Vonstein V. 2005. The subsystems approach to genome annotation and its use in the project to annotate 1000 genomes. Nucleic Acids Research 33:5691–5702.
2. Overbeek R, Olson R, Pusch GD, Olsen GJ, Davis JJ, Disz T, Edwards RA, Gerdes S, Parrello B, Shukla M, Vonstein V, Wattam AR, Xia F, Stevens R. 2014. The SEED and the Rapid Annotation of microbial genomes using Subsystems Technology (RAST). Nucleic Acids Res 42:D206-214.
3. Price MN, Arkin AP. 2017. PaperBLAST: Text Mining Papers for Information about Homologs. mSystems 2:e00039-17.
4. Lombard V, Golaconda Ramulu H, Drula E, Coutinho PM, Henrissat B. 2014. The carbohydrate-active enzymes database (CAZy) in 2013. Nucleic Acids Res 42:D490–D495.
5. Saier MH, Reddy VS, Moreno-Hagelsieb G, Hendargo KJ, Zhang Y, Iddamsetty V, Lam KJK, Tian N, Russum S, Wang J, Medrano-Soto A. 2021. The Transporter Classification Database (TCDB): 2021 update. Nucleic Acids Res 49:D461–D467.

6. Novichkov PS, Laikova ON, Novichkova ES, Gelfand MS, Arkin AP, Dubchak I, Rodionov DA. 2010. RegPrecise: a database of curated genomic inferences of transcriptional regulatory interactions in prokaryotes. Nucleic Acids Res 38:D111–D118.
7. Page AJ, Cummins CA, Hunt M, Wong VK, Reuter S, Holden MTG, Fookes M, Falush D, Keane JA, Parkhill J. 2015. Roary: rapid large-scale prokaryote pan genome analysis. Bioinformatics 31:3691–3693.
8. Altschul SF, Gish W, Miller W, Myers EW, Lipman DJ. 1990. Basic local alignment search tool. Journal of Molecular Biology 215:403–410.
9. Katoh K, Standley DM. 2013. MAFFT Multiple Sequence Alignment Software Version 7: Improvements in Performance and Usability. Mol Biol Evol 30:772–780.
10. Edgar RC. 2004. MUSCLE: multiple sequence alignment with high accuracy and high throughput. Nucleic Acids Res 32:1792–1797.
11. Snipen L, Liland KH. 2021. microseq: Basic Biological Sequence Handling (2.1.5). https://CRAN.R-project.org/package=microseq.
12. Nguyen L-T, Schmidt HA, von Haeseler A, Minh BQ. 2015. IQ-TREE: a fast and effective stochastic algorithm for estimating maximum-likelihood phylogenies. Mol Biol Evol 32:268–274.
13. Asakuma S, Hatakeyama E, Urashima T, Yoshida E, Katayama T, Yamamoto K, Kumagai H, Ashida H, Hirose J, Kitaoka M. 2011. Physiology of consumption of human milk oligosaccharides by infant gut-associated bifidobacteria. J Biol Chem 286:34583–34592.
14. Gotoh A, Katoh T, Sakanaka M, Ling Y, Yamada C, Asakuma S, Urashima T, Tomabechi Y, Katayama-Ikegami A, Kurihara S, Yamamoto K, Harata G, He F, Hirose J, Kitaoka M, Okuda S, Katayama T. 2018. Sharing of human milk oligosaccharides degradants within bifidobacterial communities in faecal cultures supplemented with Bifidobacterium bifidum. Sci Rep 8:13958.
15. Takada H, Katoh T, Katayama T. 2020. Sialylated O -Glycans from Hen Egg White Ovomucin are Decomposed by Mucin-degrading Gut Microbes. J Appl Glycosci (1999) 67:31–39.
16. Gotoh A, Nara M, Sugiyama Y, Sakanaka M, Yachi H, Kitakata A, Nakagawa A, Minami H, Okuda S, Katoh T, Katayama T, Kurihara S. 2017. Use of Gifu Anaerobic Medium for culturing 32 dominant species of human gut microbes and its evaluation based on short-chain fatty acids fermentation profiles. Bioscience, Biotechnology and Biochemistry 81:2009–2017.
17. Lenth RV, Buerkner P, Herve M, Love J, Miguez F, Riebl H, Singmann H. 2022. emmeans: Estimated Marginal Means, aka Least-Squares Means (1.7.2). https://CRAN.R-project.org/package=emmeans
18. Andrews S. 2010. FastQC: A Quality Control Tool for High Throughput Sequence Data. http://www.bioinformatics.babraham.ac.uk/projects/fastqc/.
19. Martin M. 2011. Cutadapt removes adapter sequences from high-throughput sequencing reads. 1. EMBnet.journal 17:10–12.
20. Langmead B, Salzberg SL. 2012. Fast gapped-read alignment with Bowtie 2. Nat Methods 9:357–359.
21. Bray NL, Pimentel H, Melsted P, Pachter L. 2016. Near-optimal probabilistic RNA-seq quantification. Nat Biotechnol 34:525–527.
22. Huber W, Carey VJ, Gentleman R, Anders S, Carlson M, Carvalho BS, Bravo HC, Davis S, Gatto L, Girke T, Gottardo R, Hahne F, Hansen KD, Irizarry RA, Lawrence M, Love MI,

MacDonald J, Obenchain V, Oleś AK, Pagès H, Reyes A, Shannon P, Smyth GK, Tenenbaum D, Waldron L, Morgan M. 2015. Orchestrating high-throughput genomic analysis with Bioconductor. Nat Methods 12:115–121.

23. Soneson C, Love MI, Robinson MD. 2015. Differential analyses for RNA-seq: transcript-level estimates improve gene-level inferences. F1000Res 4:1521.
24. Robinson MD, McCarthy DJ, Smyth GK. 2010. edgeR: a Bioconductor package for differential expression analysis of digital gene expression data. Bioinformatics 26:139–140.
25. Ritchie ME, Phipson B, Wu D, Hu Y, Law CW, Shi W, Smyth GK. 2015. limma powers differential expression analyses for RNA-sequencing and microarray studies. Nucleic Acids Res 43:e47.
26. Benjamini Y, Hochberg Y. 1995. Controlling the False Discovery Rate: A Practical and Powerful Approach to Multiple Testing. Journal of the Royal Statistical Society Series B (Methodological) 57:289–300.
27. Wickham, Hadley. 2016. ggplot2: Elegant Graphics for Data Analysis. Springer-Verlag, New York. https://ggplot2.tidyverse.org.
28. Kolde R. 2019. pheatmap: Pretty Heatmaps (1.0.12). https://CRAN.R-project.org/package=pheatmap.
29. Garrido D, Ruiz-Moyano S, Mills DA. 2012. Release and utilization of N-acetyl-d-glucosamine from human milk oligosaccharides by Bifidobacterium longum subsp. infantis. Anaerobe 18:430–435.
30. Viborg AH, Katayama T, Abou Hachem M, Andersen MCF, Nishimoto M, Clausen MH, Urashima T, Svensson B, Kitaoka M. 2014. Distinct substrate specificities of three glycoside hydrolase family 42 β-galactosidases from Bifidobacterium longum subsp. infantis ATCC 15697. Glycobiology 24:208–216.
31. Yoshida E, Sakurama H, Kiyohara M, Nakajima M, Kitaoka M, Ashida H, Hirose J, Katayama T, Yamamoto K, Kumagai H. 2012. Bifidobacterium longum subsp. infantis uses two different β-galactosidases for selectively degrading type-1 and type-2 human milk oligosaccharides. Glycobiology 22:361–368.
32. James K, Motherway MO, Bottacini F, van Sinderen D. 2016. Bifidobacterium breve UCC2003 metabolises the human milk oligosaccharides lacto-N-tetraose and lacto-N-neo-tetraose through overlapping, yet distinct pathways. Sci Rep 6.
33. Wada J, Ando T, Kiyohara M, Ashida H, Kitaoka M, Yamaguchi M, Kumagai H, Katayama T, Yamamoto K. 2008. Bifidobacterium bifidum lacto-N-biosidase, a critical enzyme for the degradation of human milk oligosaccharides with a type 1 structure. Appl Environ Microbiol 74:3996–4004.
34. Sakurama H, Kiyohara M, Wada J, Honda Y, Yamaguchi M, Fukiya S, Yokota A, Ashida H, Kumagai H, Kitaoka M, Yamamoto K, Katayama T. 2013. Lacto-N-biosidase encoded by a novel gene of Bifidobacterium longum subspecies longum shows unique substrate specificity and requires a designated chaperone for its active expression. J Biol Chem 288:25194–25206.
35. Fujita K, Oura F, Nagamine N, Katayama T, Hiratake J, Sakata K, Kumagai H, Yamamoto K. 2005. Identification and molecular cloning of a novel glycoside hydrolase family of core 1 type O-glycan-specific endo-alpha-N-acetylgalactosaminidase from Bifidobacterium longum. J Biol Chem 280:37415–37422.

36. Goulas T, Goulas A, Tzortzis G, Gibson GR. 2009. Comparative analysis of four beta-galactosidases from Bifidobacterium bifidum NCIMB41171: purification and biochemical characterisation. Appl Microbiol Biotechnol 82:1079–1088.
37. Katoh T, Maeshibu T, Kikkawa K-I, Gotoh A, Tomabechi Y, Nakamura M, Liao W-H, Yamaguchi M, Ashida H, Yamamoto K, Katayama T. 2017. Identification and characterization of a sulfoglycosidase from Bifidobacterium bifidum implicated in mucin glycan utilization. Biosci Biotechnol Biochem 81:2018–2027.
38. Wakinaka T, Kiyohara M, Kurihara S, Hirata A, Chaiwangsri T, Ohnuma T, Fukamizo T, Katayama T, Ashida H, Yamamoto K. 2013. Bifidobacterial α-galactosidase with unique carbohydrate-binding module specifically acts on blood group B antigen. Glycobiology 23:232–240.
39. Shimada Y, Watanabe Y, Wakinaka T, Funeno Y, Kubota M, Chaiwangsri T, Kurihara S, Yamamoto K, Katayama T, Ashida H. 2015. α-N-Acetylglucosaminidase from Bifidobacterium bifidum specifically hydrolyzes α-linked N-acetylglucosamine at nonreducing terminus of O-glycan on gastric mucin. Appl Microbiol Biotechnol 99:3941–3948.
