## Supplementary Code File 1 for "Human Milk Oligosaccharide Utilization in Intestinal Bifidobacteria is Governed by a Global Transcriptional Regulator NagR"

### Supplemental code file for manuscript titled: Human Milk Oligosaccharide Utilization in Intestinal Bifidobacteria is Governed by a Global Transcriptional Regulator NagR

Aleksandr A. Arzamasov   Aruto Nakajima   Mikiyasu Sakanaka   Miriam Ojima
Takane Katayama   Dmitry A. Rodionov   Andrei L. Osterman

#### Contents

**1 Background** **2**

**2 Reproducibility and accessibility** **2**

**3 R packages and external functions used** **3**

**4 Analysis of growth, HMO consumption, and organic acid production data** **3**

**8 Session info** **34**

### 1 Background

*Bifidobacterium longum* subsp. *infantis* (*B. infantis*) is a prevalent beneficial bacterium that colonizes the human neonatal gut and is uniquely adapted to efficiently use human milk oligosaccharides (HMOs) as a carbon and energy source. Multiple studies have focused on characterizing the elements of HMO utilization machinery in *B. infantis*; however, the regulatory mechanisms governing the expression of these catabolic pathways remain poorly understood. A bioinformatic regulon reconstruction approach used in this study implicated NagR, a transcription factor from the ROK family, as a negative global regulator of genomic loci encoding lacto-*N*-biose/galacto-*N*-biose (LNB/GNB), lacto-*N*-tetraose (LNT), and lacto-*N*-neotetraose (LNnT) utilization pathways in *B. infantis*. This conjecture was corroborated by transcriptome profiling upon *nagR* genetic inactivation and experimental assessment of binding of recombinant NagR to predicted DNA operators. The latter approach also implicated N-acetylglucosamine (GlcNAc), a universal intermediate of LNT and LNnT catabolism, and its phosphorylated derivatives as plausible NagR effectors. Reconstruction of NagR regulons in various *Bifidobacterium* lineages revealed multiple regulon expansion events, suggesting evolution from a local regulator of GlcNAc catabolism in ancestral bifidobacteria to a global regulator controlling foraging of mixtures of GlcNAc-containing host-derived glycans in infant gut-colonizing *B. infantis* and *Bifidobacterium bifidum*.

**This supplementary code file describes**:

1. Analysis of growth, HMO consumption, and organic acid production data
2. Analysis of RNA-seq data: (a) processing raw fastq files and mapping reads to the *Bifidobacterium longum* subsp. *infantis* ATCC 15697 transcriptome, (b) identification of differentially expressed genes (DEGs) between experimental conditions
3. Analysis of Electrophoretic Mobility Shift Assay (EMSA) data: processing gel quantification data and building 4-parameter logistic (4PL) models to infer $EC_{50}$ values
4. Phylogenetic analysis

---

### 2 Reproducibility and accessibility

All code used in this analysis, including the Rmarkdown document used to compile this supplementary code file, is available on GitHub **here**. Once the GitHub repo has been downloaded, navigate to `NagR_manuscript/` to find the Rmarkdown document as well as the RProject file. This should be your working directory for executing code. To fully reproduce the processing of raw fastq files (RNA-seq data analysis), you will need to download them from Gene Expression Omnibus, under accession **GSE196064**. Downloaded fastq files should be placed to `NagR_manuscript/data/rnaseq/fastq/`. Otherwise, `data/rnaseq/kallisto/` already includes Kallisto mapping outputs.

---

### 3 R packages and external functions used

A set of R packages was used for this analysis. The pacman package was used to simplify downloading and loading the required packages. All graphics and data wrangling were handled using the tidyverse suite of packages. To fully reproduce the R environment used in the analysis, use the renv package and `renv::restore()` to restore the environment from `renv.lock`.

```
# install/load the pacman package for rapid installation of packages that are not in library
if (!require("pacman")) install.packages("pacman")
# use pacman to install/load all packages needed for the analysis
pacman::p_load('tidyverse', 'emmeans', 'tximport', 'gt', 'tidyverse', 'edgeR', 'matrixStats',
               'cowplot', 'ggrepel', 'pheatmap', 'drc', 'Cairo', 'ggpubr', 'microseq')
```

A set of external R functions was used to keep the code tidy. All used R scripts with functions can be found in `NagR_manuscript/code/`.

```
source("code/profile.R") # calculates counts per million (CPM) for each gene
# plots the distribution of CPM values for each sample
source("code/deg_list.R") # selects DEGs based on input cut-offs
# outputs an annotated table with DEGs to a txt file
source("code/emsa.R") # builds a 4PL model for EMSA quantification data
# plots the fit from the 4PL model with 95% confidence intervals
```

---

### 4 Analysis of growth, HMO consumption, and organic acid production data

#### 4.1 Growth data Fig. 2

Growth of *Bifidobacterium longum* subsp. *infantis* ATCC 15697 WT and *nagR*-KO strains in MRS-CS supplemented with various carbon sources was measured by $OD_{600}$. Data points (taken at 0, 4, 8, 12, and 24 h) represent the mean of three replicates; error bars depict 95% confidence intervals for the mean.

```
# read growth data
growth_data <- read_tsv("data/growth/growth_data.txt") %>%
  filter(sugar != "control")
# plot data
set.seed(123)
growth_data %>%
  # rearrange plot order
  mutate(across(sugar, factor, levels=c("Lactose", "HMO", "Sucrose", "FOS"))) %>%
ggplot(aes(x = time, y = od, group = strain)) +
  # display mean and connect points by line
  stat_summary(
    fun = mean,
    geom='line',
    size = 0.7,
    aes(color=strain)) +
  # display 95% non-parametric confidence interval for the mean
```

```
stat_summary(
  fun.data=mean_cl_boot,
  geom='errorbar',
  width=0.6,
  aes(color=strain)) +
stat_summary(
  fun=mean,
  geom ='point',
  aes(fill=strain, shape=strain),
  size = 1.5)+
scale_color_manual(name="Strain",
                   values=c(WT="#008800", Mut="#F48326")) +
scale_fill_manual(name="Strain",
                   values=c(WT="#008800", Mut="#F48326")) +
scale_shape_manual(name="Strain",
                   values=c(WT=21, Mut=24)) +
scale_x_continuous(limits=c(-0.4, 24.4),
                    breaks=c(0, 4, 8, 12, 16, 20, 24)) +
scale_y_continuous(limits=c(0, 6.5),
                    breaks=c(0, 1, 2, 3, 4, 5, 6)) +
xlab("Time (h)") +
ylab("OD600") +
facet_wrap(. ~ sugar) +
theme_bw(14) +
theme(axis.text=element_text(size=10), strip.text = element_text(size = 14))
```

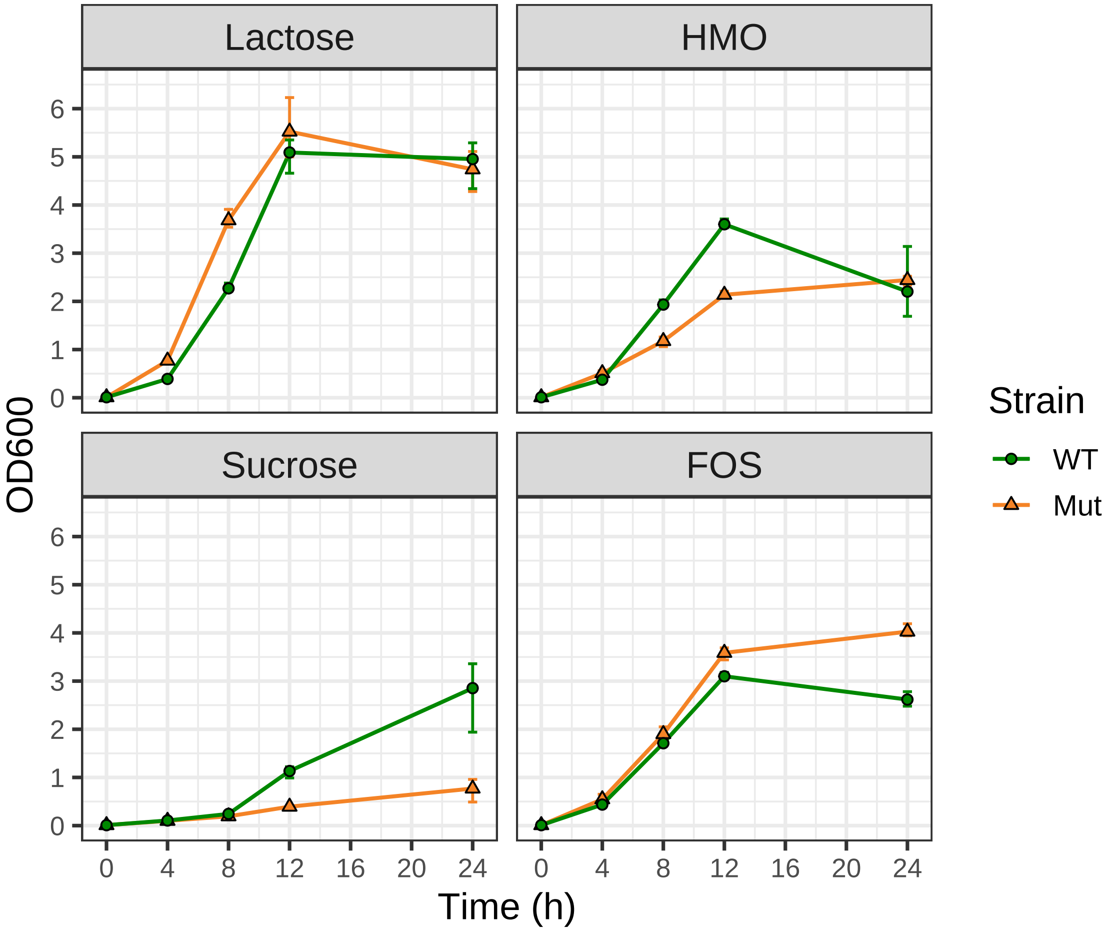

```
# save the figure
ggsave("results/figures/figure_2.pdf", device = cairo_pdf, width = 6, height = 5)
```

A linear model and post hoc comparisons implemented in the emmeans package were used to identify time points where the mean $OD_{600}$ values for WT and *nagR*-KO strains were significantly different. Computed p-values were adjusted for multiple comparisons using the Bonferroni correction.

```
# function below:
# (1) builds a linear model with two factors (strain and time)
# (2) performs pairwise comparisons of OD600 means (WT vs nagR-KO) at each time point using emmeans
# (3) computes p-values and adjusts for multiple comparisons using the Bonferroni correction
compare_growth_means <- function(measurement_data){
  lm_strain <- lm(od ~ strain + time + strain:time, data=measurement_data)
  compare_means_strain <- emmeans(lm_strain, ~ strain | time)
  summary(pairs(compare_means_strain), by = NULL, adjust = "bonferroni")
}

# for each carbon source, compare OD600 values (WT vs nagR-KO) at each time point
growth_data_split <- growth_data %>%
  mutate(across(c(strain, time), factor)) %>%
  group_by(sugar) %>%
  split(f = as.factor(.$sugar))
lapply(growth_data_split, compare_growth_means)
```

```
## $FOS
##  contrast time estimate     SE df t.ratio p.value
##  Mut - WT 0       0.000 0.0908 20   0.000  1.0000
##  Mut - WT 4       0.110 0.0908 20   1.212  1.0000
##  Mut - WT 8       0.190 0.0908 20   2.093  0.2463
##  Mut - WT 12      0.487 0.0908 20   5.362  0.0002
##  Mut - WT 24      1.410 0.0908 20  15.535  <.0001
##
## P value adjustment: bonferroni method for 5 tests
##
## $HMO
##  contrast time estimate    SE df t.ratio p.value
##  Mut - WT 0       0.000 0.217 20   0.000  1.0000
##  Mut - WT 4       0.137 0.217 20   0.630  1.0000
##  Mut - WT 8      -0.757 0.217 20  -3.489  0.0116
##  Mut - WT 12     -1.463 0.217 20  -6.748  <.0001
##  Mut - WT 24      0.240 0.217 20   1.107  1.0000
##
## P value adjustment: bonferroni method for 5 tests
##
## $Lactose
##  contrast time estimate    SE df t.ratio p.value
##  Mut - WT 0       0.000 0.266 20   0.000  1.0000
##  Mut - WT 4       0.380 0.266 20   1.430  0.8403
##  Mut - WT 8       1.413 0.266 20   5.320  0.0002
##  Mut - WT 12      0.433 0.266 20   1.631  0.5926
##  Mut - WT 24     -0.213 0.266 20  -0.803  1.0000
##
## P value adjustment: bonferroni method for 5 tests
##
## $Sucrose
##  contrast time estimate    SE df t.ratio p.value
##  Mut - WT 0     0.00000 0.218 20   0.000  1.0000
##  Mut - WT 4    -0.00667 0.218 20  -0.031  1.0000
##  Mut - WT 8    -0.05000 0.218 20  -0.230  1.0000
##  Mut - WT 12   -0.74000 0.218 20  -3.401  0.0142
##  Mut - WT 24   -2.08333 0.218 20  -9.576  <.0001
##
## P value adjustment: bonferroni method for 5 tests
```

---

#### 4.2 HMO consumption Fig. S2C

HMO consumption of *Bifidobacterium longum* subsp. *infantis* ATCC 15697 WT and *nagR*-KO strains grown in MRS-CS-HMO was measured by normal phase high-performance liquid chromatography (HPLC). Data points (taken at 0, 4, 8, 12, and 24 h) represent the mean of three replicates; error bars depict 95% confidence intervals for the mean.

```r
# read HMO consumption data
hmo_data <- read_tsv("data/metabolomics/HMO_concentration.txt")
# plot data
set.seed(123)
hmo_data %>%
  # rearrange plot order
  mutate(across(sugar, factor, levels=c("Lactose", "2'FL", "3FL", "DFL",
                                        "LNT", "LNnT", "LNFP I",
                                        "LNFP II/III", "LNDFH I",
                                        "LNDFH II",
                                        "LNTriose II", "Fucose"))) %>%
  ggplot(aes(x = time, y = conc, group = strain)) +
  # display mean and connect points by line
  stat_summary(
    fun = mean,
    geom='line',
    size = 1,
    aes(color=strain)) +
  # display 95% non-parametric confidence intervals for the mean
  stat_summary(
    fun.data=mean_cl_boot,
    geom='errorbar',
    width=0.6,
    aes(color=strain)) +
  stat_summary(
    fun=mean,
    geom ='point',
    aes(fill=strain, shape=strain),
    size = 3) +
  scale_color_manual(name="Strain",
                     values=c(WT="#008800", Mut="#F48326")) +
  scale_fill_manual(name="Strain",
                    values=c(WT="#008800", Mut="#F48326")) +
  scale_shape_manual(name="Strain",
                     values=c(WT=21, Mut=24)) +
  scale_x_continuous(limits=c(-0.4, 24.4),
                     breaks=c(0, 4, 8, 12, 16, 20, 24)) +
  xlab("Time (h)") +
  ylab("Concentration (mM)") +
  # set different scales for each plot
  facet_wrap(. ~ sugar, scales = "free") +
  theme_bw(14)
```

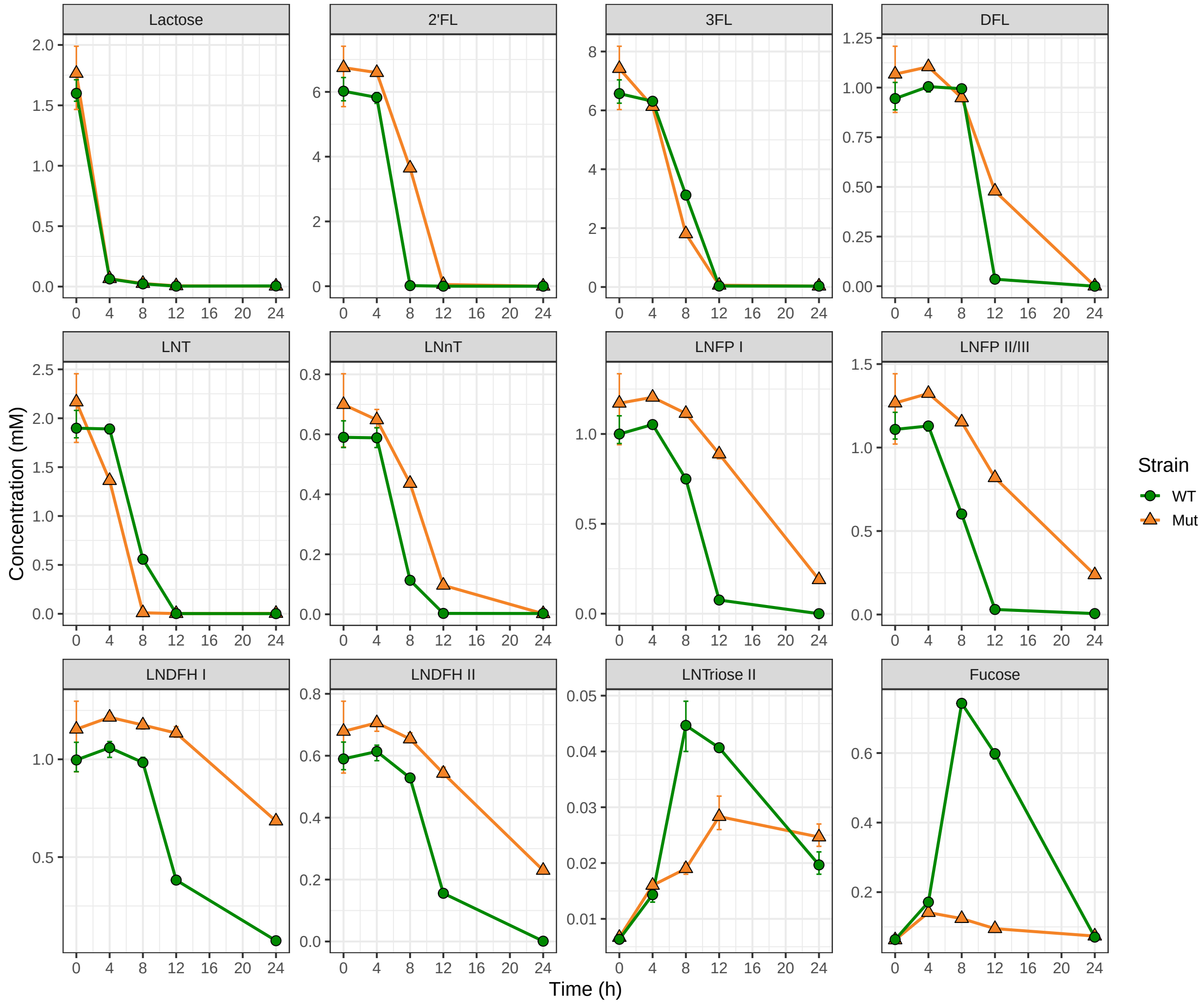

```r
# save the figure
ggsave("results/figures/figure_S2C.pdf", device = cairo_pdf, width = 12, height = 10)
```

A linear model and post hoc comparisons implemented in the emmeans package were used to identify time points where the mean HMO concentrations for WT and *nagR*-KO strains were significantly different. Computed p-values were adjusted for multiple comparisons using the Bonferroni correction.

```r
# function below:
# (1) builds a linear model with two factors (strain and time)
# (2) performs pairwise comparisons of concentration means (WT vs nagR-KO) at each time point
# using emmeans
# (3) compute p-values and adjusts for multiple comparisons using Bonferroni correction
compare_growth_means <- function(measurement_data){
  lm_strain <- lm(conc ~ strain + time + strain:time, data=measurement_data)
  compare_means_strain <- emmeans(lm_strain, ~ strain | time)
  summary(pairs(compare_means_strain), by = NULL, adjust = "bonferroni")
}

# for each carbon source, compare OD600 values (WT vs nagR-KO) at each time point
hmo_data_split <- hmo_data %>%
```

```
  mutate(across(c(strain, time), factor)) %>%
  group_by(sugar) %>%
  split(f = as.factor(.$sugar))
lapply(hmo_data_split, compare_growth_means)
```

```
## $`2'FL`
##  contrast time estimate    SE df t.ratio p.value
##  Mut - WT 0      0.7273 0.293 20   2.484  0.1099
##  Mut - WT 4      0.7660 0.293 20   2.616  0.0828
##  Mut - WT 8      3.6280 0.293 20  12.389  <.0001
##  Mut - WT 12     0.0517 0.293 20   0.176  1.0000
##  Mut - WT 24     0.0000 0.293 20   0.000  1.0000
##
## P value adjustment: bonferroni method for 5 tests
##
## $`3FL`
##  contrast time estimate    SE df t.ratio p.value
##  Mut - WT 0      0.8510 0.334 20   2.548  0.0958
##  Mut - WT 4     -0.1780 0.334 20  -0.533  1.0000
##  Mut - WT 8     -1.3160 0.334 20  -3.941  0.0040
##  Mut - WT 12     0.0273 0.334 20   0.082  1.0000
##  Mut - WT 24     0.0000 0.334 20   0.000  1.0000
##
## P value adjustment: bonferroni method for 5 tests
##
## $DFL
##  contrast time  estimate     SE df t.ratio p.value
##  Mut - WT 0     0.123000 0.0491 20   2.506  0.1048
##  Mut - WT 4     0.099667 0.0491 20   2.031  0.2790
##  Mut - WT 8    -0.046000 0.0491 20  -0.937  1.0000
##  Mut - WT 12    0.443333 0.0491 20   9.032  <.0001
##  Mut - WT 24    0.000667 0.0491 20   0.014  1.0000
##
## P value adjustment: bonferroni method for 5 tests
##
## $Fucose
##  contrast time estimate      SE df  t.ratio p.value
##  Mut - WT 0      -0.001 0.00439 20   -0.228  1.0000
##  Mut - WT 4      -0.030 0.00439 20   -6.831  <.0001
##  Mut - WT 8      -0.619 0.00439 20 -140.941  <.0001
##  Mut - WT 12     -0.503 0.00439 20 -114.605  <.0001
##  Mut - WT 24      0.003 0.00439 20    0.683  1.0000
##
## P value adjustment: bonferroni method for 5 tests
##
## $Lactose
##  contrast time estimate    SE df t.ratio p.value
##  Mut - WT 0     0.16500 0.074 20   2.229  0.1874
##  Mut - WT 4     0.00200 0.074 20   0.027  1.0000
##  Mut - WT 8     0.00333 0.074 20   0.045  1.0000
##  Mut - WT 12    0.00233 0.074 20   0.032  1.0000
##  Mut - WT 24   -0.00200 0.074 20  -0.027  1.0000
##
```

```
## P value adjustment: bonferroni method for 5 tests
##
## $`LNDFH I`
##  contrast time estimate     SE df t.ratio p.value
##  Mut - WT 0       0.157 0.0565 20   2.783  0.0574
##  Mut - WT 4       0.156 0.0565 20   2.753  0.0613
##  Mut - WT 8       0.192 0.0565 20   3.390  0.0145
##  Mut - WT 12      0.752 0.0565 20  13.294  <.0001
##  Mut - WT 24      0.612 0.0565 20  10.824  <.0001
##
## P value adjustment: bonferroni method for 5 tests
##
## $`LNDFH II`
##  contrast time estimate     SE df t.ratio p.value
##  Mut - WT 0      0.0890 0.0355 20   2.510  0.1040
##  Mut - WT 4      0.0927 0.0355 20   2.613  0.0833
##  Mut - WT 8      0.1257 0.0355 20   3.544  0.0102
##  Mut - WT 12     0.3873 0.0355 20  10.922  <.0001
##  Mut - WT 24     0.2287 0.0355 20   6.448  <.0001
##
## P value adjustment: bonferroni method for 5 tests
##
## $`LNFP I`
##  contrast time estimate     SE df t.ratio p.value
##  Mut - WT 0       0.171 0.0591 20   2.900  0.0442
##  Mut - WT 4       0.152 0.0591 20   2.567  0.0919
##  Mut - WT 8       0.364 0.0591 20   6.168  <.0001
##  Mut - WT 12      0.813 0.0591 20  13.757  <.0001
##  Mut - WT 24      0.189 0.0591 20   3.199  0.0225
##
## P value adjustment: bonferroni method for 5 tests
##
## $`LNFP II/III`
##  contrast time estimate     SE df t.ratio p.value
##  Mut - WT 0       0.158 0.0621 20   2.546  0.0962
##  Mut - WT 4       0.195 0.0621 20   3.137  0.0260
##  Mut - WT 8       0.551 0.0621 20   8.874  <.0001
##  Mut - WT 12      0.789 0.0621 20  12.714  <.0001
##  Mut - WT 24      0.232 0.0621 20   3.739  0.0065
##
## P value adjustment: bonferroni method for 5 tests
##
## $LNnT
##  contrast time  estimate    SE df t.ratio p.value
##  Mut - WT 0     0.109333 0.037 20   2.951  0.0395
##  Mut - WT 4     0.060000 0.037 20   1.620  0.6048
##  Mut - WT 8     0.323000 0.037 20   8.719  <.0001
##  Mut - WT 12    0.093667 0.037 20   2.528  0.0999
##  Mut - WT 24   -0.000667 0.037 20  -0.018  1.0000
##
## P value adjustment: bonferroni method for 5 tests
##
## $LNT
##  contrast time estimate    SE df t.ratio p.value
```

```
##  Mut - WT 0       0.268 0.104 20   2.572  0.0909
##  Mut - WT 4      -0.528 0.104 20  -5.068  0.0003
##  Mut - WT 8      -0.547 0.104 20  -5.253  0.0002
##  Mut - WT 12      0.000 0.104 20   0.000  1.0000
##  Mut - WT 24      0.002 0.104 20   0.019  1.0000
##
## P value adjustment: bonferroni method for 5 tests
##
## $`LNTriose II`
##  contrast time  estimate      SE df t.ratio p.value
##  Mut - WT 0     0.000333 0.00171 20   0.195  1.0000
##  Mut - WT 4     0.001667 0.00171 20   0.977  1.0000
##  Mut - WT 8    -0.025667 0.00171 20 -15.043  <.0001
##  Mut - WT 12   -0.012333 0.00171 20  -7.229  <.0001
##  Mut - WT 24    0.005000 0.00171 20   2.930  0.0414
##
## P value adjustment: bonferroni method for 5 tests
```

---

#### 4.3 Organic acid production Fig. S2D

Organic acid production of *Bifidobacterium longum* subsp. *infantis* ATCC 15697 WT and *nagR*-KO strains grown in MRS-CS-HMO was measured by anion-exchange chromatography. Since MRS-CS contains baseline levels of acetate, formate, and lactate, measurements at 0 h were subtracted from subsequent time points (4, 8, 12, and 24 h). Data represent the mean of three replicates; error bars depict 95% confidence intervals for the mean.

```
# read SCFA production data
scfa_data <- read_tsv("data/metabolomics/organic_acid_concentration.txt")
# plot data
set.seed(123)
scfa_data %>%
  # rearrange plot order
  mutate(across(scfa, factor, levels=c("Acetate", "Formate", "Lactate"))) %>%
  ggplot(aes(x = time, y = conc_substr, group = strain)) +
  # display mean and connect points by line
  stat_summary(
    fun = mean,
    geom='line',
    size = 1,
    aes(color=strain)) +
  # display 95% non-parametric confidence intervals for the mean
  stat_summary(
    fun.data=mean_cl_boot,
    geom='errorbar',
    width=0.6,
    aes(color=strain)) +
  stat_summary(
    fun=mean,
    geom ='point',
    aes(fill=strain, shape=strain),
    size = 3) +
```

```
  scale_color_manual(name="Strain",
                     values=c(WT="#008800", Mut="#F48326")) +
  scale_fill_manual(name="Strain",
                     values=c(WT="#008800", Mut="#F48326")) +
  scale_shape_manual(name="Strain",
                     values=c(WT=21, Mut=24)) +
  scale_x_continuous(limits=c(-0.4, 24.4),
                     breaks=c(0, 4, 8, 12, 16, 20, 24)) +
  xlab("Time (h)") +
  ylab("Concentration (mM)") +
  facet_wrap(. ~ scfa, scales = "free") +
  theme_bw(14)
```

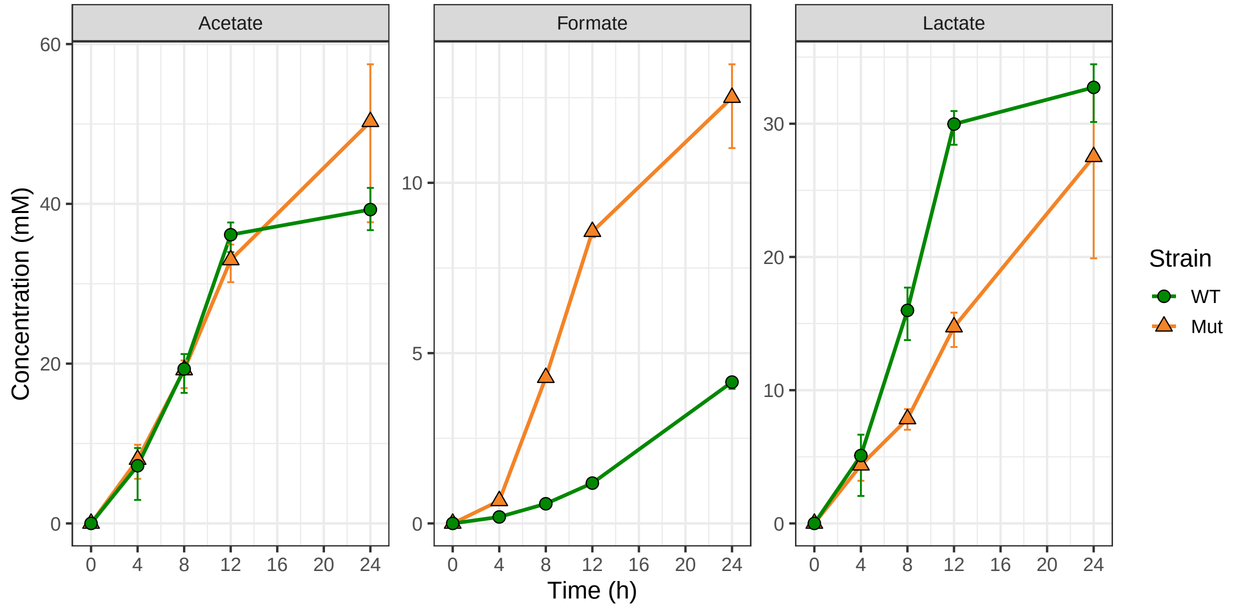

```
# save the figure
ggsave("results/figures/figure_S2D.pdf", device = cairo_pdf, width = 10, height = 5)
```

A linear model and post hoc comparisons implemented in the emmeans package were used to identify time points where the mean organic acid concentrations for WT and *nagR*-KO strains were significantly different. Computed p-values were adjusted for multiple comparisons using the Bonferroni correction.

```
# function below:
# (1) builds a linear model with two factors (strain and time)
# (2) performs pairwise comparisons of concentration means (WT vs nagR-KO) at each time point
# using emmeans
# (3) compute p-values and adjusts for multiple comparisons using Bonferroni correction
compare_growth_means <- function(measurement_data){
  lm_strain <- lm(conc ~ strain + time + strain:time, data=measurement_data)
  compare_means_strain <- emmeans(lm_strain, ~ strain | time)
  summary(pairs(compare_means_strain), by = NULL, adjust = "bonferroni")
}
```

```r
# for each carbon source, compare OD600 values (WT vs nagR-KO) at each time point
scfa_data_split <- scfa_data %>%
  mutate(across(c(strain, time), factor)) %>%
  group_by(scfa) %>%
  split(f = as.factor(.$scfa))
lapply(scfa_data_split, compare_growth_means)
```

```
## $Acetate
##  contrast time estimate   SE df t.ratio p.value
##  Mut - WT 0     -0.8400 3.32 20  -0.253  1.0000
##  Mut - WT 4     -0.0567 3.32 20  -0.017  1.0000
##  Mut - WT 8     -0.9633 3.32 20  -0.290  1.0000
##  Mut - WT 12    -3.9933 3.32 20  -1.204  1.0000
##  Mut - WT 24    10.1567 3.32 20   3.062  0.0308
##
## P value adjustment: bonferroni method for 5 tests
##
## $Formate
##  contrast time estimate    SE df t.ratio p.value
##  Mut - WT 0     -0.0367 0.342 20  -0.107  1.0000
##  Mut - WT 4      0.4333 0.342 20   1.269  1.0000
##  Mut - WT 8      3.6667 0.342 20  10.736  <.0001
##  Mut - WT 12     7.3400 0.342 20  21.491  <.0001
##  Mut - WT 24     8.3100 0.342 20  24.332  <.0001
##
## P value adjustment: bonferroni method for 5 tests
##
## $Lactate
##  contrast time estimate   SE df t.ratio p.value
##  Mut - WT 0      -0.983 2.12 20  -0.464  1.0000
##  Mut - WT 4      -1.713 2.12 20  -0.808  1.0000
##  Mut - WT 8      -9.130 2.12 20  -4.306  0.0017
##  Mut - WT 12    -16.220 2.12 20  -7.649  <.0001
##  Mut - WT 24     -6.190 2.12 20  -2.919  0.0424
##
## P value adjustment: bonferroni method for 5 tests
```

---

### 5 Analysis of RNA-seq data

#### 5.1 Processing of raw fastq files and read mapping (Optional)

The code chuck below describes processing of raw fastq files and mapping reads to the *Bifidobacterium longum* subsp. *infantis* ATCC 15697 transcriptome. The following software is required (can be installed to one conda environment):

1. FastQC(v0.11.9)
2. Cutadapt (v3.4)
3. Bowtie2 (v 2.4.4)
4. Kallisto (v0.46.2)

5. MultiQC (v1.11)
6. Parallel (v20210222)

**Note**: due to size limitation, raw fastq files could not be stored in the GitHub repo. Thus, you will need to download the fastq files from the Gene Expression Omnibus, under accession GSE196064. Put downloaded fastq.gz files to `data/rnaseq/fastq/`.

The reference fasta files used for building Bowtie2 and Kallisto indeces can be found in `data/rnaseq/refs/`.

**Note**: exact file names (without the .fastq.qz extension) should be entered into `data/rnaseq/runids.txt`.

Alternatively, you can run `code/qc_readmapping.sh` instead of the code chunk below.

Summary of the script:

1. Quality control of raw reads was carried out using FastQC
2. Illumina sequencing adapters and short reads (< 20 bp) were removed using Cutadapt
3. Reads were mapped against rRNA and tRNA gene sequences extracted from the *Bifidobacterium longum* subsp. *infantis* ATCC 15697 genome (GenBank accession no. CP001095.1) using Bowtie2. Unmapped (filtered) reads were saved and used further
4. Filtered reads were mapped to the *Bifidobacterium longum* subsp. *infantis* ATCC 15697 (CP001095.1) using Kallisto
5. The quality of raw/filtered reads, as well as the results of Bowtie2/Kallisto mapping were summarized in `data/rnaseq/multiqc_report.html` generated via MultiQC

```
source ~/.bash_profile
echo $BASH_VERSION
set -ex

# required software: fastqc (v0.11.9), cutadapt (v3.4), bowtie2 (v2.4.4), kallisto (v0.46.2)
# multiqc (v1.11), and parallel (v20210222)
# sample names should be in data/rnaseq/runids.txt

# activate conda enviroment with required software
eval "$(command conda 'shell.bash' 'hook' 2> /dev/null)" # initializes conda in sub-shell
conda activate transcriptomics
conda info|egrep "conda version|active environment"

# create directories
mkdir data/rnaseq/qc1 # qc results for raw reads
mkdir data/rnaseq/qc2 # qc results for filtered reads
mkdir data/rnaseq/fq_trim # trimmed reads
mkdir data/rnaseq/fq_filt # filtered reads
mkdir data/rnaseq/sam # sam files produced during bowtie2 alignment; will be deleted
mkdir data/rnaseq/kallisto # kallisto mapping results

# run fastqc on raw reads
cat data/rnaseq/runids.txt | parallel "fastqc data/rnaseq/fastq/{}.fastq.gz \
--outdir data/rnaseq/qc1"

# trim adapters using cutadapt
cat data/rnaseq/runids.txt | parallel "cutadapt -m 20 -a AGATCGGAAGAGCACACGTCTGAACTCCAGTC \
-o data/rnaseq/fq_trim/{}.fastq.gz data/rnaseq/fastq/{}.fastq.gz \
&> data/rnaseq/fq_trim/{}.fastq.qz.log"
```

```
# filter reads mapping to rRNA and tRNA genes
# build bowtie2 index
bowtie2-build data/rnaseq/refs/Binfantis_ATCC15697_rRNA_tRNA.fasta \
data/rnaseq/refs/Binfantis_ATCC15697_rRNA_tRNA
# align reads via bowtie2; save ones that did not align to a separate file
cat data/rnaseq/runids.txt | \
parallel "bowtie2 -x data/rnaseq/refs/Binfantis_ATCC15697_rRNA_tRNA \
-U data/rnaseq/fq_trim/{}.fastq.gz \
-S data/rnaseq/sam/{}.sam \
--un data/rnaseq/fq_filt/{}.fastq \
&> data/rnaseq/fq_filt/{}.log"
cat data/rnaseq/runids.txt | parallel "gzip data/rnaseq/fq_filt/{}.fastq"

# run fastqc on filtered reads
cat data/rnaseq/runids.txt | parallel "fastqc data/rnaseq/fq_filt/{}.fastq.gz \
--outdir data/rnaseq/qc2"

# pseudolalign reads to transcriptome
# build kallisto index
kallisto index -i data/rnaseq/refs/Binfantis_ATCC15697_transcriptome.index \
data/rnaseq/refs/Binfantis_ATCC15697_transcriptome.fasta
# map reads to indexed reference via kallisto
cat data/rnaseq/runids.txt | parallel "kallisto quant \
-i data/rnaseq/refs/Binfantis_ATCC15697_transcriptome.index \
-o data/rnaseq/kallisto/{} \
--single \
-l 200 \
-s 20 \
data/rnaseq/fq_filt/{}.fastq.gz \
&> data/rnaseq/kallisto/{}_2.log"

# run multiqc
export LC_ALL=en_US.utf-8
export LANG=en_US.utf-8
multiqc -d . -o data/rnaseq

# remove directories with intermediate files
rm -rf data/rnaseq/qc1
rm -rf data/rnaseq/qc2
rm -rf data/rnaseq/fq_filt
rm -rf data/rnaseq/sam
```

---

#### 5.2 Importing count data into R

TxImport was used to read Kallisto output into the R environment.

*Note*: before running the code, double-check that file names in the file_names column in `data/rnaseq/studydesign.txt` are identical to file names in `data/rnaseq/runids.txt`.

```r
# read the study design file
targets <- read_tsv("data/rnaseq/studydesign.txt")
# set file paths to Kallisto output folders with quantification data
files <- file.path("data/rnaseq/kallisto", targets$file_name, "abundance.tsv")
# check that all output files are present
all(file.exists(files))
```

```
## [1] TRUE
```

```r
# use 'tximport' to import Kallisto output into R
txi_kallisto <- tximport(files,
                         type = "kallisto",
                         txOut = TRUE, # import at transcript level
                         countsFromAbundance = "lengthScaledTPM")

# capture variables of interest from the study design
condition <- as.factor(targets$condition)
condition <- factor(condition, levels = c("WT_Lac", "Mut_Lac", "WT_LNnT", "Mut_LNnT"))
batch <- as.factor(targets$batch)
strain <- as.factor(targets$strain)
carb <- as.factor(targets$carb)
# capture sample labels for later use
sampleLabels <- targets$sample

# saw a table with raw counts for GEO submission
raw_counts <- as.tibble(txi_kallisto$counts, rownames = "locus_tag")
colnames(raw_counts) <- c("geneID", sampleLabels)
write_tsv(raw_counts, "results/tables/Arzamasov_raw_count_matrix.txt")

# use gt package to produce the study design table
gt(targets) %>%
  cols_align(
    align = "left",
    columns = TRUE
  )
```

| sample | file_name | condition | batch | strain | carb |
|---|---|---|---|---|---|
| WT_Lac1 | 01_WT_Lac1_S53_merged_lanes | WT_Lac | 1 | ATCC15697 | Lac |
| Mut_Lac1 | 02_Mut_Lac1_S54_merged_lanes | Mut_Lac | 1 | M3 | Lac |
| WT_LNnT1 | 03_WT_LNnT1_S55_merged_lanes | WT_LNnT | 2 | ATCC15697 | LNnT |
| Mut_LNnT1 | 04_Mut_LNnT1_S56_merged_lanes | Mut_LNnT | 2 | M3 | LNnT |
| WT_Lac2 | 05_WT_Lac2_S57_merged_lanes | WT_Lac | 1 | ATCC15697 | Lac |
| Mut_Lac2 | 06_Mut_Lac2_S58_merged_lanes | Mut_Lac | 1 | M3 | Lac |
| WT_LNnT2 | 07_WT_LNnT2_S59_merged_lanes | WT_LNnT | 2 | ATCC15697 | LNnT |
| Mut_LNnT2 | 08_Mut_LNnT2_S60_merged_lanes | Mut_LNnT | 2 | M3 | LNnT |
| WT_Lac3 | 09_WT_Lac3_S61_merged_lanes | WT_Lac | 1 | ATCC15697 | Lac |
| Mut_Lac3 | 10_Mut_Lac3_S62_merged_lanes | Mut_Lac | 1 | M3 | Lac |
| WT_LNnT3 | 11_WT_LNnT3_S63_merged_lanes | WT_LNnT | 2 | ATCC15697 | LNnT |
| Mut_LNnT3 | 12_Mut_LNnT3_S64_merged_lanes | Mut_LNnT | 2 | M3 | LNnT |

##### 5.3 Filtering and normalization

```
myDGEList <- DGEList(txi_kallisto$counts)
# plot unfiltered, non-normalized CPM
p1 <- profile(myDGEList, sampleLabels, "Unfiltered, non-normalized")
# filter counts
cpm <- cpm(myDGEList)
keepers <- rowSums(cpm>1)>=3 # only keep genes that have cpm>1 (== not zeroes)
# in more than 2 samples (minimal group size)
myDGEList.filtered <- myDGEList[keepers,]
# plot filtered, non-normalized CPM
p2 <- profile(myDGEList.filtered, sampleLabels, "Filtered, non-normalized")
# normalize counts via the TMM method implemented in edgeR
myDGEList.filtered.norm <- calcNormFactors(myDGEList.filtered, method = "TMM")
# plot filtered, normalized CPM
p3 <- profile(myDGEList.filtered.norm, sampleLabels, "Filtered, TMM normalized")
# compare distributions of the CPM values
plot_grid(p1, p2, p3, labels = c('A', 'B', 'C'), label_size = 12)
```

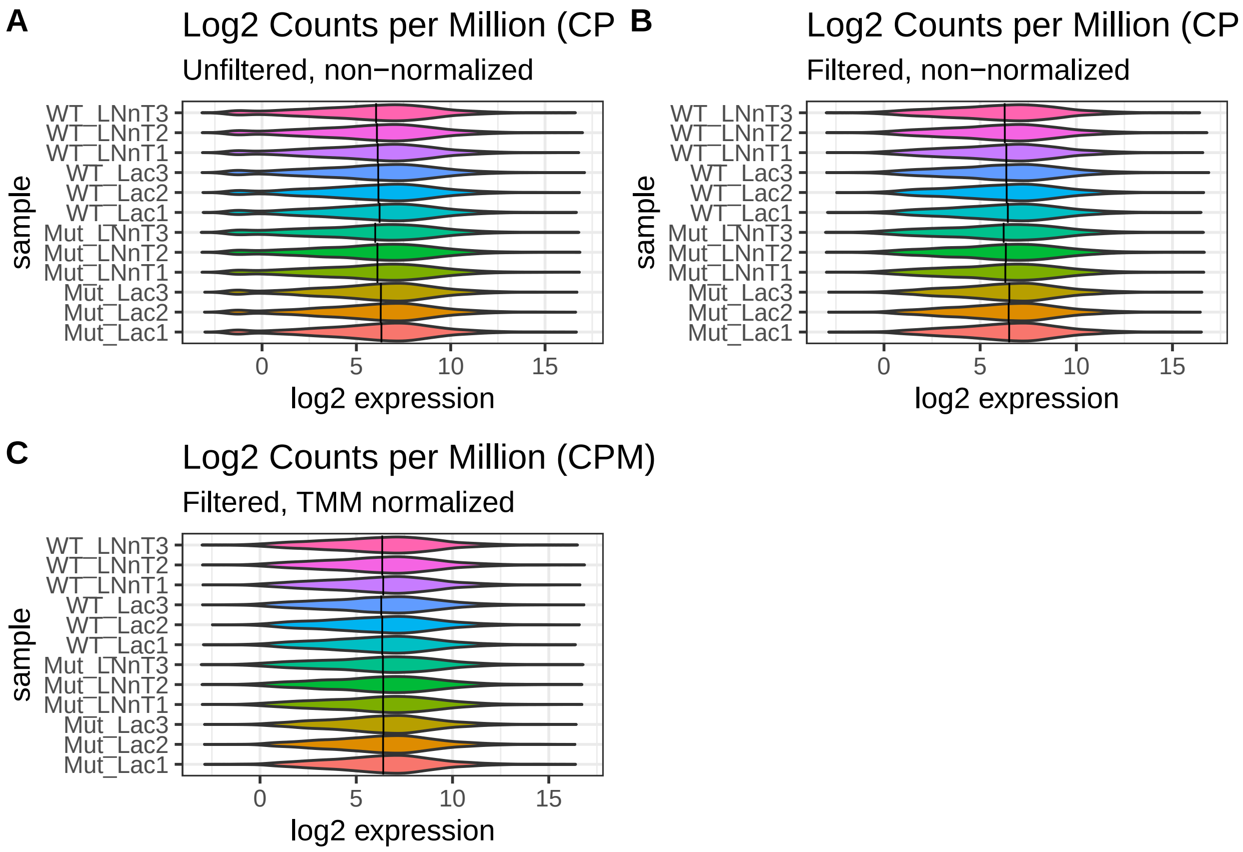

Filtering was carried out to remove lowly expressed genes. Genes with less than 1 count per million (CPM) in at least 3 or more samples were filtered out. This procedure reduced the number of genes from **2508** to **2366**. In addition, the TMM method was used for between-sample normalization.

##### 5.4 PCA plot Fig. 3B

Principal Component Analysis (PCA) plots reduce complex datasets to a 2D representation where each axis represents a source of variance (known or unknown) in the dataset. As you can see from the plots below, Principal Component 1 (PC1; X-axis), which accounts for >38% of the variance in the data, is separating the samples based on carbon source. PC2 (Y-axis) accounts for a smaller source of variance (~21%) and can be attributed to variation between strains of *Bifidobacterium longum* subsp. *infantis* ATCC 15697: WT and *nagR*-KO.

```
# running PCA
log2.cpm.filtered.norm <- cpm(myDGEList.filtered.norm, log=TRUE)
pca.res <- prcomp(t(log2.cpm.filtered.norm), scale.=F, retx=T)
pc.var <- pca.res$sdev^2 # sdev^2 captures eigenvalues from the PCA result
pc.per <- round(pc.var/sum(pc.var)*100, 1) # calculate percentage of the total variation
# explained by each eigenvalue
# converting PCA result into a tibble for plotting
pca.res.df <- as_tibble(pca.res$x)

# plotting PCA
ggplot(pca.res.df) +
  aes(x=PC1, y=PC2, label=sampleLabels, shape = carb, fill = strain) +
  geom_point(size=4) +
  scale_shape_manual(name = "Carb. source",
                     breaks=c("Lac","LNnT"),
                     values=c(21, 24),
                     labels=c("Lactose", "LNnT")) +
  scale_fill_manual(name = "Strain",
                     breaks=c("ATCC15697","M3"),
                     values=c("#008800", "#ffcf34"),
                     labels=c("WT", "nagR-KO")) +
  guides(fill = guide_legend(override.aes=list(shape=21))) +
  xlab(paste0("PC1 (",pc.per[1],"%",")")) +
  ylab(paste0("PC2 (",pc.per[2],"%",")")) +
  labs(title= "PCA of B. infantis ATCC 15697: WT vs nagR-KO",
       color = "strain", shape="carb") +
  coord_fixed(ratio=1.2) +
  theme_bw() +
  theme(plot.title = element_text(face="bold"))
```

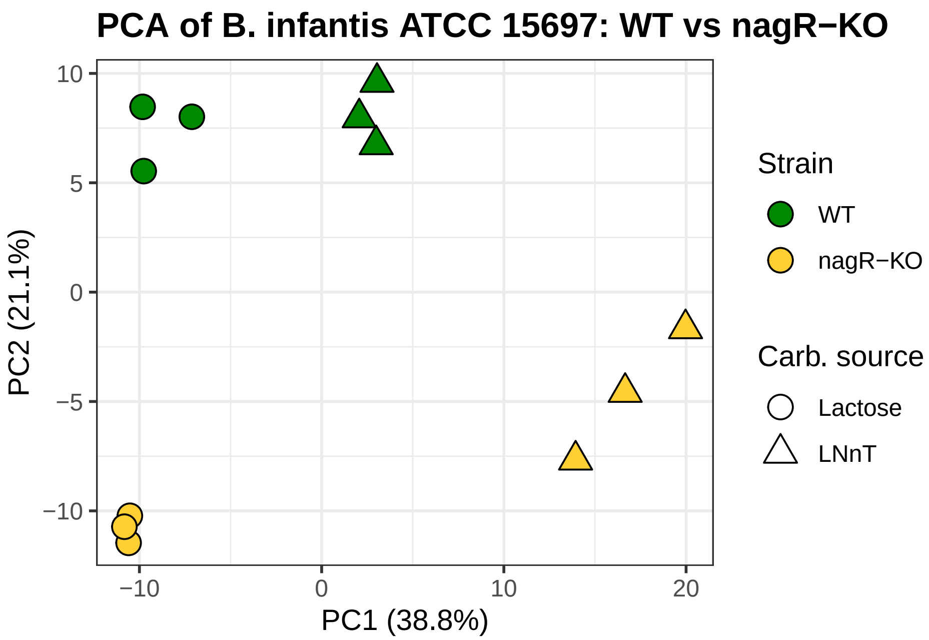

```
# save the figure
ggsave("results/figures/figure_3B.pdf", device = "pdf", width = 5, height = 5)
```

---

##### 5.5 Differentially expressed genes Table S2A-B

To identify differentially expressed genes (DEGs), precision weights were first applied to each gene based on its mean-variance relationship using VOOM. Linear modeling and bayesian stats were employed using Limma to find genes that were up- or down-regulated more than 2-fold at false-discovery rate (FDR) of 0.01.

```
# setting up model matrix without intercept
design <- model.matrix(~0 + condition)
colnames(design) <- levels(condition)
# using VOOM function from Limma package to apply precision weights to each gene
v.DEGList.filtered.norm <- voom(myDGEList.filtered.norm, design, plot = TRUE)
```

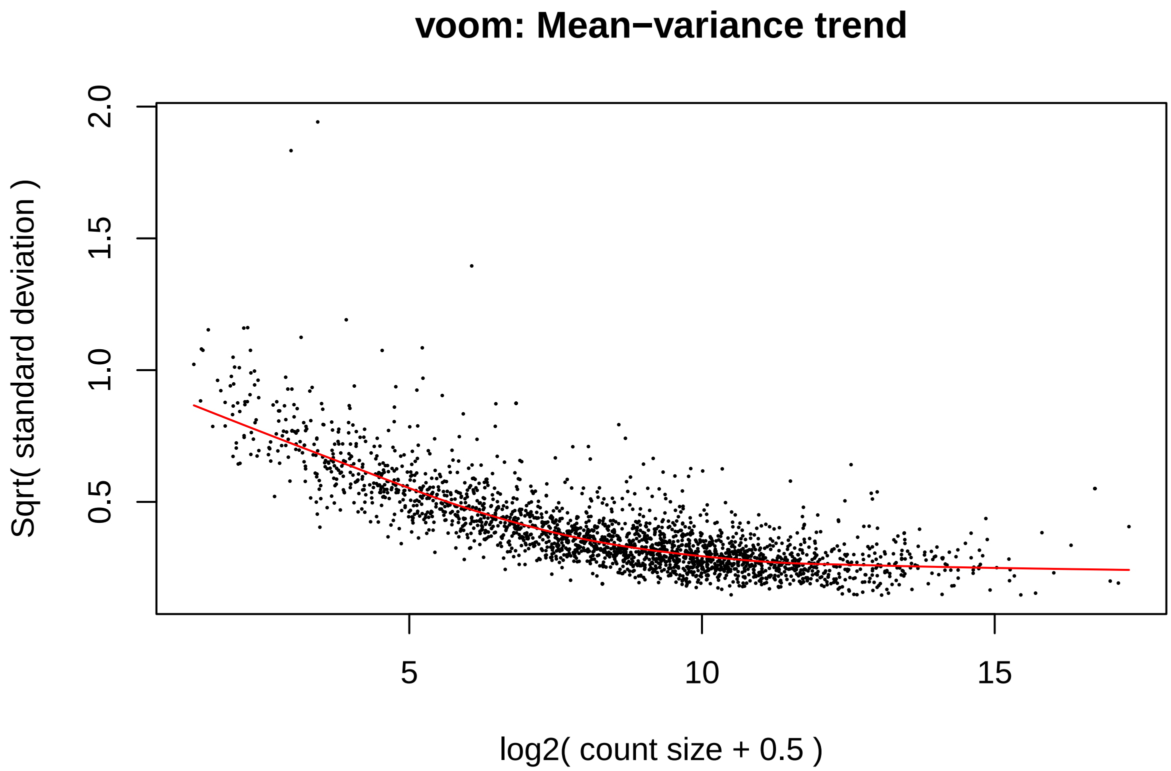

```
fit <- lmFit(v.DEGList.filtered.norm, design)
# setting up contrast matrix for pairwise comparisons of interest
contrast.matrix <- makeContrasts(Mut_WT_Lac = Mut_Lac - WT_Lac,
                                Mut_WT_LNnT = Mut_LNnT - WT_LNnT,
                                LNnT_WT = WT_LNnT - WT_Lac,
                                LNnT_Mut = Mut_LNnT - Mut_Lac,
                                levels=design)
fits <- contrasts.fit(fit, contrast.matrix)
# extracting stats
ebFit <- eBayes(fits)
```

---

DEGs were annotated based on a RAST-annotated version of the *Bifidobacterium longum* subsp. *infantis* ATCC 15697 genome, which was additionally subjected to extensive manual curation in the web-based mcSEED environment, a private clone of the publicly available SEED platform. The manual curation focused on annotating genes encoding functional roles (transporters, glycoside hydrolases, downstream catabolic enzymes, transcriptional regulators) involved in carbohydrate metabolism.

DEGs: *Bifidobacterium longum* subsp. *infantis* ATCC 15697 grown in MRS-CS-Lac: *nagR*-KO vs WT

```
# create a master annotation table
seed.ann <- read_tsv('data/rnaseq/annotation/SEED_annotations.tsv')
corr <- read_tsv('data/rnaseq/annotation/Binfantis_ATCC15697_GenBank_vs_mcSEED.txt')
final.ann <- right_join(seed.ann, corr, by = c('seed_id' = 'seed_id')) %>%
```

```
  dplyr::select(locus_tag, annotation)

# annotate DEGs
# Mut_Lac vs WT_Lac
myTopHits.Mut <- topTable(ebFit, adjust ="BH", coef=1, number=2600, sort.by="logFC")
deg_list(myTopHits.Mut, -1, 1, 0.01, "results/tables/table_S2A.txt")
```

---

DEGs: *Bifidobacterium longum* subsp. *infantis* ATCC 15697 WT: grown in MRS-CS-LNnT vs grown in MRS-CS-Lac

```
# annotate DEGs
# WT_LNnT vs WT_Lac
myTopHits.WT <- topTable(ebFit, adjust ="BH", coef=3, number=2600, sort.by="logFC")
deg_list(myTopHits.WT, -1, 1, 0.01, "results/tables/table_S2B.txt")
```

---

#### 5.6 Volcano plot: *Bifidobacterium longum* subsp. *infantis* ATCC 15697 grown in MRS-CS-Lac: *nagR*-KO vs WT Fig. 3C

Volcano plots are convenient ways to represent gene expression data because they combine magnitude of change (X-axis) with significance (Y-axis). Since the Y-axis is the inverse log10 of the adjusted Pvalue, higher points are more significant. In the case of this particular plot, there are many genes in the upper right of the plot, which represent genes that are significantly **upregulated** in the *nagR*-KO mutant grown in MRS-CS-Lac, compared to WT grown in MRS-CS-Lac.

```
# list stats for all genes in the dataset to be used for making volcano plot
myTopHits <- topTable(ebFit, adjust ="BH", coef=1, number=2600, sort.by="logFC")
myTopHits.df <- myTopHits %>%
  as_tibble(rownames = "geneID")
# select only genes with significant logFC and adj.P.Val
myTopHits.df.de <- subset(myTopHits.df, (logFC > 1 | logFC < -1) & adj.P.Val < 0.01)
# create a vector containing locus_tags of genes predicted to be in the NagR regulon
targets.nagR <- c("Blon_0879", "Blon_0881", "Blon_0882", "Blon_2171", "Blon_2172","Blon_2173",
                  "Blon_2174", "Blon_2175", "Blon_2176", "Blon_2177", "Blon_2331", "Blon_2332",
                  "Blon_2341", "Blon_2342", "Blon_2343", "Blon_2344", "Blon_2345", "Blon_2346",
                  "Blon_2347", "Blon_2348", "Blon_2349", "Blon_2350", "Blon_2351", "Blon_2352",
                  "Blon_2354")
# create a vector containing locus_tags of genes predicted to be in the CscR regulon
targets.cscR <- c("Blon_0789", "Blon_0788", "Blon_0787", "Blon_0786")
# subset volcano plot data based targets.nagR and targets.cscR
myTopHits.nagR <- subset(myTopHits.df, geneID %in% targets.nagR)
myTopHits.cscR <- subset(myTopHits.df, geneID %in% targets.cscR)
# subset data labels(NagR regulon) for volcano plot
myTopHits.df$nagR <- myTopHits.df$geneID
myTopHits.nagR_selected <- myTopHits.df$nagR %in% myTopHits.nagR$geneID
myTopHits.df$nagR[!myTopHits.nagR_selected] <- NA
# subset data labels(CscR regulon) for volcano plot
myTopHits.df$cscR <- myTopHits.df$geneID
```

```
myTopHits.cscR_selected <- myTopHits.df$cscR %in% myTopHits.cscR$geneID
myTopHits.df$cscR[!myTopHits.cscR_selected] <- NA

# create the volcano plot
ggplot(myTopHits.df) +
  aes(y=-log10(adj.P.Val), x=logFC, text = paste("Symbol:", geneID)) +
  geom_point(size=3, shape = 16, color="black", alpha=.3) +
  geom_point(mapping=NULL, myTopHits.nagR, size = 3, shape = 16, color= "sienna2",
             inherit.aes = TRUE) +
  geom_point(mapping=NULL, myTopHits.cscR , size = 3, shape = 16, color= "slateblue",
             inherit.aes = TRUE) +
  geom_text_repel(aes(label = nagR), size = 3, fontface=2, color="black", min.segment.length = 0,
                  seed = 42, box.padding = 0.5, max.overlaps = 100) +
  geom_text_repel(aes(label = cscR), size = 3, fontface=2, color="black", min.segment.length = 0,
                  seed = 42, box.padding = 0.5, max.overlaps = 100) +
  geom_hline(yintercept = -log10(0.01), linetype="longdash", colour="grey", size=0.6) +
  geom_vline(xintercept = 1, linetype="longdash", colour="#BE684D", size=0.6) +
  geom_vline(xintercept = -1, linetype="longdash", colour="#2C467A", size=0.6) +
  annotate("text", x=-6, y=-log10(0.01)+0.3,
           label=paste("Padj<0.01"), size=5, fontface="bold") +
  scale_x_continuous(limits=c(-7,5), breaks = -7:5) +
  labs(title="Volcano plot",
       subtitle = "B. infantis ATCC15697 grown in MRS-CS-Lac: nagR-KO vs WT") +
  theme(plot.title = element_text(face="bold")) +
  theme_cowplot()
```

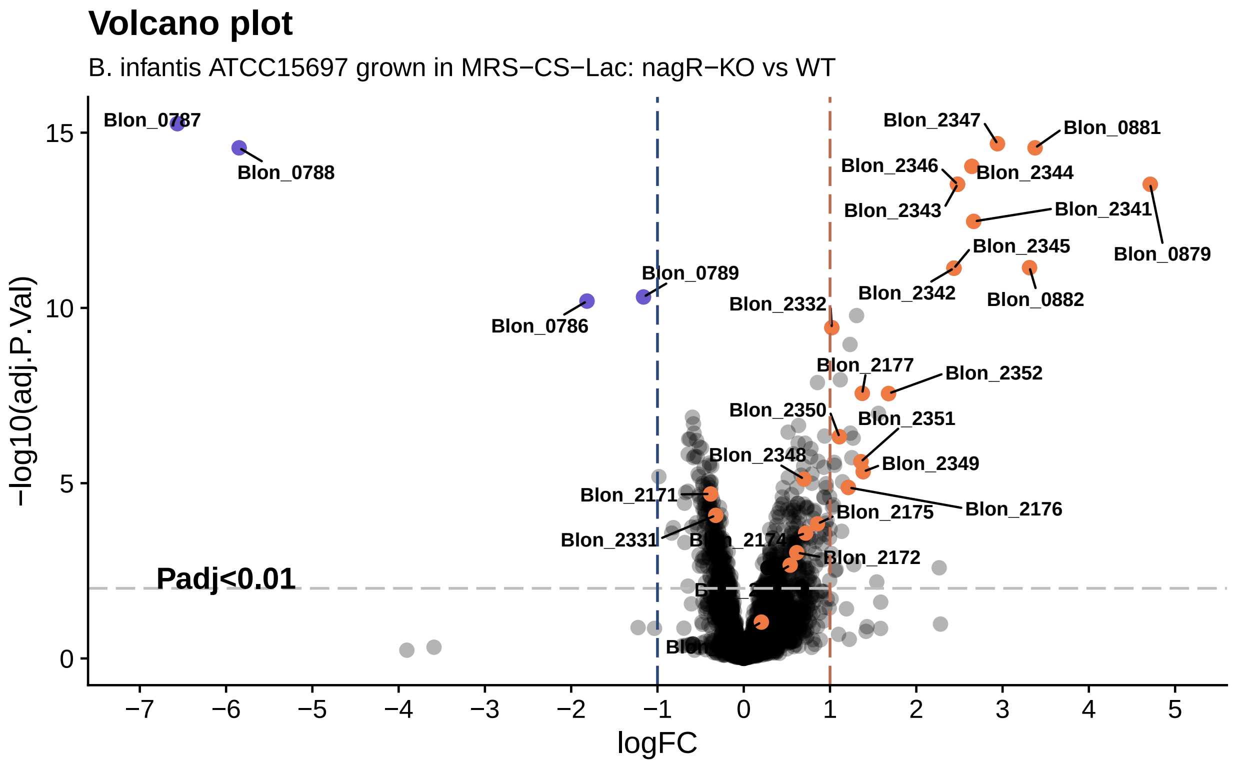

```r
# save the figure
ggsave("results/figures/figure_3C.pdf", device = "pdf", width = 8, height = 5)
```

---

#### 5.7 Volcano plot: *Bifidobacterium longum* subsp. *infantis* ATCC 15697 WT: grown in MRS-CS-LNnT vs grown in MRS-CS-Lac Fig. 3D

```r
# listing stats for all genes in the dataset to be used for making volcano plot
myTopHits3 <- topTable(ebFit, adjust ="BH", coef=3, number=2600, sort.by="logFC")
myTopHits.df3 <- myTopHits3 %>%
  as_tibble(rownames = "geneID")
# select only genes with significant logFC and adj.P.Val
myTopHits.df3.de <- subset(myTopHits.df3, (logFC > 1 | logFC < -1) & adj.P.Val < 0.01)
# subset volcano plot data based targets.nagR and targets.cscR
myTopHits.nagR3 <- subset(myTopHits.df3, geneID %in% targets.nagR)
myTopHits.cscR3 <- subset(myTopHits.df3, geneID %in% targets.cscR)
# subset volcano plot data labels (NagR-controlled genes)
myTopHits.df3$nagR <- myTopHits.df3$geneID
myTopHits.nagR_selected3 <- myTopHits.df3$nagR %in% myTopHits.nagR3$geneID
myTopHits.df3$nagR[!myTopHits.nagR_selected3] <- NA
# subset volcano plot data labels (CscR-controlled genes)
myTopHits.df3$cscR <- myTopHits.df3$geneID
myTopHits.cscR_selected3 <- myTopHits.df3$cscR %in% myTopHits.cscR3$geneID
myTopHits.df3$cscR[!myTopHits.cscR_selected3] <- NA

# create a volcano plot
ggplot(myTopHits.df3) +
  aes(y=-log10(adj.P.Val), x=logFC, text = paste("Symbol:", geneID)) +
  geom_point(size=3, shape = 16, color = "black", alpha =.3) +
  geom_point(mapping=NULL, myTopHits.nagR3, size = 3, shape = 16, color = "sienna2",
             inherit.aes = TRUE) +
  geom_point(mapping=NULL, myTopHits.cscR3 , size = 3, shape = 16, color = "slateblue",
             inherit.aes = TRUE) +
  geom_text_repel(aes(label = nagR), size = 3, fontface=2, color="black", min.segment.length = 0,
                  seed = 42, box.padding = 0.5, max.overlaps = 100) +
  geom_text_repel(aes(label = cscR), size = 3, fontface=2, color="black", mmin.segment.length = 0,
                  seed = 42, box.padding = 0.5, max.overlaps = 100) +
  geom_hline(yintercept = -log10(0.01), linetype="longdash", colour="grey", size=0.6) +
  geom_vline(xintercept = 1, linetype="longdash", colour="#BE684D", size=0.6) +
  geom_vline(xintercept = -1, linetype="longdash", colour="#2C467A", size=0.6) +
  annotate("text", x=-6, y=-log10(0.01)+0.3,
           label=paste("Padj<0.01"), size=5, fontface="bold") +
  scale_x_continuous(limits=c(-5.5,5), breaks = -6:5) +
  labs(title="Volcano plot",
       subtitle = "B. infantis ATCC15697 WT: grown in MRS-CS-LNnT vs MRS-CS-Lac") +
  theme(plot.title = element_text(face="bold")) +
  theme_cowplot()
```

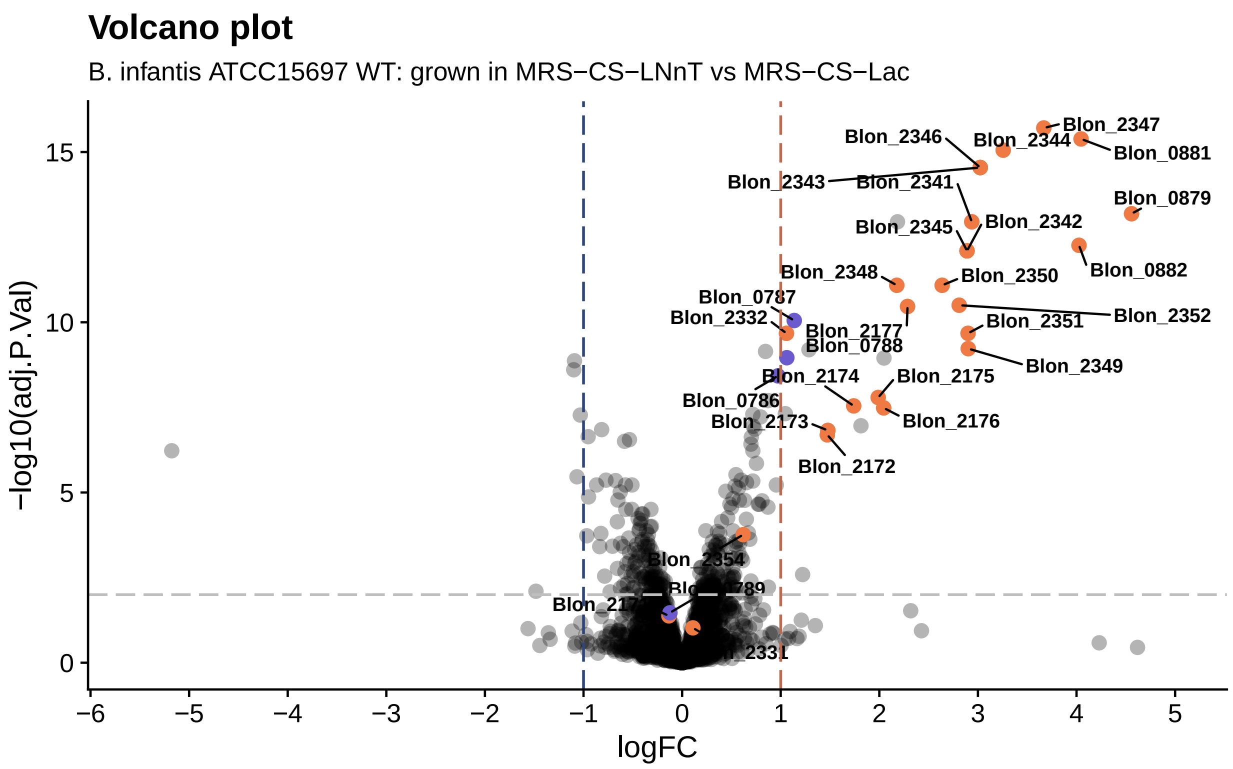

```
# save the figure
ggsave("results/figures/figure_3D.pdf", device = "pdf", width = 8, height = 5)
```

---

#### 5.8 Heatmap Fig. S3

Heatmap was plotted using the pheatmap package. Data are scaled by the row Z-score.

```
colnames(v.DEGList.filtered.norm$E) <- sampleLabels
diffGenes <- v.DEGList.filtered.norm$E
diffGenes.df <- as_tibble(diffGenes, rownames = "geneID")
diffGenes.df.two_conditions <- subset(diffGenes.df, geneID %in% myTopHits.df.de$geneID
                                      | geneID %in% myTopHits.df3.de$geneID) %>%
    relocate(geneID, WT_Lac1, WT_Lac2, WT_Lac3, Mut_Lac1, Mut_Lac2, Mut_Lac3,
             WT_LNnT1, WT_LNnT2, WT_LNnT3, Mut_LNnT1, Mut_LNnT2, Mut_LNnT3)
diffGenes.two_conditions <- as.matrix(diffGenes.df.two_conditions[,-1])
rownames(diffGenes.two_conditions) <- diffGenes.df.two_conditions$geneID
# plot the heatmap
pheatmap(diffGenes.two_conditions,
         scale = "row",
         cluster_rows = F,
         cluster_cols = F,
         angle_col = 45,
         gaps_col = c(3, 6, 9),
         cellwidth = 10,
         cellheight = 10,
```

```
       filename = "results/figures/figure_S3.pdf"
       )
```

---

### 6 Analysis of Electrophoretic Mobility Shift Assay (EMSA) data

#### 6.1 Determining $EC_{50}$ for NagR Fig. 5

$EC_{50}$ for NagR is defined as NagR concentration (nM) at which shift of 50% of a particular DNA probe (containing a predicted NagR operator) is observed in an EMSA experiment. To calculate $EC_{50}$ values, EMSA gels were visualized via Odyssey CLx. Bands were quantified in Image Studio v5.2, and shift percentages (percent of the probe shifted at a particular NagR concentration) were calculated. The resulting data (`data/emsa/NagR_data.txt`) were imported into R and approximated by a 4-parameter logistic (4PL) equation implemented in the drc package. In the 4PL model, the lower limit was fixed at 0 and the upper limit at 1.

```
# read gel quantification data
emsa_data <- read_tsv("data/emsa/NagR_data.txt")
# split data for each probe into separate tibbles
data_Blon_0879 <- filter(emsa_data, site == "Blon_0879")
data_Blon_0881_I <- filter(emsa_data, site == "Blon_0881_I")
data_Blon_2177 <- filter(emsa_data, site == "Blon_2177")
data_Blon_2344 <- filter(emsa_data, site == "Blon_2344")
data_Blon_2347 <- filter(emsa_data, site == "Blon_2347")
data_Blon_2350 <- filter(emsa_data, site == "Blon_2350")
probes_data <- lst(data_Blon_0879, data_Blon_0881_I, data_Blon_2177,
              data_Blon_2344, data_Blon_2347, data_Blon_2350)
# fit the 4-parameter logistic model
models <- lapply(probes_data, fourPL_model)
# print summary for each model
model_summary <- lapply(models, summary)
print(model_summary)
```

```
## $data_Blon_0879
##
## Model fitted: Log-logistic (ED50 as parameter) with lower limit at 0 and upper limit at 1 (2 parms)
##
## Parameter estimates:
##
##                      Estimate Std. Error t-value   p-value
## Slope:(Intercept) -3.41086    0.38351 -8.8939 2.022e-05 ***
## EC50:(Intercept)  13.48218    0.48959 27.5378 3.261e-09 ***
## ---
## Signif. codes:  0 '***' 0.001 '**' 0.01 '*' 0.05 '.' 0.1 ' ' 1
##
## Residual standard error:
##
##  0.04324023 (8 degrees of freedom)
##
## $data_Blon_0881_I
```

```
##
## Model fitted: Log-logistic (ED50 as parameter) with lower limit at 0 and upper limit at 1 (2 parms)
##
## Parameter estimates:
##
##                    Estimate Std. Error t-value   p-value
## Slope:(Intercept) -2.20635    0.44946 -4.9088  0.001181 **
## EC50:(Intercept)  22.65238    1.81796 12.4603 1.608e-06 ***
## ---
## Signif. codes:  0 '***' 0.001 '**' 0.01 '*' 0.05 '.' 0.1 ' ' 1
##
## Residual standard error:
##
##  0.06889999 (8 degrees of freedom)
##
## $data_Blon_2177
##
## Model fitted: Log-logistic (ED50 as parameter) with lower limit at 0 and upper limit at 1 (2 parms)
##
## Parameter estimates:
##
##                     Estimate Std. Error t-value   p-value
## Slope:(Intercept)  -0.92028    0.11233 -8.1928 3.677e-05 ***
## EC50:(Intercept)  140.39781   23.07344  6.0848 0.0002944 ***
## ---
## Signif. codes:  0 '***' 0.001 '**' 0.01 '*' 0.05 '.' 0.1 ' ' 1
##
## Residual standard error:
##
##  0.07237161 (8 degrees of freedom)
##
## $data_Blon_2344
##
## Model fitted: Log-logistic (ED50 as parameter) with lower limit at 0 and upper limit at 1 (2 parms)
##
## Parameter estimates:
##
##                    Estimate Std. Error t-value   p-value
## Slope:(Intercept) -7.07459    0.79865 -8.8582 2.082e-05 ***
## EC50:(Intercept)  14.03981    0.23161 60.6174 6.096e-12 ***
## ---
## Signif. codes:  0 '***' 0.001 '**' 0.01 '*' 0.05 '.' 0.1 ' ' 1
##
## Residual standard error:
##
##  0.03009977 (8 degrees of freedom)
##
## $data_Blon_2347
##
## Model fitted: Log-logistic (ED50 as parameter) with lower limit at 0 and upper limit at 1 (2 parms)
##
## Parameter estimates:
##
##                    Estimate Std. Error t-value   p-value
```

```
## Slope:(Intercept) -3.07428    0.34441 -8.9261  0.000294 ***
## EC50:(Intercept)  13.42257    0.58037 23.1278 2.812e-06 ***
## ---
## Signif. codes:  0 '***' 0.001 '**' 0.01 '*' 0.05 '.' 0.1 ' ' 1
##
## Residual standard error:
##
##  0.03630326 (5 degrees of freedom)
##
## $data_Blon_2350
##
## Model fitted: Log-logistic (ED50 as parameter) with lower limit at 0 and upper limit at 1 (2 parms)
##
## Parameter estimates:
##
##                     Estimate Std. Error t-value   p-value
## Slope:(Intercept)  -0.84253    0.13914 -6.0553 0.0003041 ***
## EC50:(Intercept)  178.26884   41.23875  4.3228 0.0025363 **
## ---
## Signif. codes:  0 '***' 0.001 '**' 0.01 '*' 0.05 '.' 0.1 ' ' 1
##
## Residual standard error:
##
##  0.09711114 (8 degrees of freedom)
```

Plot quantification data and the resulting 4PL fit with 95% confidence intervals.

```
# plot data
plot_Blon_0879 <- plot_fit(models$data_Blon_0879, data_Blon_0879, 1000)
plot_Blon_0881_I <- plot_fit(models$data_Blon_0881_I, data_Blon_0881_I, 1000)
plot_Blon_2177 <- plot_fit(models$data_Blon_2177, data_Blon_2177, 2000)
plot_Blon_2344 <- plot_fit(models$data_Blon_2344, data_Blon_2344, 1000)
plot_Blon_2347 <- plot_fit(models$data_Blon_2347, data_Blon_2347, 1000)
plot_Blon_2350 <- plot_fit(models$data_Blon_2350, data_Blon_2350, 2000)
plot_grid(plot_Blon_0879, plot_Blon_0881_I, plot_Blon_2177,
               plot_Blon_2344, plot_Blon_2347, plot_Blon_2350,
          labels = c('Blon_0879', 'Blon_0881_I', 'Blon_2177',
                     'Blon_2344', 'Blon_2347', 'Blon_2350'),
          ncol = 1,
          label_size = 12,
          label_x = 0, label_y = 0,
          hjust = -0.5, vjust = -0.5)
```

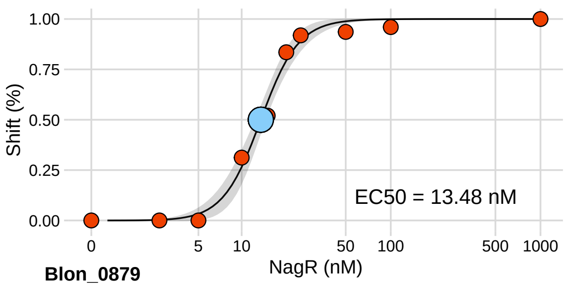

**Blon_0879**

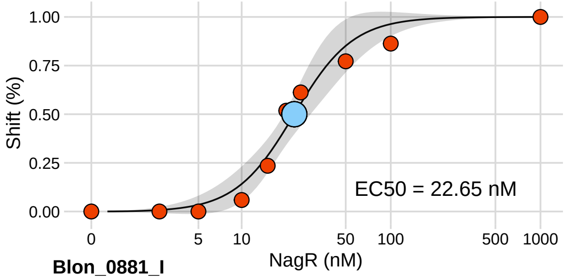

**Blon_0881_I**

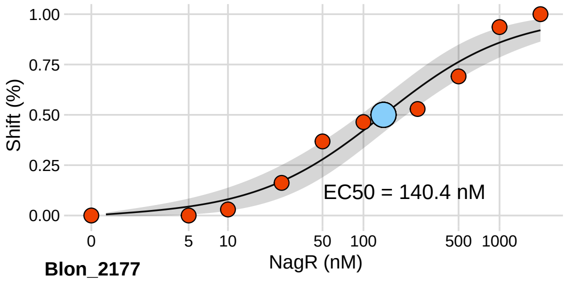

**Blon_2177**

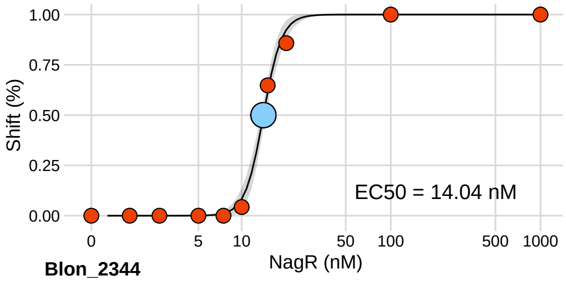

**Blon_2344**

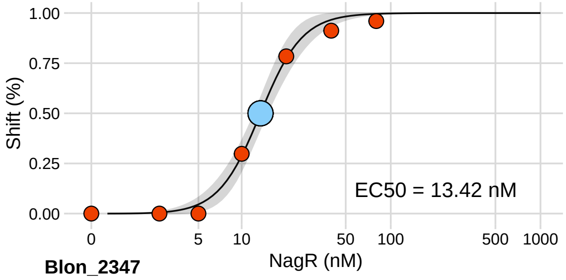

**Blon_2347**

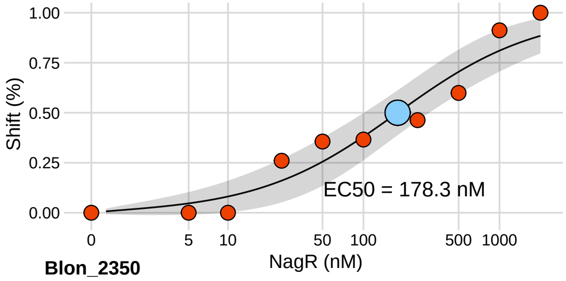

**Blon_2350**

```
# save the figure
ggsave("results/figures/figure_5.pdf", device = cairo_pdf, width = 5, height = 15)
```

---

#### 6.2 Correlation between RNA-seq and EMSA-data Fig. 4C

Pearson correlation coefficient was calculated between $\log_2 EC_{50}$ values determined by EMSA and $\log_2$(FoldChange) of genes in the RNA-seq experiment (*nagR*-KO grown in MRS-CS-Lac vs WT grown in MRS-CS-Lac).

```
# read data
emsa_data_corr <- read_tsv("data/emsa/NagR_EMSA_vs_RNA_seq.txt")
# plot data
ggplot(emsa_data_corr, aes(x = logFC_mut_Lac, y = logEC50)) +
  geom_point(size = 3) +
  geom_smooth(method = lm) +
  scale_x_continuous(name ="log2FC") +
  scale_y_continuous(name ="log2EC50") +
  # print Pearson R^2
  stat_cor(method = "pearson",
           label.y = 1.0,
           aes(label = paste(..r.label.., ..p.label.., sep = "~`,`~"))) +
  theme_cowplot()
```

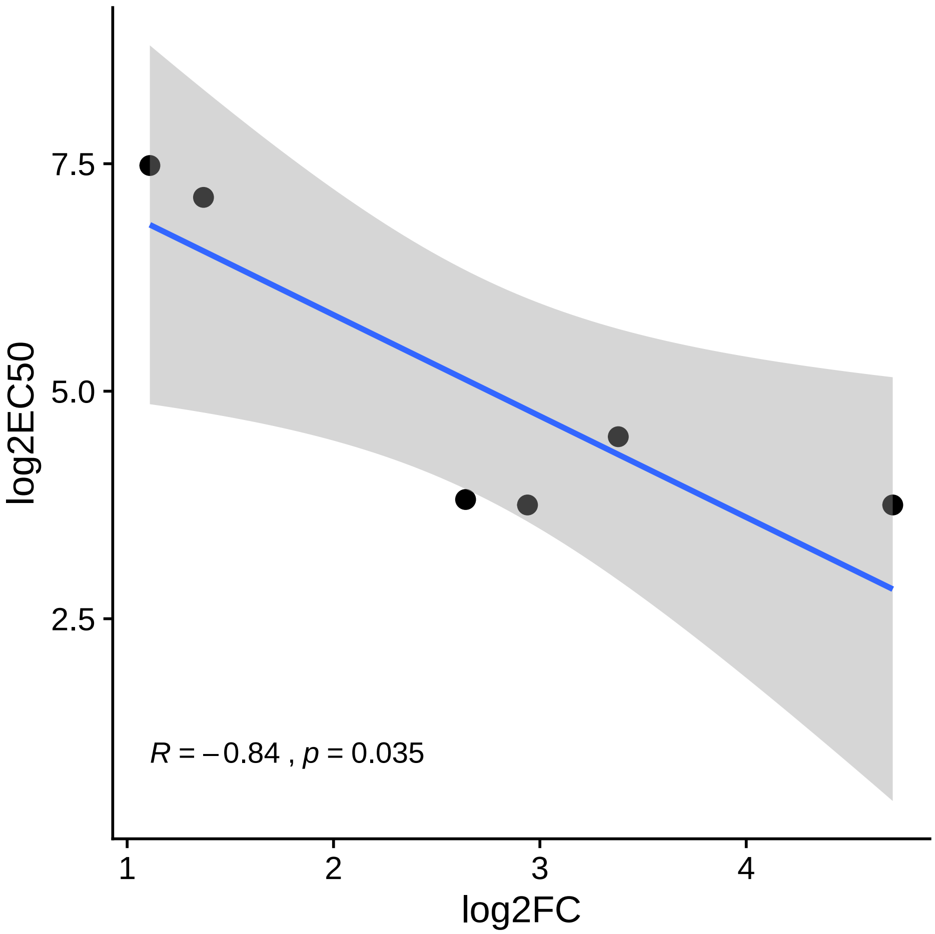

```
# save the figure
ggsave("results/figures/figure_4C.pdf", device = cairo_pdf, width = 5, height = 5)
```

---

##### 6.3 Determining $EC_{50}$ values for NagR effectors Fig. 4D

$EC_{50}$ value for an effector is defined as effector concentration which inhibits 50% of the "control" shift (shift without the effector). To calculate $EC_{50}$ values, EMSA gels were visualized via Odyssey CLx. Bands were quantified in Image Studio v5.2, and shift percentages (how much of the probe is shifted at a particular effector concentration) were calculated. The shift percentage w/ effector was divided by shift percentage w/o effector ("control" shift). The resulting data (`data/emsa/NagR_effector_data.txt`) were imported into R and approximated by a 4PL equation implemented in the drc package. In the 4PL model, the upper limit was fixed at 1.

```
# 4-parameter logistic model; upper limit is fixed to 1
# input: tibble with data
fourPL_model_eff <- function(site_data){
  drm(percent_max_shift~conc_mM,
      data = site_data,
      fct = LL.4(fixed=c(NA,NA,1,NA), names=c("Slope", "Min", "Max", "EC50")))
```

```r
}
# read gel quantification data
emsa_data_eff <- read_tsv("data/emsa/NagR_effector_data.txt")
# split data for each effector into separate tibbles
data_GlcNAc <- filter(emsa_data_eff, effector == "GlcNAc")
data_GlcNAc1P <- filter(emsa_data_eff, effector == "GlcNAc1P")
data_GlcNAc6P <- filter(emsa_data_eff, effector == "GlcNAc6P")
probes_data_eff <- lst(data_GlcNAc, data_GlcNAc1P, data_GlcNAc6P)
# fit the 4-parameter logistic model
models_eff <- lapply(probes_data_eff, fourPL_model_eff)
# print summary for each model
model_summary_eff <- lapply(models_eff, summary)
print(model_summary_eff)
```

```
## $data_GlcNAc
##
## Model fitted: Log-logistic (ED50 as parameter) (3 parms)
##
## Parameter estimates:
##
##                    Estimate Std. Error t-value   p-value
## Slope:(Intercept) 2.100877   0.494005   4.2527 0.0027887 **
## Min:(Intercept)   0.527303   0.023641 22.3046 1.726e-08 ***
## EC50:(Intercept)  0.327653   0.047489   6.8996 0.0001246 ***
## ---
## Signif. codes:  0 '***' 0.001 '**' 0.01 '*' 0.05 '.' 0.1 ' ' 1
##
## Residual standard error:
##
##  0.0409181 (8 degrees of freedom)
##
## $data_GlcNAc1P
##
## Model fitted: Log-logistic (ED50 as parameter) (3 parms)
##
## Parameter estimates:
##
##                    Estimate Std. Error t-value   p-value
## Slope:(Intercept) 1.677098   0.315507   5.3156 0.0007148 ***
## Min:(Intercept)   0.132259   0.038519   3.4336 0.0089070 **
## EC50:(Intercept)  0.487074   0.057809   8.4255 3.001e-05 ***
## ---
## Signif. codes:  0 '***' 0.001 '**' 0.01 '*' 0.05 '.' 0.1 ' ' 1
##
## Residual standard error:
##
##  0.05658287 (8 degrees of freedom)
##
## $data_GlcNAc6P
##
## Model fitted: Log-logistic (ED50 as parameter) (3 parms)
##
## Parameter estimates:
```

```
##
##                     Estimate Std. Error t-value p-value
## Slope:(Intercept)  2.79546    8.26298  0.3383  0.7521
## Min:(Intercept)    0.50531    0.54244  0.9315  0.4043
## EC50:(Intercept)   4.46491    7.74990  0.5761  0.5954
##
## Residual standard error:
##
##  0.06341104 (4 degrees of freedom)
```

Plot quantification data and the resulting 4PL fit with 95% confidence intervals.

```
# GlcNAc fit for ggplot
conf_site_GlcNAc <- expand.grid(conc_mM=exp(seq(log(0.01), log(10), length=100)))
# calculate predicted values and respective 95% confidence intervals
# from the model using the "predict" function
pm_site_GlcNAc <- predict(models_eff$data_GlcNAc,
                          newdata=conf_site_GlcNAc,
                          interval="confidence")
# new data with predictions/confidence intervals
conf_site_GlcNAc$p <- pm_site_GlcNAc[,1]
conf_site_GlcNAc$pmin <- pm_site_GlcNAc[,2]
conf_site_GlcNAc$pmax <- pm_site_GlcNAc[,3]
EC50_GlcNAc <- models_eff$data_GlcNAc$coefficients[3]
new_GlcNAc <- data.frame(x=c(EC50_GlcNAc))
yvalue_GlcNAc <- predict(models_eff$data_GlcNAc, newdata = new_GlcNAc)
# GlcNAc1P fit for ggplot
conf_site_GlcNAc1P <- expand.grid(conc_mM=exp(seq(log(0.01), log(10), length=100)))
# calculate predicted values and respective 95% confidence intervals
# from the model using the "predict" function
pm_site_GlcNAc1P <- predict(models_eff$data_GlcNAc1P,
                            conf_site_GlcNAc1P,
                            interval="confidence")
# new data with predictions/confidence intervals
conf_site_GlcNAc1P$p <- pm_site_GlcNAc1P[,1]
conf_site_GlcNAc1P$pmin <- pm_site_GlcNAc1P[,2]
conf_site_GlcNAc1P$pmax <- pm_site_GlcNAc1P[,3]
EC50_GlcNAc1P <- models_eff$data_GlcNAc1P$coefficients[3]
new_GlcNAc1P <- data.frame(x=c(EC50_GlcNAc1P))
yvalue_GlcNAc1P <-  predict(models_eff$data_GlcNAc1P, newdata = new_GlcNAc1P)
# plot data
ggplot() +
  geom_line(data=conf_site_GlcNAc, aes(x=conc_mM, y=p), color = "#ffcf34") +
  geom_ribbon(data=conf_site_GlcNAc, aes(x=conc_mM, y=p, ymin=pmin, ymax=pmax), alpha=0.2) +
  geom_line(data=conf_site_GlcNAc1P, aes(x=conc_mM, y=p), color = "#008800") +
  geom_ribbon(data=conf_site_GlcNAc1P, aes(x=conc_mM, y=p, ymin=pmin, ymax=pmax), alpha=0.2) +
  geom_point(data = data_GlcNAc, aes(x = conc_mM, y = percent_max_shift, fill="GlcNAc"),
             size=4, pch=21) +
  geom_point(data = data_GlcNAc1P, aes(x = conc_mM, y = percent_max_shift, fill="GlcNAc1P"),
             size=4, pch=21) +
  geom_point(aes(x = EC50_GlcNAc, y = yvalue_GlcNAc, fill="GlcNAc"), size = 7, pch=21) +
  geom_point(aes(x = EC50_GlcNAc1P, y = yvalue_GlcNAc1P, fill="GlcNAc1P"), size = 7, pch=21) +
  geom_point(aes(x = 0, y = 1), size = 4, pch=21, fill = "black") +
  scale_fill_manual(name="Effector",
```

```
    values=c(GlcNAc="#ffcf34", GlcNAc1P="#008800")) +
  scale_x_continuous(trans=scales::pseudo_log_trans(base=10),
                     limits=c(0, 10),
                     breaks=c(0, 0.5, 1, 3, 5, 10)) +
  scale_y_continuous(limits=c(0, 1),
                     breaks=c(0, 0.25, 0.5, 0.75, 1)) +
  annotate("text", x=1.7, y=0.8,
           label= paste("EC50 =",signif(EC50_GlcNAc, 4), "mM"),
           size = 4.5) +
  annotate("text", x=1, y=0.4,
           label= paste("EC50 =",signif(EC50_GlcNAc1P, 4), "mM"),
           size = 4.5) +
  xlab("Effector (mM)") +
  ylab("Shift w/ effector / Shift w/o effector (%)") +
  theme_cowplot(12)
```

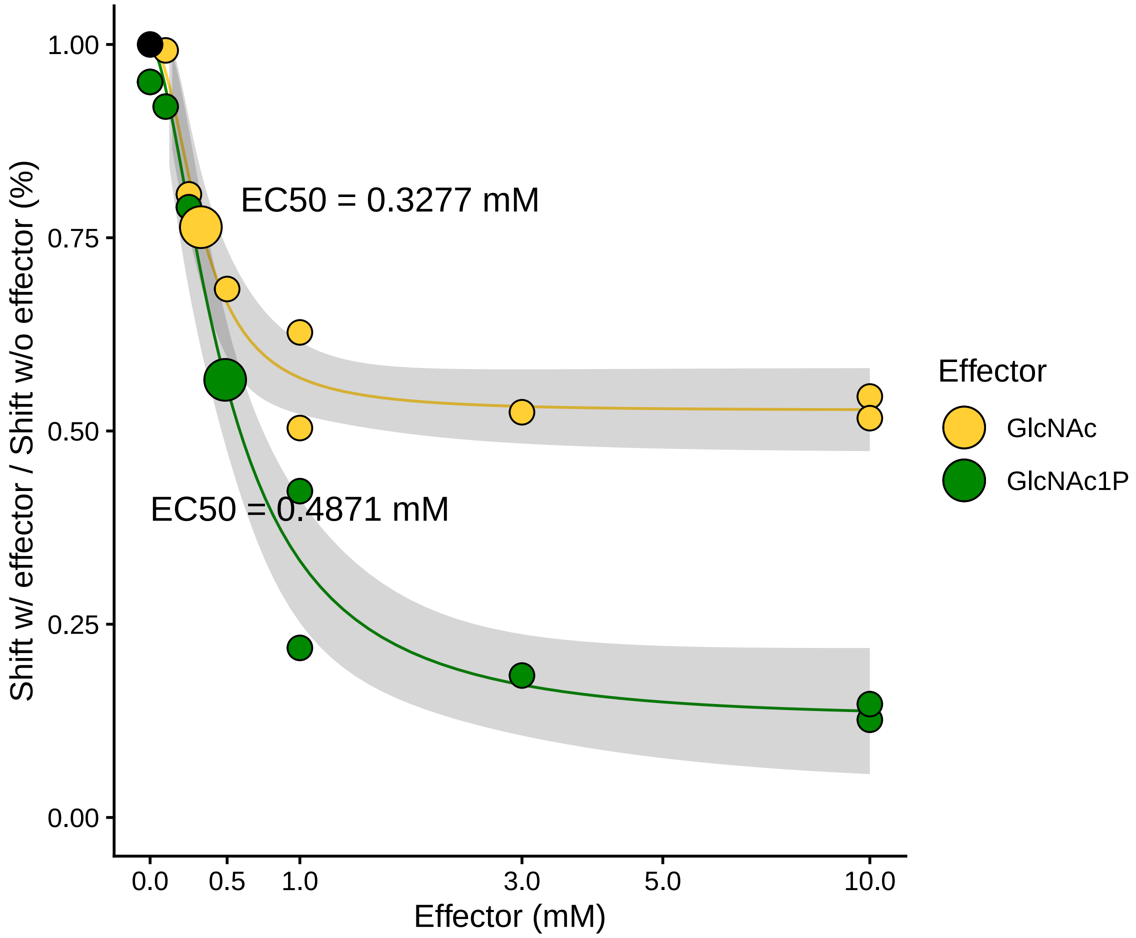

```
# save the figure
ggsave("results/figures/figure_4E.pdf", device = cairo_pdf, width = 6, height = 5)
```

### 7 Phylogenetic analysis

#### 7.1 Processing of multiple sequence alignments (msa)

The code chuck below describes:

1. Trimming of the alignment of NagR proteins from 25 *Bifidbacteriaceae* genomes. The alignment was generated via MUSCLE (v3.8.31)
2. Trimming of the alignment of 247 core genes from 25 *Bifidbacteriaceae* genomes. Core genes were identified via Roary (v3.12.0) and the following command: `roary -p 16 -e -n -i 70 -f pangenome_70_percent *.gff`. The alignment was generated via MAFFT (v7.313)

```
# NagR proteins
# read fasta file with msa
msa.nagR <- readFasta("results/evolution/NagR_aligned.fa")
#str_length(msa.nagR$Sequence[1])
# trim msa
msa.nagR.trimmed <- msaTrim(msa.nagR, gap.end = 0.9, gap.mid = 0.9)
#str_length(msa.nagR.trimmed$Sequence[1])

# write fasta file with trimmed msa
writeFasta(msa.nagR.trimmed, "results/evolution/NagR_aligned_trimmed.fa", width = 0)

# species core genes
# read fasta file with msa
msa.species <- readFasta("results/evolution/species_core_genes.aln")
#str_length(msa.species$Sequence[1])
# trim msa
msa.species.trimmed <- msaTrim(msa.species, gap.end = 0.9, gap.mid = 0.9)
#str_length(msa.species.trimmed$Sequence[1])

# write fasta file with trimmed msa
writeFasta(msa.species.trimmed, "results/evolution/species_core_genes_trimmed.aln", width = 0)
```

The resulting trimmed alignments were used to build phylogenetic trees in IQ-TREE (v1.6.12). The following commands were used:

1. `iqtree -s NagR_aligned_trimmed.fa -m LG+F+G4 -bb 1000 -nt AUTO -ntmax 16` for NagR tree
2. `iqtree -s core_gene_alignment_trimmed.aln -m GTR+F+R10 -bb 1000 -nt AUTO -ntmax 16` for species tree

Constructed phylogenetic trees, as well as files for visualization in iTOL, can be found in `results/evolution/`.

---

### 8 Session info

The output from running 'sessionInfo' is shown below and details all packages necessary to reproduce the results in this report.

```
sessionInfo()
```

```
## R version 4.1.2 (2021-11-01)
## Platform: x86_64-apple-darwin17.0 (64-bit)
## Running under: macOS Catalina 10.15.7
##
## Matrix products: default
## BLAS:   /Library/Frameworks/R.framework/Versions/4.1/Resources/lib/libRblas.0.dylib
## LAPACK: /Library/Frameworks/R.framework/Versions/4.1/Resources/lib/libRlapack.dylib
##
## locale:
## [1] en_US.UTF-8/en_US.UTF-8/en_US.UTF-8/C/en_US.UTF-8/en_US.UTF-8
##
## attached base packages:
## [1] stats     graphics  grDevices datasets  utils     methods   base
##
## other attached packages:
##  [1] microseq_2.1.5     rlang_0.4.12       data.table_1.14.2  ggpubr_0.4.0
##  [5] Cairo_1.5-14       drc_3.0-1          MASS_7.3-54        pheatmap_1.0.12
##  [9] ggrepel_0.9.1      cowplot_1.1.1      matrixStats_0.61.0 edgeR_3.36.0
## [13] limma_3.50.0       gt_0.3.1           tximport_1.22.0    emmeans_1.7.2
## [17] forcats_0.5.1      stringr_1.4.0      dplyr_1.0.7        purrr_0.3.4
## [21] readr_2.1.1        tidyr_1.1.4        tibble_3.1.6       ggplot2_3.3.5
## [25] tidyverse_1.3.1    pacman_0.5.1       knitr_1.37         tinytex_0.36
## [29] rmarkdown_2.11
##
## loaded via a namespace (and not attached):
##  [1] TH.data_1.1-0       colorspace_2.0-2    ggsignif_0.6.3
##  [4] ellipsis_0.3.2      htmlTable_2.4.0     estimability_1.3
##  [7] base64enc_0.1-3     fs_1.5.2            rstudioapi_0.13
## [10] farver_2.1.0        bit64_4.0.5         fansi_0.5.0
## [13] mvtnorm_1.1-3       lubridate_1.8.0     xml2_1.3.3
## [16] codetools_0.2-18    splines_4.1.2       Formula_1.2-4
## [19] jsonlite_1.7.2      broom_0.7.11        cluster_2.1.3
## [22] dbplyr_2.1.1        png_0.1-7           BiocManager_1.30.16
## [25] compiler_4.1.2      httr_1.4.2          backports_1.4.1
## [28] assertthat_0.2.1    Matrix_1.4-0        fastmap_1.1.0
## [31] cli_3.1.0           htmltools_0.5.2     tools_4.1.2
## [34] gtable_0.3.0        glue_1.6.0          Rcpp_1.0.7
## [37] carData_3.0-5       cellranger_1.1.0    rhdf5filters_1.6.0
## [40] vctrs_0.3.8         nlme_3.1-153        xfun_0.29
## [43] rvest_1.0.2         lifecycle_1.0.1     renv_0.15.0
## [46] gtools_3.9.2        rstatix_0.7.0       zoo_1.8-9
## [49] scales_1.1.1        vroom_1.5.7         hms_1.1.1
## [52] parallel_4.1.2      sandwich_3.0-1      rhdf5_2.38.1
## [55] RColorBrewer_1.1-2  yaml_2.2.1          gridExtra_2.3
## [58] rpart_4.1-15        latticeExtra_0.6-29 stringi_1.7.6
## [61] highr_0.9           plotrix_3.8-2       checkmate_2.0.0
## [64] pkgconfig_2.0.3     evaluate_0.14       lattice_0.20-45
## [67] Rhdf5lib_1.16.0     labeling_0.4.2      htmlwidgets_1.5.4
## [70] bit_4.0.4           tidyselect_1.1.1    magrittr_2.0.1
## [73] R6_2.5.1            generics_0.1.1      Hmisc_4.6-0
## [76] multcomp_1.4-18     DBI_1.1.2           mgcv_1.8-38
```

```
## [79] pillar_1.6.4        haven_2.4.3         foreign_0.8-81
## [82] withr_2.4.3         nnet_7.3-16         survival_3.2-13
## [85] abind_1.4-5         modelr_0.1.8        crayon_1.4.2
## [88] car_3.0-12          utf8_1.2.2          tzdb_0.2.0
## [91] jpeg_0.1-9          locfit_1.5-9.4      grid_4.1.2
## [94] readxl_1.3.1        reprex_2.0.1        digest_0.6.29
## [97] xtable_1.8-4        munsell_0.5.0
```
