## Supplementary material for "Human Milk Oligosaccharide Utilization in Intestinal Bifidobacteria is Governed by a Global Transcriptional Regulator NagR": Figure S1

**FIG S1.** Multiple sequence alignments of upstream regions of NagR-regulated genes in *Bifidobacterium* spp. Predicted NagR operators are highlighted in yellow. Predicted promoter elements (-35 and -10 sequences) and in bold black. Experimentally described transcription start sites (TSS) in *B. breve* UCC2003 (65) are in bold blue. Predicted Shine-Dalgarno sequences are in bold red. Coding sequences are highlighted in grey; initiation codons are underlined. Conserved nucleotides are marked with asterisks.

***nagK* (Blon_0879)**

```
B. infantis ATCC15697   AGTGTCGCCTTGGGTGAATTTGTTAAGATAGTTGTCAATAGTCGGATGCACACTTATACCAAGCGAAACGTTCGATGAAGAACAGGGGAGGGCTGATGTCCGAAA
B. longum NCC2705       AGTGTCGTCTTGGGGGAATTTGTTAAGATGGTTGTCAATAGTCGGATGCACACTGATATCAAGCGAAACGTTCGATGAAGAACAGGGGAGGACTGATGTCTGAAA
B. suis JDM301          AGTGTCGTCTTGGGGGAATTTGTTAAGATGGTTGTCAATAGTCGGATGCACACTGATATCAAGCGAAACGTTCGATGAAGAACAGGGGAGGACTGATGTCTGAAA
B. breve UCC2003        GCGCATGGTTTTCTGTAATTTGTTAAGATAGTTGTCAATACATAAATGCATACTGATTCCAAGCATATTGTTCGACGAAGAACGGGGGATAGGCAATGACTGATA
                              *  **     ************* **********     ***** *** **  *****  *  ****** ******* *****      *** * ** *
```

***nagB* (Blon_0881)**

```
B. infantis ATCC15697   CAGTTTAGACGGTGAAGCACAATTTCGTCGTTGAAATTGTTATGAAACTTCACCAATAAATTCCAGTGTACCTGTGCGAAGCGTCAGGAAACACCGTGAGTGTTC
B. longum NCC2705       CAGCTTAGACGGTGAAGCGCAATTTCGTCGTTGAAACTGTTATGAAACTTCACTAATAAATTCCAGTGTACCTGTGTGAAGCGTCAGGAAACACCGTGAGTGTTC
B. suis JDM301          CAGCTTAGACGGTGAAGCGCAATTTCGTCGTTGAAACTGTTATGAAACTTCACTAATAAATTCCAGTGTACCTGTGTGAAGCGTCAGGAAACACCGTGAGTGTTC
B. breve UCC2003        ACCACACGGTATGTGAGATTCCGCTCGGCGTTGAATCTGTTAGGGAACTTCACTAATACATTCCAGTGTACTCGATTGTTCTGGTTTGGGAA--CGCCGAGTGTC
                               *       **       *** *******  ***** * ******** **** ************  *   *    *    *  *   **       **

B. infantis ATCC15697   ACAAGGTCCAAAATTGTTAGTATCATTAACAAATAGCGAATCG----------CCAATCGGCACTATGCCGAATGGTGGTTGGAACGCCGCACATCACCCGGACG
B. longum NCC2705       ATAAGGTCCAAAATTGTTAGTATCATTAACAAATAGCGAATCA----------CCAATCGGCGCTACGCCGAACGGTGGTTGGAACGCCGCGCGGCACCCGGACG
B. suis JDM301          ATAAGGTCCAAAATTGTTAGTATCATTAACAAATAGCGAATCA----------CCAATCGGCGCTACGCCGAACGGTGGTTGGAACGCCGCGCGGCACCCGGACG
B. breve UCC2003        CGTACGTCCATAATTGTAAGTAGTATTAACAAATAACTGGTTAGGTTGATGCGAACATCGACATAATCGGATTGCACATTG-----------------TGGAAAG
                           * ***** ****** ****  *********** *   *               **** *   *             *                    * * *

B. infantis ATCC15697   TGACAAGGATGTTGCCATTATCCGCAAGACGGGCGGCAGTACAGCGACTCGCATCGGGCGGGCCGCACGGAA-CTGGAGGAAACCGATGCCGGAAATCAT
B. longum NCC2705       TGACAAGGATGTTGCCATTATCCGCAAGACGGGCGGCGGTACAGCGACTCGCATCGGGCGGGCCGCACGGAA-CTGGAGGAAACCGATGCCGGAAATCAT
B. suis JDM301          TGACAAGGATGTTGCCATTATCCGCAAGACGGGCGGCGGTACAGCGACTCGCATCGGGCGGGCCGCACGGAA-CTGGAGGAAACCGATGCCGGAAATCAT
B. breve UCC2003        TGACAAAGAAGTTGCCATGAACCATGATGGGCGGCTTTGGGCAGAAGCCCGTGTCTAGCGGGCCTCATGAACGGAGGAAGATACCTATGCCGGAAATCAT
                        ****** ** ******** * **   *   * *     *  ***   * **  **  ******* ** * *    *** ** *** **************
```

***gltA* (Blon_2177)**

```
B. infantis ATCC15697   TAATTGTTAGTTGGGTTGACAATAAAAGAGCAAAGGCCTAAATTAGCA-ATCAGTCATCACACTGAGGTGATG-----ATTCCATTA--CGAAGGAGT-AGAGATATG
B. longum NCC2705       TAATTGTTAGCTAAGTTGACAATAAAAGAGCAAAGGCCTAAATTAGCA-ATCAGTCATCACGCTGAGGTGATG-----ATTTCCCTA--GGAAGGAGT-AGAGATATG
B. suis JDM301          TAATTGTTAGCTAAGTTGACAATAAAAGAGCAAAGGCCTAAATTAGCA-ATCAGTCATCACGCTGAGGTGATG-----ATTTCACTA--GGAAGGAGT-AGAGATATG
B. breve UCC2003        AAATTGTTAGTTAGGTTGACAATAAAGATTTGAGCGCCTAAACTGATT-AACAGTCGTCACAGAGGAGTGATG-----ACGCCATTAT-CGAAGGAGT-AGAGATATG
B. bifidum PRL2010      CAAATGTTAATGTAGTTGACATTTCCTTGGCATGCCTTTACACTAACTAATTAGTTGCCACGACGAGGAGGCAACATCCATTTCAAT---GGAGGAGTTAGGAATATG
B. pseudocat. JCM1200   TAAATGTAAGGAAACTTGACAAAAAAGGATTACGTAGGTATGCTGTGA-CTATCCGGTGCAAGGAGGTGCCGG-----AATAATCTC-ACGAAGGAG--AGAGATATG
B. scardovii JCM12489   TTATTGTTAAGGTAGTTGACATTTATTGTTTCCTGTTCTAAAGTAAAT-ACCGGTTGCCGCACCGCCGTGGCAGC-GTTTCATCTCG--TGGAGGAGT-GGAAACATG
                          * *** *      ******                 **   *                                              * *****   *  * ***
```

***lacS* (Blon_2332)**

```
B. infantis ATCC15697   -CCGTCCAAAAAAACATCCGCACCCCCGTGATAAAACTTGTATGCTTACTTTACAAAATCAATGAGTGATTCCATGCTCTGGGTTCTTAAGGAGAAGATTAATGA
B. longum NCC2705       -ACGGTGATAAAACTTGTATGCTTACGGTGATAAAACTTGTATGCTTACTTTACAAAATCAATGAGTGATTCCATGCTCTGGGTTCTTAAGGAGAAGATTAATGA
B. suis JDM301          -CCGTTTACTAATGTTTTTTACTTTCGGTGATAAAACTTGTATGCTTACTTTACAAAATCAATGAGTGATTCCATGCTCTGGGTTCTTAAGGAGAAGATTAATGA
B. longum 1897B         CGCTGTTTATCTTGCTCTTTACTTTCGGTGATAAAACTTGTATGCTTACTTTACAAAGTCAATGGATGATTCCA-GCTCTGGGATCTTAAGGAGAAGACTAATGA
                          *                      * ****************************** ******  ******** ******** ********** *** ******
```

***hmoA2* (Blon_2344)**

```
B. infantis ATCC15697    CCCCTTTGTTCATTCGGCCGTCGCGCGCTTCCCCGGTCGCCCGTCGTGGTGCCACATATTGTTAGGCATGTTGACAAAATGCTGCGAAGAGGCATATATTACTGT
B. infantis IN-F29       CCCCTTTGTTCATTCGGCCGTCGCGCGTTTCCCCGGTCGCCCGTCGTGGTGCCGCATATTGTTAGGCATGTTGACAAAATGCTGCGAAGAGGCATATATTACCTA
B. infantis NCTC11817    CCCCTTTGTTCATTCGGCCGTCGCGCGCTTCCCCGGTCGCCCGTCGTGGTGCCACATATTGTTAGGCATGTTGACAAAATGCTGCGAAGAGGCATATATTACTGT
                         ************************** ************************* ************************************************

B. infantis ATCC15697    ATGTTCGTCTCACAGTTGTGATGGACGCTTATAGTGTTTTCATTCTGCAGAAAGGGAGAATGATGAGAAGAACCGCGATGAGGGT
B. infantis IN-F29       ATGTCCGTCTCACGGTTGTGATGGACGCTTATAGTGTTTTCATTCTGCAGAAAGGGAGAATGATGAGAAGAACCGCGATGAGGGT
B. infantis NCTC11817    ATGTTCGTCTCACAGTTGTGATGGACGCTTATAGTGTTTTCATTCTGCAGAAAGGGAGAATGATGAGAAGAACCGCGATGAGGGT
                         **** ******** **********************************************************************
```

***hmoA* (Blon_2347)**

```
B. infantis ATCC15697    CGTCGACGGAATCGGCGGTTTTCCGCTGTCTGGTGAGGGAGGGTGCTGAGGCATAGCGAAACGGTGCCGATATTTCATATATGTTAAGGACGTTGACAAAATATC
B. infantis IN-F29       CGTCGACGGAATCGGCGGTTTTCCGCTGTCTGGTGAGGGAGGGTGCTGAGGCATAGCGAAACGGTGCCGATATTTCATATATGTTAAGGACGTTGACAAAATATC
B. infantis Bi-26        CGTCGACGGAATCGGCGGTTTTCCGCTGTCTGGTGAGGGAGGGTGCTGAGGCATAGCGAAACGGTGCCGATATTTCATATATGTTAAGGACGTTGACAAAATATC
B. infantis R0033        CGTCGACGGAATCGGCGGTTTTCCGCTGTCTGGTGAGGGAGGGTGCTGAGGCATAGCGAAACGGTGCCGATATTTCATATATGTTAAGGACGTTGACAAAATATC
B. infantis BT1          CGTCGACGGAATCGGCGGTTTTCCGCTGTCTGGTGAGGGAGGGTGCTGAGGCATAGCGAAACGGTGCCGATATTTCATATATGTTAAGGACGTTGACAAAATATC
B. infantis NCTC11817    CGTCGACGGAATCGGCGGTTTTCCGCTGTCTGGTGAGGGAGGGTGCTGAGGCATAGCGAAACGGTGCCGATATTTCATATATGTTAAGGACGTTGACAAAATATC
                         *********************************************************************************************************

B. infantis ATCC15697    TCCCGAGACTTATCCTAGGTGCGTCCACCTCGCAGACGTGGCGGGCGCATCCAGGATCATAATTTCAAAGGAGAGACAATGAGAA
B. infantis IN-F29       TCCCGAGACTTATCCTGAATGCGTCCACCTCACAGACGTGGCGGGCGCATCCAGGATCATAATTTCAAAGGAGAGACAATGAGAA
B. infantis Bi-26        TCCCGAGACTTATCCTAGGTGCGTCCACCTCGCAGACGTGGCGGGCGCATCCAGGATCATAATTTCAAAGGAGAGACAATGAGAA
B. infantis R0033        TCCCGAGACTTATCCTAGGTGCGTCCACCTCGCAGACGTGGCGGGCGCATCCAGGATCATAATTTCAAAGGAGAGACAATGAGAA
B. infantis BT1          TCCCGAGACTTATCCTAGGTGCGTCCACCTCACAGACGTGGCGGGCGCATCCAGGATCATAATTTCAAAGGAGAGACAATGAGAA
B. infantis NCTC11817    TCCCGAGACTTATCCTAGGTGCGTCCACCTCGCAGACGTGGCGGGCGCATCCAGGATCATAATTTCAAAGGAGAGACAATGAGAA
                         ****************   ************ *****************************************************
```

***hmoA3* (Blon_2350)**

```
B. infantis ATCC15697    TGGCGCGACTCACATATATGTTAAGAATGTTGACGAACTCCATCTCCCGTGCCTATCATGAGTGCGTCCGTCCCGCAGATGCGGCGGGCGCGCTCAAATATCATC
B. infantis IN-F29       TGGCGCGACTCACATATATGTTAAGAATGTTGACGAACTCCATCTCCCGTGCCTATCATGAATGCGTCCGTCCCGCAGATGCGGCGGATACGCTCAAATATCATC
B. infantis Bi-26        TGGCGCGACTCACATATATGTTAAGAATGTTGACGAACTCCATCTCCCGTGCCTATCATGAGTGCGTCCGTCCCGCAGATGCGGCGGATGCGCTCAAATATCATC
B. infantis R0033        TGGCGCGACTCACATATATGTTAAGAATGTTGACGAACTCCATCTCCCGTGCCTATCATGAGTGCGTCCGTCCCGCAGATGCGGCGGATGCGCTCAAATATCATC
B. infantis BT1          TGGCGCGACTCACATATATGTTAAGAATGTTGACGAACTCCATCTCCCGTGCCTATCATGAATGCGTCCGTCCCGCAGATGCGGCGGATGCGCTCAAATATCATC
B. infantis NCTC11817    TGGCGCGACTCACATATATGTTAAGAATGTTGACGAACTCCATCTCCCGTGCCTATCATGAGTGCGTCCGTCCCGCAGATGCGGCGGGCGCGCTCAAATATCATC
                         ************************************************************* *************************   ***************

B. infantis ATCC15697    ATGTCAAAGGAGAGACGATGAGAAAACAAACCGTGCTGAAGGCGG
B. infantis IN-F29       ATGTCAAAGGAGAGACGATGAGAAGACAAACCGTGATGAAGGCGG
B. infantis Bi-26        ATGTCAAAGGAGAGACGATGAGGAGACAAACCGTGATGAAGGCGG
B. infantis R0033        ATGTCAAAGGAGAGACGATGAGGAGACAAACCGTGATGAAGGCGG
B. infantis BT1          ATGTCAAAGGAGAGACGATGAGGAGACAAACCGTGATGAAGGCGG
B. infantis NCTC11817    ATGTCAAAGGAGAGACGATGAGAAAACAAACCGTGCTGAAGGCGG
                         ********************** * ********** *********
```

***hmoA4* (Blon_2351)**

```
B. infantis ATCC15697    GAGCGGGTTTTGACATGCTGCGTTCTGTTTAGTGGATTACATGGTGGCGGGAATGACTATGTGTGGCGCGACTCACATATATGTTAAGAATGTTGACGAACTCCA
B. infantis BT1          GAGCGGGTTCTGAAGTGCTGCGTTCTATTTAGTGGATTACATGGTGGCGGGAGCGGCGATGTGTGGCGCGACTCACATATATGTTAAGAATGTTGACGAACTCCA
B. infantis NCTC11817    GAGCGGGTTTTGACATGCTGCGTTCTGTTTAGTGGATTACATGGTGGCGGGAATGACTATGTGTGGCGCGACTCACATATATGTTAAGAATGTTGACGAACTCCA
                         ********* ***  *********** *************************  * * ***********************************************

B. infantis ATCC15697    TCTCCCGTGCCTATCATGAATGCGTCCGTCCCGCAGATGCGGCGGATGCGCTCAAATATCATCATGTCAAAGGAGAGACGATGAG
B. infantis BT1          TCTCCCGTGCCTATCATGAATGCGTCCGTCTCGCAGATGCGGCGGGCGCGCTCAAATATCATCATGTCAAAGGAGAGACGATGAG
B. infantis NCTC11817    TCTCCCGTGCCTATCATGAATGCGTCCGTCCCGCAGATGCGGCGGATGCGCTCAAATATCATCATGTCAAAGGAGAGACGATGAG
                         ****************************** **************  **************************************
```

***hmoA5* (Blon_2352)**

```
B. infantis ATCC15697    GCGGCGGACGTGCTGCGTTCCATTTAACAGGTTACGTGGTGGCGGGAGCGGCGATGTGTGGCGCGACTCACATATATGTTAAGAATGTTGACGAACTCCATCTCC
B. infantis EK3          GCGGCGGACGTGCTGCGTTCCATTTAACAGGTTACGTGGTGGCGGCAGCGGCGATGTGTGGCGCGACTCACATATATGTTAAGAATGTTGACGAACTCCATCTCC
B. infantis Bi-26        GCGGCGGACGTGCTGCGTTCCATTTAACAGGTTACGTGGTGGCGGCAGCGGCGATGTGTGGCGCGACTCACATATATGTTAAGAATGTTGACGAACTCCATCTCC
B. infantis R0033        GCGGCGGACGTGCTGCGTTCCATTTAACAGGTTACGTGGTGGCGGCAGCGGCGATGTGTGGCGCGACTCACATATATGTTAAGAATGTTGACGAACTCCATCTCC
B. infantis NCTC11817    GCGGCGGACGTGCTGCGTTCCATTTAACAGGTTACGTGGTGGCGGGAGCGGCGATGTGTGGCGCGACTCACATATATGTTAAGAATGTTGACGAACTCCATCTCC
                         ********************************************* ***********************************************************

B. infantis ATCC15697    CGTGCCTATCATGAGTGCGTCCGTCCCGCAGATGCGGCGGATGCTCTCAAATATCATCATGTCAAAGGAGAGACGATGAGAAGAC
B. infantis EK3          CGTGCCTATCATGAGTGCGTCCGTCCCGCAGATGCGGCGGATACGCTCAAATATCATCATGTCAAAGGAGAGACGATGAGAAGAC
B. infantis Bi-26        CGTGCCTATCATGAGTGCGTCCGTCCCGCAGATGCGGCGGATACGCTCAAATATCATCATGTCAAAGGAGAGACGATGAGAAGAC
B. infantis R0033        CGTGCCTATCATGAGTGCGTCCGTCCCGCAGATGCGGCGGATACGCTCAAATATCATCATGTCAAAGGAGAGACGATGAGAAGAC
B. infantis NCTC11817    CGTGCCTATCATGAGTGCGTCCGTCCCGCAGATGCGGCGGATGCTCTCAAATATCATCATGTCAAAGGAGAGACGATGAGAAGAC
                         ****************************************** * ****************************************
```

***hmoA6* (Blon_2354)**

```
B. infantis ATCC15697    CGCGCGGCGGACGTGCTGCGTTCCATTTAGCCGGTTGCGTGGTGACGGGAGCGGCGGTGCGAGCGGGCCTGATGCAAATATGTTAAGGCTGTTGACAGTGCTGGC
B. infantis IN-F29       CGCGCGGCGGACGTGCTGCGTTCCATCTAGACCGTTGCAT--TGGCGGGAGCGGCGGTGCGAGCGGACTTGATGCAA-TATGTTAAGGCTGTTGACAGTGCTGGC
B. infantis EK3          CGCGCGGCGGACGTGCTGCGTTCCATTTAGCCGGTTGCGTGGTGACGGGAGCGGCGGTGCGAGCGGGCCTGATGCAAATATGTTAAGGCTGTTGACAATGCGGGC
B. infantis Bi-26        CGCGCGGCGGACGTGCTGCGTTCCATTTAGCCGGTTGCGTGGTGACGGGAGCGGCGGTGCGAGCGGGCCTGATGCAAATATGTTAAGGCTGTTGACAATGCGGGC
B. infantis R0033        CGCGCGGCGGACGTGCTGCGTTCCATTTAGCCGGTTGCGTGGTGACGGGAGCGGCGGTGCGAGCGGGCCTGATGCAAATATGTTAAGGCTGTTGACAATGCGGGC
B. infantis BT1          CGCGCGGCGGACGTGCTGCGTTCCATCTAGACCGTTGCAT--TGGCGGGAGCGGCGGTGCGAGCGGGCCTGATGCAAATATGTTAAGGCTGTTGACAGTGCTGGC
B. infantis NCTC11817    CGCGCGGCGGACGTGCTGCGTTCCATTTAGCCGGTTGCGTGGTGACGGGAGCGGCGGTGCGAGCGGGCCTGATGCAAATATGTTAAGGCTGTTGACAGTGCTGGC
                         ************************** *** * ***** *  ** ********************* * ******** ******************* *** ***

B. infantis ATCC15697    TCCCGTACCTATCATGGTTGCGTCCATCTCATAGGTGTGATGGATGCAACCGAAGTCATCATCGACTCAAAGGAGAGACAATGAG
B. infantis IN-F29       TCCCGTACTTATCATGGTTGCGTCCATCTCATAGGTGTGATGGACGCAACCGAAGTCATCATCGACTCAAAGGAGAGACAATGAG
B. infantis EK3          TCCCGTACTTATCATGGTTGCGTCCATCTCATAGGCGTGATGGACGCAACCGAAGTCATCATCGACTCAAAGGAGAGACAATGAG
B. infantis Bi-26        TCCCGTACTTATCATGGTTGCGTCCATCTCATAGGCGTGATGGACGCAACCGAAGTCATCATCGACTCAAAGGAGAGACAATGAG
B. infantis R0033        TCCCGTACTTATCATGGTTGCGTCCATCTCATAGGCGTGATGGACGCAACCGAAGTCATCATCGACTCAAAGGAGAGACAATGAG
B. infantis BT1          TCCCATACTTATCATGGTTGCGTCCATCTCATAGGTGTGATGGACGCAACCGAAGTCATCATCGACTCAAAGGAGAGACAATGAG
B. infantis NCTC11817    TCCCGTACCTATCATGGTTGCGTCCATCTCATAGGTGTGATGGATGCAACCGAAGTCATCATCGACTCAAAGGAGAGACAATGAG
                         **** *** ************************** ******** ****************************************
```
