## Supplementary material for "Human Milk Oligosaccharide Utilization in Intestinal Bifidobacteria is Governed by a Global Transcriptional Regulator NagR": Figure S2

**FIG S2**. Construction and metabolic profiling of the *B. infantis* ATCC 15697 *nagR*-KO mutant. (A) Schematic representation of insertional inactivation of *nagR*. The region used for a single crossover recombination event is in orange. The primers for genomic PCR (green) were designed to anneal outside the region used for recombination. (B) Results of genomic PCR. The amplicon sizes were expected to be 679 bp and 3164 bp for WT and *nagR*-KO strains, respectively. Four clones were analyzed. (C) HMO consumption and (D) organic acid production profiles of *B. infantis* ATCC 15697 WT and *nagR*-KO strains grown in MRS-CS-HMO. Data points represent the mean of three biological replicates. Error bars depict 95% confidence intervals for the mean. Timepoints where metabolite concentrations for WT and *nagR*-KO strains were significantly different (*, Padj<0.05) were identified using a linear regression. Bonferroni correction was used to adjust for multiple testing.

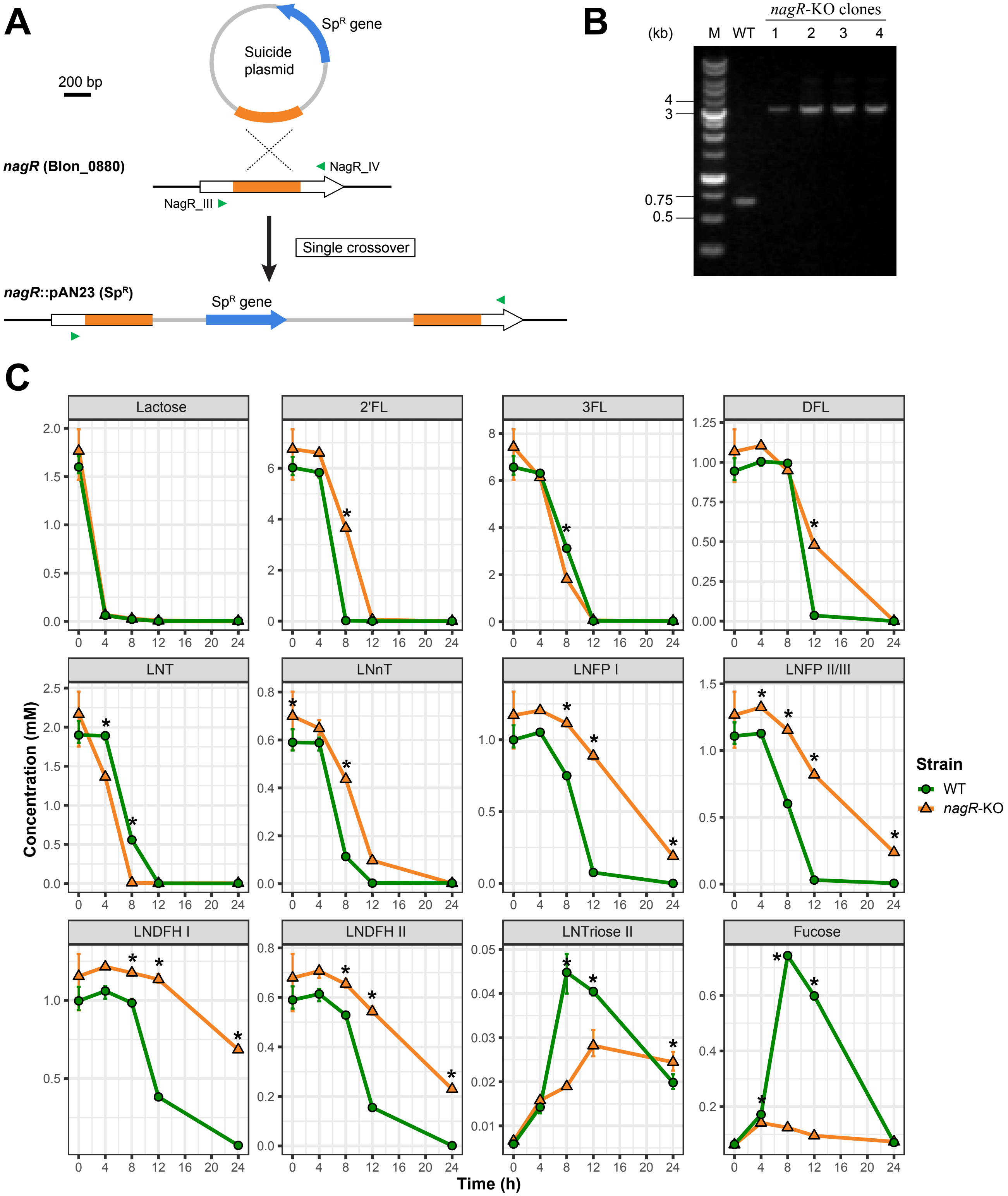

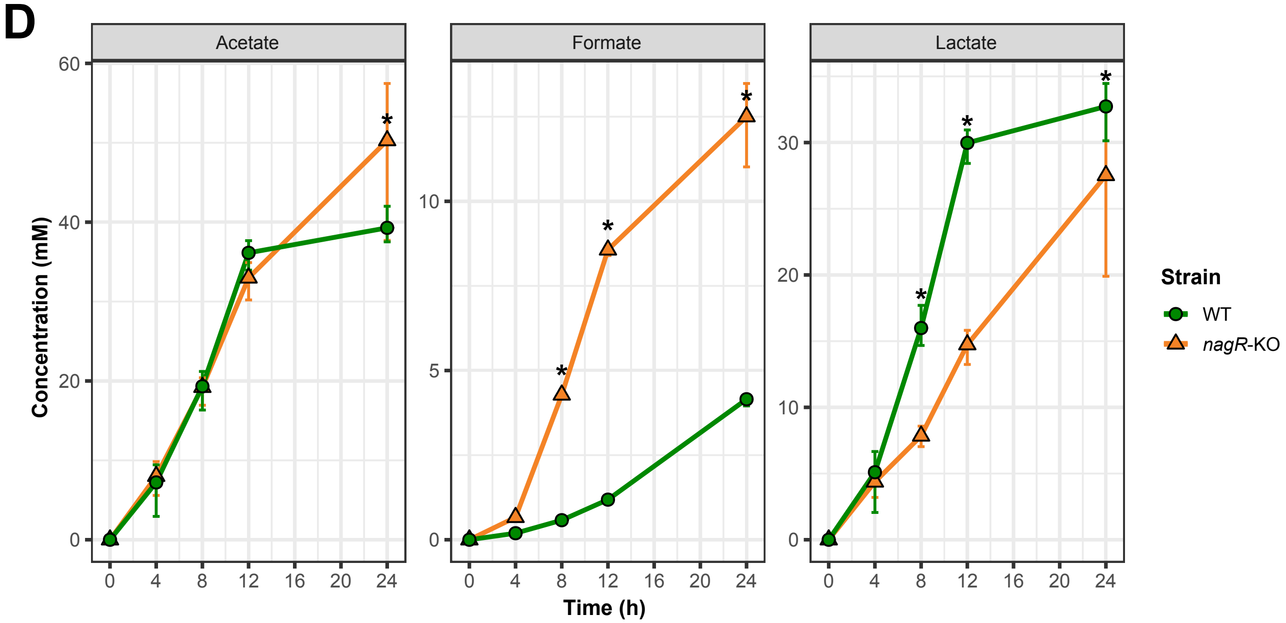
D
Acetate
Formate
Lactate
Concentration (mM)
Time (h)
Strain
WT
nagR-KO
0
4
8
12
16
20
24
60
40
20
10
5
30
