## Supplementary material for "Human Milk Oligosaccharide Utilization in Intestinal Bifidobacteria is Governed by a Global Transcriptional Regulator NagR": Figure S3

**FIG S3**. Heatmap depicting the expression of genes from Table S2A-B across four experimental conditions (WT and *nagR*-KO strains grown in MRS-CS supplemented with Lac or LNnT). Genes constituting the predicted NagR regulon are in bold.
Genomic clusters from Fig. 1B are marked by lines.

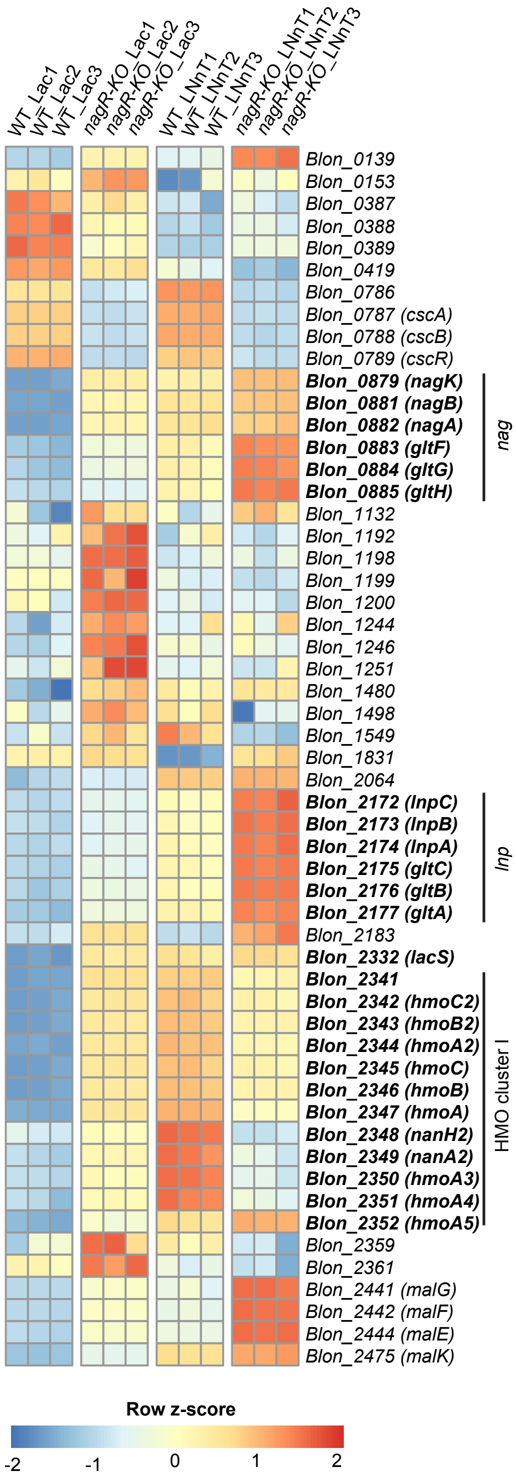
