## Supplementary material for "Human Milk Oligosaccharide Utilization in Intestinal Bifidobacteria is Governed by a Global Transcriptional Regulator NagR": Figure S4

**FIG S4**. (A) EMSA gels depicting titration of DNA probes (1 nM) containing predicted NagR operators with recombinant NagR.
(B) EMSA gels depicting titration of the NagR-DNA probe complex with various effector molecules. 25 nM of NagR and 1 nM of the *hmoA* probe were used in all reactions.

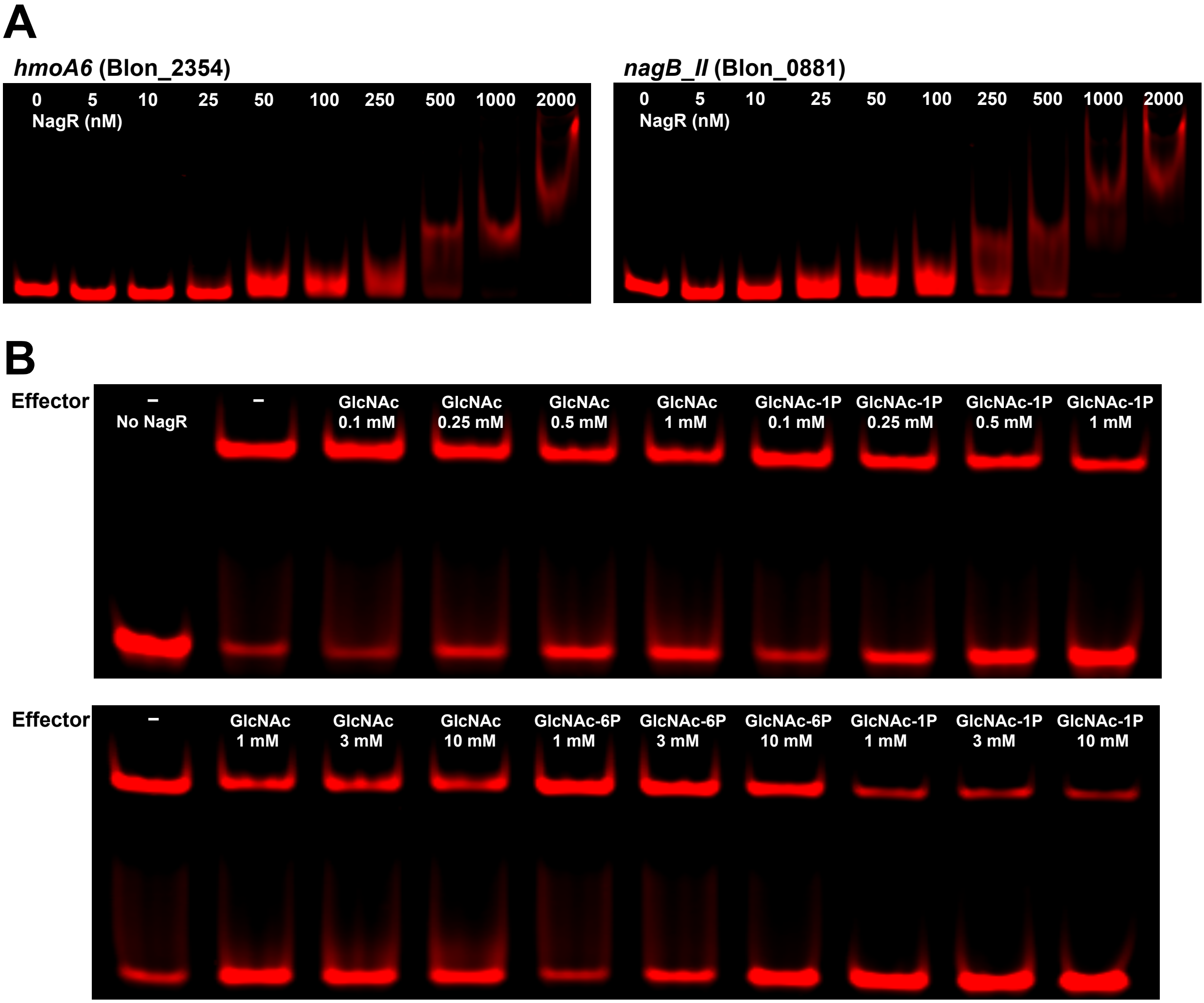
