## Supplementary material for "Human Milk Oligosaccharide Utilization in Intestinal Bifidobacteria is Governed by a Global Transcriptional Regulator NagR": Table S1

**Table S1.** Oligonucleotides used in this study. IRD oligonucleotides are 5'-labeled with the IRD700 dye, whereas RC oligonucleotides are unlabeled. Predicted NagR operators are in bold.

### Targeted gene disruption of the *nagR* gene

>NagR_I
CGGTACCCGGGGATCACTGCTGAGCATCGATACCGA
>NagR_II
CCAGCTCAAGGGATCCGATGGAAATATGGCCGATTTCG
>NagR_III
AGATGCTGGAGGAAGGATTGCTG
>NagR_IV
TGATGCTGCTCCACGTCTGC

### Cloning the of the *nagR* gene

>NagR_HisN_F
GATATAGGATCCATGCCGTATCTGGGTCTGAATAC
>NagR_HisN_R
GATATAGTCGACTTATTTCGTCAGGTGCTGAAAAACTG

### EMSA analysis

>nagK_IRD
TCGCCTTGGGTGAA**TTTGTTAAGATAGTTGTCAAT**AGTCGGATGCACAC
>nagK_RC
GTGTGCATCCGACT**ATTGACAACTATCTTAACAAA**TTCACCCAAGGCGA
>nagB_I_IRD
TCACAAGGTCCAAA**ATTGTTAGTATCATTAACAAA**TAGCGAATCGCCAA
>nagB_I_RC
TTGGCGATTCGCTA**TTTGTTAATGATACTAACAAT**TTTGGACCTTGTGA
>nagB_II_IRD
ATTTCGTCGTTGAA**ATTGTTATGAAACTTCACCAA**TAAATTCCAGTGTA
>nagB_II_RC
TACACTGGAATTTA**TTGGTGAAGTTTCATAACAAT**TTCAACGACGAAAT
>gltA_IRD
CGGGACGCGCGGTA**ATTGTTAGTTGGGTTGACAAT**AAAAGAGCAAAGGC
>gltA_RC
GCCTTTGCTCTTTT**ATTGTCAACCCAACTAACAAT**TACCGCGCGTCCCG
>hmoA2_IRD
TCGTGGTGCCACAT**ATTGTTAGGCATGTTGACAAA**ATGCTGCGAAGAGG
>hmoA2_RC
CCTCTTCGCAGCAT**TTTGTCAACATGCCTAACAAT**ATGTGGCACCACGA
>hmoA_IRD
GCCGATATTTCATA**TATGTTAAGGACGTTGACAAA**ATATCTCCCGAGAC
>hmoA_RC
GTCTCGGGAGATAT**TTTGTCAACGTCCTTAACATA**TATGAAATATCGGC

>hmoA3_IRD
GCGCGACTCACATA**TATGTTAAGAATGTTGACGAA**CTCCATCTCCCGTG
>hmoA3_RC
CACGGGAGATGGAG**TTCGTCAACATTCTTAACATA**TATGTGAGTCGCGC
>hmoA6_IRD
GGGCCTGATGCAAA**TATGTTAAGGCTGTTGACAGT**GCTGGCTCCCGTAC
>hmoA6_RC
GTACGGGAGCCAGC**ACTGTCAACAGCCTTAACATA**TTTGCATCAGGCCC
>ybaO_IRD
GGGAACCGCACAGAATAAATTGTCGTGATTTCACCTTTAAAATAAAATTAAAAGAGAAAAAAAT
TCTCTGTGGAAGG
>ybaO_RC
CCTTCCACAGAGAATTTTTTTCTCTTTTAATTTTATTTTAAAGGTGAAATCACGACAATTTATT
CTGTGCGGTTCCC
